# A method to map and interpret pleiotropic loci using summary statistics of multiple traits

**DOI:** 10.1101/2020.06.16.155879

**Authors:** Cue Hyunkyu Lee, Huwenbo Shi, Bogdan Pasaniuc, Eleazar Eskin, Buhm Han

**Affiliations:** Department of Biomedical Sciences, Seoul National University College of Medicine, Seoul, Republic of Korea; Department of Convergence Medicine, University of Ulsan College of Medicine, Asan Medical Center, Seoul, Republic of Korea; Bioinformatics Interdepartmental Program, University of California, Los Angeles, Los Angeles, California, USA; Department of Human genetics, University of California, Los Angeles, Los Angeles, California, USA; Department of Pathology and Laboratory Medicine, University of California, Los Angeles, Los Angeles, California, USA; Department of Computer Science, University of California, Los Angeles, Los Angeles, California, USA; Department of Computational Medicine, University of California, Los Angeles, Los Angeles, California, USA; Interdisciplinary Program in Bioengineering, Seoul National University, Seoul, Republic of Korea

## Abstract

The identification of pleiotropic loci and the interpretation of the associations at these loci are essential to understand the shared etiology of related traits. A common approach to map pleiotropic loci is to use an existing meta-analysis method to combine summary statistics of multiple traits. This strategy does not take into account the complex genetic architectures of traits such as genetic correlations and heritabilities. Furthermore, the interpretation is challenging because phenotypes often have different characteristics and units. We propose PLEIO, a summary-statistic-based framework to map and interpret pleiotropic loci in a joint analysis of multiple traits. Our method maximizes power by systematically accounting for the genetic correlations and heritabilities of the traits in the association test. Any set of related phenotypes, binary or quantitative traits with differing units, can be combined seamlessly. In addition, our framework offers interpretation and visualization tools to help downstream analyses. Using our method, we combined 18 traits related to cardiovascular disease and identified 20 novel pleiotropic loci, which showed five different patterns of associations. Our method is available at https://github.com/hanlab-SNU/PLEIO.

## 2 Introduction

Recent genome-wide association studies (GWAS) have shown that some genetic variants are associated with multiple traits, a phenomenon called pleiotropy^1,2^. The identification of pleiotropic loci is important to understand the shared etiology of the related traits and to find common drug targets. To identify pleiotropic loci, several studies proposed multi-trait analyses that combine summary statistics of multiple traits into one^3-5^. Due to the similarity of this task with meta-analysis, studies have employed meta-analysis methods^6-9^. Another category of multi-trait analyses is to maximize power for a single trait by incorporating other traits as prior information^10-13^. In this study, we focused on the former, the joint analysis of multiple traits.

Applying an existing meta-analysis method to multi-trait analyses is not optimal for several reasons. First, the meta-analysis methods completely ignore the genetic architectures of the traits. The magnitude and direction of the genetic correlations suggest the expected pattern of the genetic effects for multiple traits. The heritabilities suggest the expected strengths of the genetic effects for multiple traits. Therefore, a lot of information resides in the genetic architecture, which can help map pleiotropic loci. Second, the meta-analysis methods depend on the scales and units of the phenotypes. The units often differ among quantitative traits, and the effect size definitions differ between binary and continuous traits. Most meta-analysis methods directly use the raw numbers of effect sizes as input, so they are not optimal for analyzing heterogeneous traits. For the same reason, interpretation tools such as the forest plot^14^ or m-value^15^ are less useful. Third, environmental correlations exist among traits collected from the same individuals. Without systematically correcting for environmental correlations, a naïve application of meta-analysis methods can inflate false positives.

Here, we propose an optimized multi-trait method to map and interpret pleiotropic loci called PLEIO (Pleiotropic Locus Exploration and Interpretation using Optimal test). As with meta-analysis methods, our method uses only summary statistics. Our method starts by estimating the genetic correlations, environmental correlations, and heritabilities from the whole-genome summary statistics. We then standardize the effect sizes of heterogeneous traits, converting the effect sizes of binary traits to effect sizes for liabilities. An advantage of the standardization is that the analysis and interpretation become independent of the phenotypic units. Another advantage is that heritabilities can suggest the expected strengths of the effect sizes of a pleiotropic locus under the standardized scale. We have developed an optimized association test that takes into account both the genetic correlations and heritabilities to maximize power. Our test is a variance component test to check non-zero variance of the random genetic effect, where we model the genetic effect to follow the genetic covariance. Our test correctly controls the false positive rate by accounting for the environmental correlations. To increase efficiency in finding the maximum likelihood estimate, we developed an optimization technique using the spectral decomposition of the variance. Even with this technique, obtaining the p-value is computationally challenging because the small number of traits induce small sample problem. We overcome this challenge by implementing an importance sampling method that shortens the computation time for combining 100 traits to 1 day.

Real data analyses and simulations show that PLEIO is powerful for identifying pleiotropic loci. Unlike other methods that performed well under specific situations, PLEIO was consistently powerful in all scenarios because it learned the genetic architecture from data and optimized itself to that situation. In addition to the powerful association test, PLEIO offers tools for the interpretation and visualization of the pleiotropic loci. We used PLEIO to combine 18 traits related to cardiovascular disease and identified 20 novel pleiotropic loci. These loci were categorized into five groups based on their association patterns, which may represent distinct pathways. PLEIO is freely available at https://github.com/hanlab-SNU/PLEIO.

## 3 Results

### 3.1 Overview of Method

PLEIO is a multi-trait framework to map and interpret pleiotropic loci (**Figure 1**). Assume a toy example that combines three traits (A, B, and C) (**Supplementary Figure 1**). At SNP *X*_1_, we observed the effect sizes (betas) of (2.2, 2.8, -1.2), and at another SNP *X*_2_, we observed the effect sizes of (−1.5, 0.4, -2.7). We assume that the variances of all estimates were identical. Then, if we apply the fixed effects meta-analysis (inverse-variance method), we get the same p-value for both SNPs (*P*=0.03) because the average beta is the same. However, suppose we know that traits A and B have a positive genetic correlation, and trait C has a negative genetic correlation with the rest. Then, SNP *X*_1_ is more likely to be a true signal than SNP *X*_2_ because the effect directions conform to the genetic correlations. Moreover, suppose we know that trait B has the greatest heritability and trait C has the smallest heritability. Then, the association at SNP *X*_1_ is even more likely because the relative strengths of the effect size conform to the heritabilities. Our method accounts for both the genetic correlations and heritabilities and gives a more significant p-value at SNP *X*_1_ (P= 0.0006) than SNP *X*_2_ (P=0.1).

**Figure 1.**
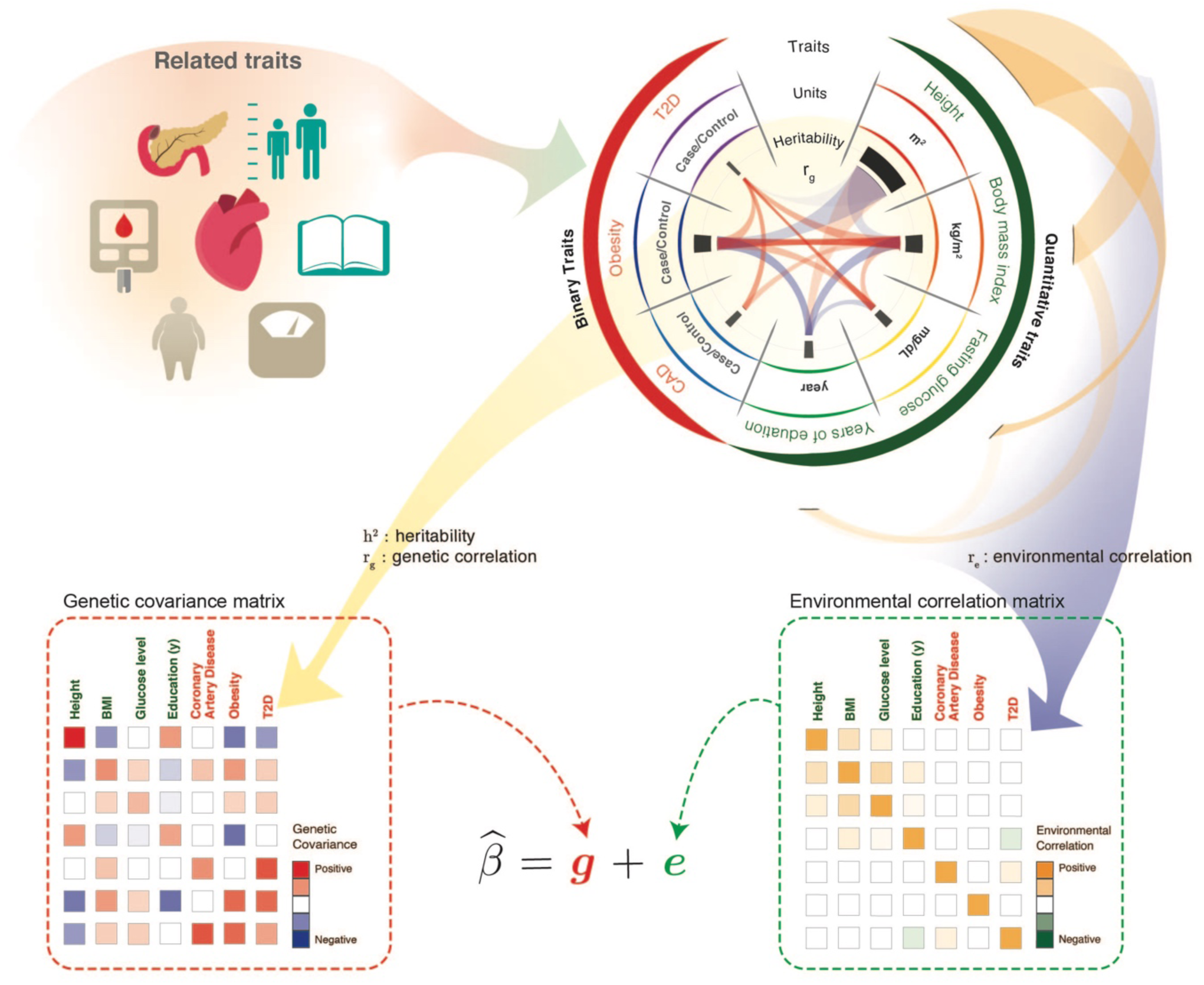
Overview of the PLEIO framework.

PLEIO consists of five steps. First, we apply the linkage disequilibrium (LD) score regression to the genome-wide summary data of traits to obtain the genetic correlations **C**^**2**^and the heritabilities **h**^**2**^. We summarize **C**_**g**_ and **h**^**2**^ into the genetic covariance **Ω**. It is not obvious how to estimate the environmental correlation **C**_**e**_, so we propose a strategy for that. Second, we transform the effect sizes 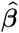 into the standardized effect sizes 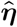, converting the effect sizes of binary traits to the effect sizes for liabilities. Third, we apply our variance component test to map pleiotropic loci. We assume 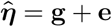 where **g** is the genetic effect and **e** is the error. Our main assumption is that the genetic effects follow the genetic covariance, Var(**g**) = **τ**^2^**Ω**. We then test the hypothesis **τ**^2^ > 0 versus **τ**^2^ = 0. To find the maximum likelihood estimate (MLE) of **τ**^2^ efficiently, we utilize an optimization technique using spectral decomposition of the variance. Fourth, we apply an importance sampling method to assess the p-value. Fifth, we report and visualize the results to help interpretation.

### 3.2 Evaluation of False positive rates

We evaluated the false positive rate (FPR) of PLEIO using simulations. We assumed the null hypothesis of no genetic effect at a SNP for *T* traits. There can be different situations under the null hypothesis. For example, the *T* statistics can be independent or correlated due to environmental correlations (**C** _**e**_), which reflect sample overlap, or the input parameter information for PLEIO, namely the heritability and genetic correlation estimates, can differ. Overall, we varied four factors: (1) the number of traits (*T*), (2) the environmental correlation matrix (**C**_**e**_), (3) the heritability parameter for PLEIO (**h**^**2**^), and (4) the genetic correlation parameter for PLEIO (**C**_**g**_).

Specifically, we adopted three different numbers of traits (*T*= 5, 10, and 20). We assumed that the sample overlap may or may not exist and set the non-diagonals of **C** _**e**_ to 0.5 in the former and 0 in the latter. We assumed two different patterns for the input parameter **h**^2^. In the “equal **h**^**2**^” situation, we assumed the same heritability input (h^2^ = 0.5) for all traits. In the “different **h**^**2**^” situation, we assumed a linearly increasing heritabilities from h^2^ = 0.1 to 0.5. We assumed two different patterns for the input parameter **C**_**g**_. In the “uniform **C**_**g**_” situation, the non-diagonals of **C**_**g**_ were all set to 0.3. In the “partitioned **C**_**g**_” situation, we assumed two subgroups and set non-diagonals to 0.3 within a group and 0 between groups. Thus, we tested 24 different situations (3 × 2 × 2 × 2). We generated one million null datasets per each situation and calculated FPR at α = 0.05. **Supplementary Table 1** shows that FPR of PLEIO is well calibrated in all situations.

Next, we examined FPR at a lower threshold. We increased the number of null datasets to a billion to measure FPR at the conventional GWAS threshold (5 × 10^−8^). We tested three numbers of traits (*T* = 5, 10, and 20) while assuming the equal **h**^**2**^, partitioned **C**_**g**_, and no sample overlap. **Supplementary Table 2** shows that PLEIO’s FPR is well calibrated for α down to 5 × 10^−8^. See **Supplementary Note** for a detailed explanation for the observed effect size generation.

### 3.3 Power simulations

We compared the power of PLEIO against three meta-analysis approaches: the Lin-Sullivan method (LS)^16^, RE2C^6^, and ASSET^9^. LS is a generalization of the fixed effects model, and RE2C is a generalization of the Han-Eskin random effects model^15^. ASSET is a subset-based method assuming that the true effects could only exist in a subset of the studies. All these methods can take into account the environmental correlations due to sample overlap. We confirmed that FPRs were well calibrated with each method (**Supplementary Table 3**).

We assessed the power of the methods in various simulation settings. Each setting defined a specific genetic correlation structure **C**_**g**_, heritabilities **h**^**2**^, phenotypic units (**U**), and the types of traits (quantitative (**Q**) or binary (**B**)). In each setting, we assumed *T* = 7 traits and repeated simulations 10,000 times. See **Supplementary Note** for a detailed explanation for the observed effect size generation.

First, we assumed a fixed heritability and very high correlations (*r*^2^ = 0.99) among the 7 traits. This represents the situation in which the same traits were collected in multiple studies. In this situation, all methods performed similarly well except ASSET (**Figure 2A**). With a sample size of *N* = 50,000, the powers of PLEIO, LS, and RE2C were 64.9%, 64.8%, and 63.9%. It was natural that LS performed well because LS is optimized for the fixed-effect situation. PLEIO also performed similarly, because PLEIO learns the high genetic correlations from data and adjusts itself to the situation.

**Figure 2.**
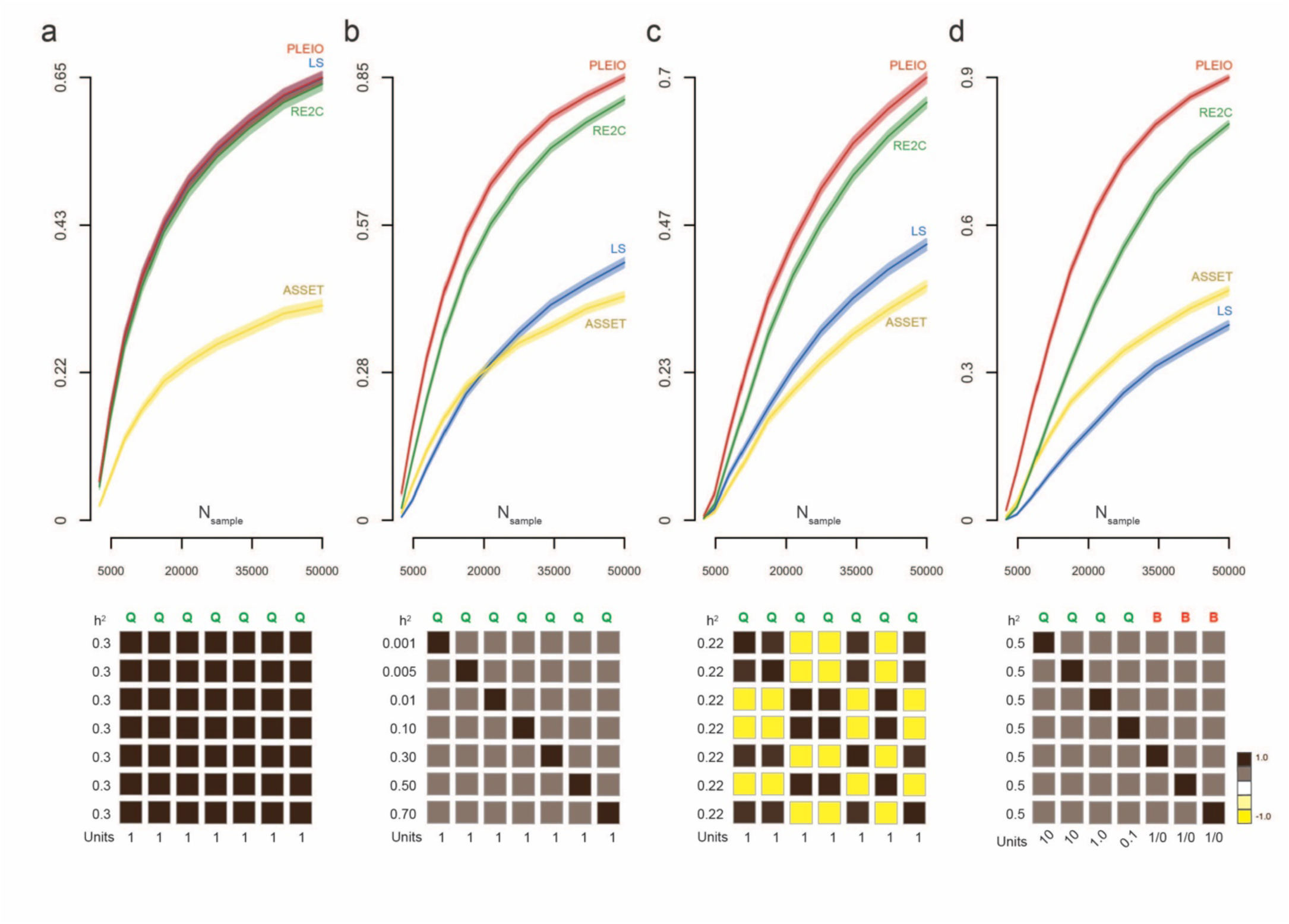
The results of the power test. Line plots show the power of PLEIO (red), RE2C (green), ASSET (yellow), and LS (blue), with the simulation settings shown at the bottom. The letters Q and B indicate the type of a phenotype ([Q]uantitative or [B]inary). ‘h^2^’ denotes the heritabilities of the traits; ‘Units’ denotes the phenotypic units, and the matrix shows the genetic correlation structure among traits. (a) We assumed a homogeneous situation that the same traits were studied multiple times, which is the assumption of the fixed effects meta-analysis. (b) We varied the heritabilities of the traits from 0.001 to 0.7. (c) We assumed that there were two subgroups. The correlation within the first group (three traits) was set to 0.95, and the correlation within the second group (four traits) was set to 0.9. The correlation between the two groups was set to -0.9. (d) We assumed a mixture of binary and quantitative traits and varied the phenotypic units among the quantitative traits from 0.1 to 10.

Second, we assumed different heritabilities for 7 traits, varying from 0.001 to 0.7. We assumed a uniform genetic correlation *r* = 0.5 between all trait pairs. In this situation, PLEIO outperformed the others (**Figure 2B**). With a sample size of *N* = 50,000, PLEIO achieved a power of 84.9%, while the second best method (RE2C) achieved 80.6% and the third best method (LS) achieved 49.4%. PLEIO achieved high power because PLEIO can take into account different heritabilities of the traits.

Third, we assumed a block structured correlation pattern. We divided 7 traits into two groups (3 traits and 4 traits). We set the correlations in the first group to 0.95 and the correlations in the second group to 0.9. We set the correlations between the groups to a negative value of -0.9. We assumed a uniform heritability of 0.26 for all traits. PLEIO showed the highest power among all methods (**Figure 2C**). With a sample size of *N* = 50,000, PLEIO achieved a power of 76.3%, while the second best method (RE2C) achieved 72.7% and the third best method (LS) achieved 48.3%. PLEIO achieved high power because PLEIO can take into account the genetic correlation structure of the traits.

Fourth, we assumed a mixture of quantitative and qualitative traits. We assumed 4 quantitative traits and 3 binary traits. For quantitative traits, we assumed different phenotypic units ranging from 0.1*U* to 10*U*. We assumed a fixed heritability and a uniform genetic correlation for all traits. Again, PLEIO achieved the highest power. With a sample size of *N* = 50,000, PLEIO achieved a power of 89.6%, while the second best method (RE2C) achieved 80.2% and the third best method (ASSET) achieved 46.5%. PLEIO achieved high power because PLEIO can systematically combine heterogeneous traits by standardizing the effect sizes.

So far, we varied only one factor in each simulation: different heritabilities, a complex pattern of genetic correlations, and different phenotypic units. In reality, all three can occur together. We simulated such a combined situation and found that the power gain was the greatest. PLEIO achieved a power of 71.0%, while the power of the second best method (RE2C) was only 56.6% (**Supplementary Figure 2**). The power increased by a quarter, which can be interpreted to mean that in this specific situation 50 instead of 40 loci can be found with our method.

### 3.4 Joint analysis of multiple traits related to Cardiovascular disease

We applied PLEIO to identify pleiotropic loci associated with traits related to cardiovascular disease (CVD). To this end, we collected summary statistics of 18 traits from multiple consortia (**Supplementary Table 4)**. We selected 12 binary traits from the Neale lab’s UK Biobank GWAS results (http://www.nealelab.is/uk-biobank, **Supplementary Table 5**) by searching with the terms: heart, hypertension, obesity, lipoproteins, cholesterol, and diabetes. We collected 4 lipid traits from the Global Lipid consortium^17^, 1 binary trait (coronary artery disease) from the CARDIoGRAM+C4D consortium^18^, and 1 trait (fasting glucose) from the MAGIC (**M**eta-**A**nalysis of **G**lucose and **I**nsulin-related traits **C**onsortium)^19^. In total, we collected 13 binary and 5 quantitative traits. See **Online Methods** for details of the trait selection. Quantitative traits had differing units. Lipid traits had the unit of mg/dl, whereas the fasting glucose had the unit of mmol/l^17,19^. We used 1,777,412 imputed SNPs overlapping among all datasets. These traits showed differing heritabilities and non-zero genetic and environmental correlations (**Supplementary Figure 3**).

PLEIO identified 618 GWAS top hits (**Figure 3** and **Supplementary Table 6**). Among those, we found 20 independent novel variants. These had no known associations to CVD traits and were not significant in each single study (**Supplementary Table 7**). The local Manhattan plots of these loci are shown in **Supplementary Figure 4.** We used the Variant Effect Predictor (VEP v.97.2) in ENSEMBL GRCh37 and obtained the annotations of these variants. The 20 variants included 10 intronic variants, 4 intergenic variants, 2 downstream gene variants, 1 upstream gene variant, 1 5-prime UTR variant, 1 missense variant, and 1 non-coding transcript exonal variant. The 618 top hits included 368 intronic variants, 113 intergenic variants, 37 upstream gene variants, 27 downstream variants, 23 3-prime UTR variants, 21 missense variants, 12 non-coding transcript exon variants, 11 synonymous variants, and 6 5-prime UTR variants. The detailed annotations are in **Supplementary tables 8 and 9. Figure 3A** shows a circular plot whose radial position indicates the genomic position, and the heights of the points are the statistical significances of the variants. Using the 618 top hits, we performed an additional analysis with DAVID v.6.8. Given the list of genes obtained by VEP, we used DAVID to search for the presence of known trait-gene associations based on the Genetic Association Database (GAD, **Supplementary table 10**). We curated the reported trait-gene associations into 8 categories: Coronary artery disease, Fasting glucose, Hypertension, Diabetes, High density lipoprotein, Low density lipoprotein, Total Cholesterol, and Total glycerides. That is, we categorized the variants into 8 groups based on the trait category of the known association. We visualized the results in the inner circle of **Figure 3A**, where each ribbon indicates a pair of genes in the same phenotypic category. **Figure 3B** shows the Manhattan plot of the PLEIO association results. We also compared the results of PLEIO to the original summary statistics using a mirrored Manhattan plot (**Supplementary Figure 5**). For a detailed description of this analysis, see **Online Methods**.

**Figure 3.**
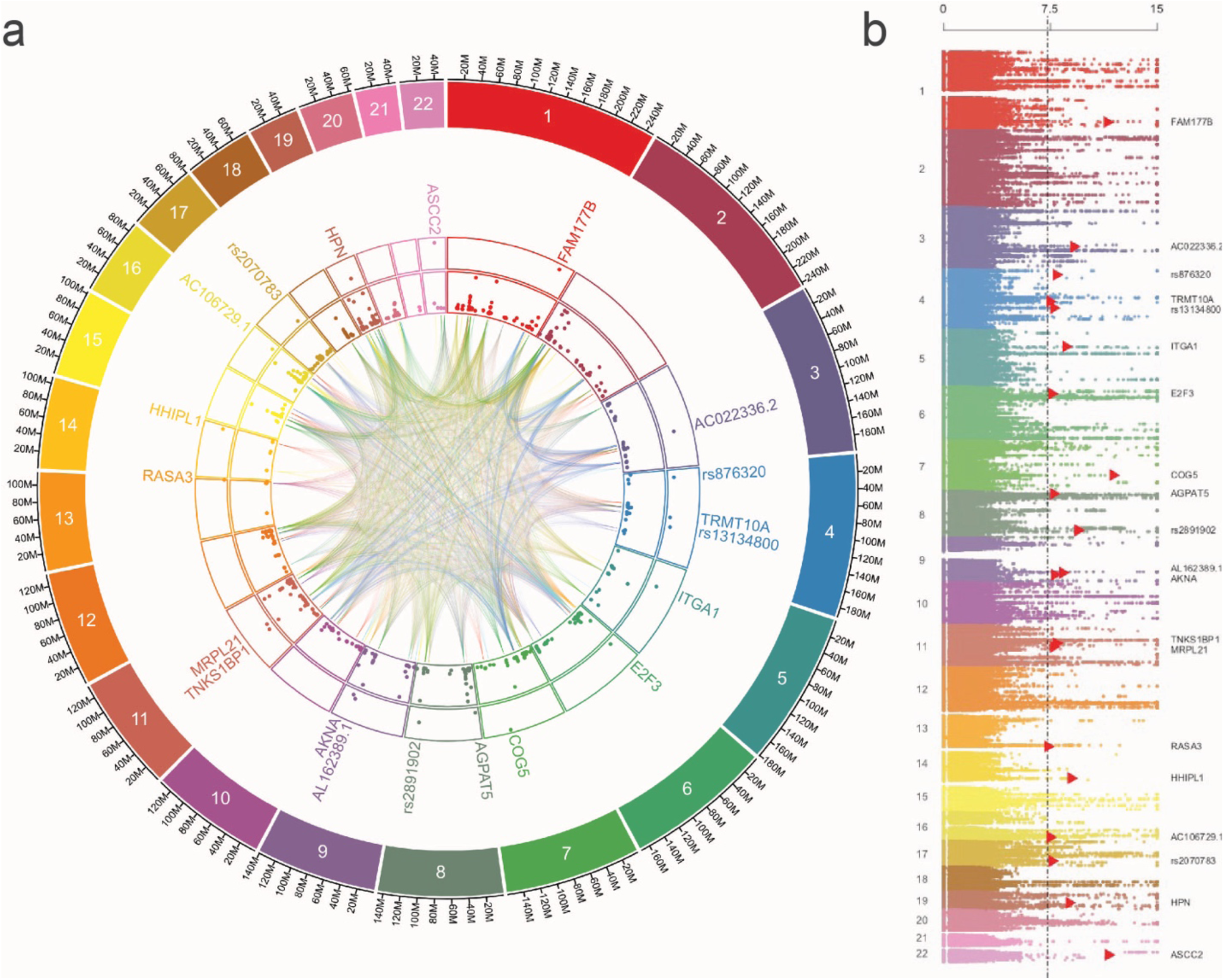
The summary of the real data-analysis. (a) The circular plot shows the locations and the statistical significances of the 20 novel variants (outer edge) and the 618 GWAS top SNPs (inner edge). The inner ribbons connect the variants in the same functional category found by the DAVID analysis. (b) The Manhattan plot of the PLEIO association results. Red triangles indicate the 20 novel loci.

### 3.5 Interpretation of the PLEIO analysis results

We visualized the multi-trait associations of each locus using a circular plot, which we call *pleiotropy plot*. The pleiotropy plot includes the local Manhattan plot and the bar plot of the standardized effect sizes. The inner ribbons show the genetic correlations as colors and the explained heritabilities by the locus as widths. We drew pleiotropy plots of the 20 novel variants we identified (**Figure 4** and **Supplementary Figure 6**). Based on the patterns observed in these plots, we categorized the 20 variants into five groups, which may imply distinct underlying pathways (**Figure 5**).

**Figure 4.**
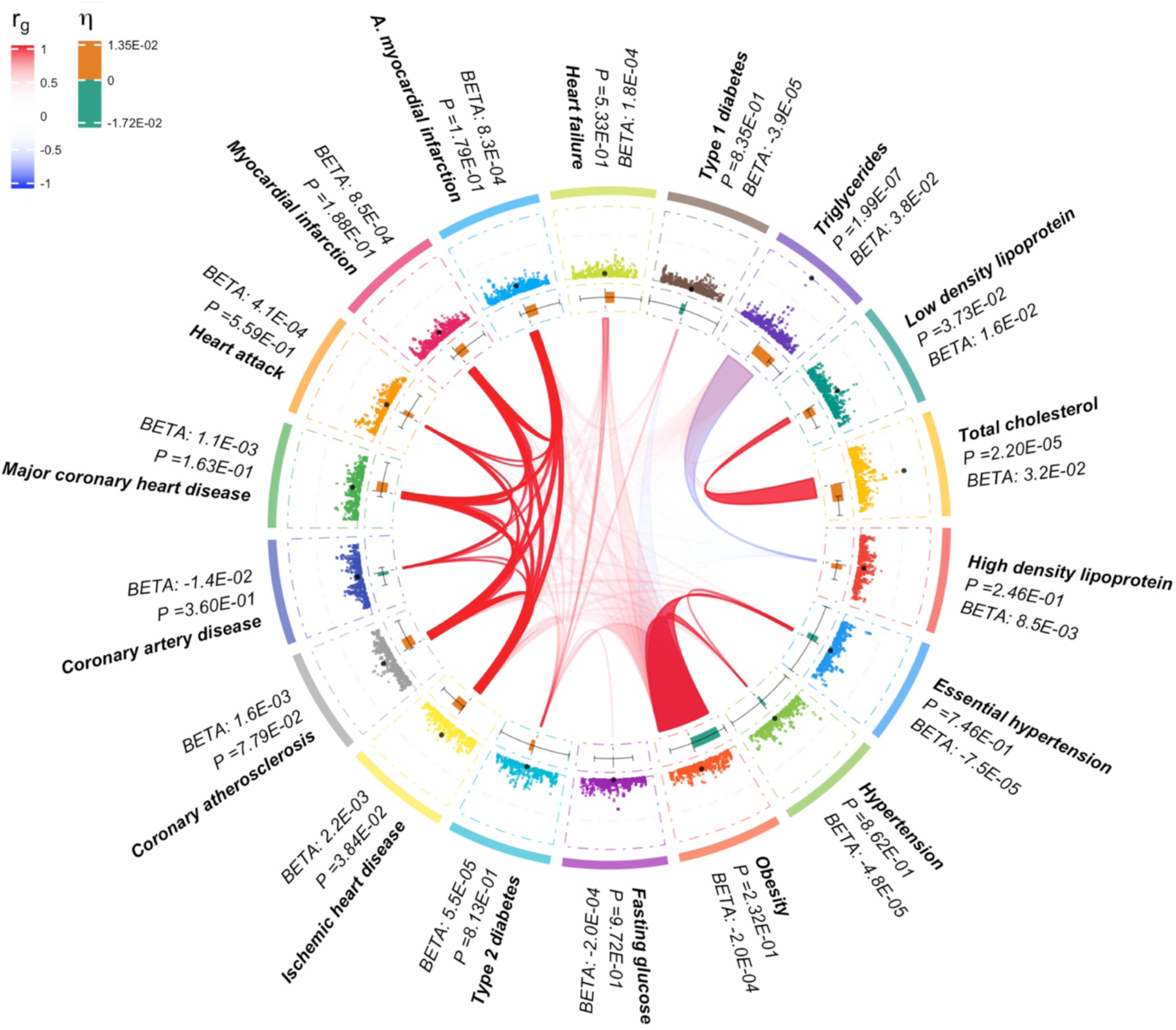
Pleiotropy plot of rs1688030, an intronic variant of the *HPN* gene. The outer edge is the local Manhattan plots for each trait within 1 Mb window. The horizontal bar plot shows the direction and size of the standardized effect size (*η*) with 95% confidence interval for each trait. The inner ribbons show the genetic correlations (as the color: positive *r*_*g*_ as red and negative *r*_*g*_ as blue) and the explained heritability by the locus (as the width of the ribbon end).

**Figure 5.**
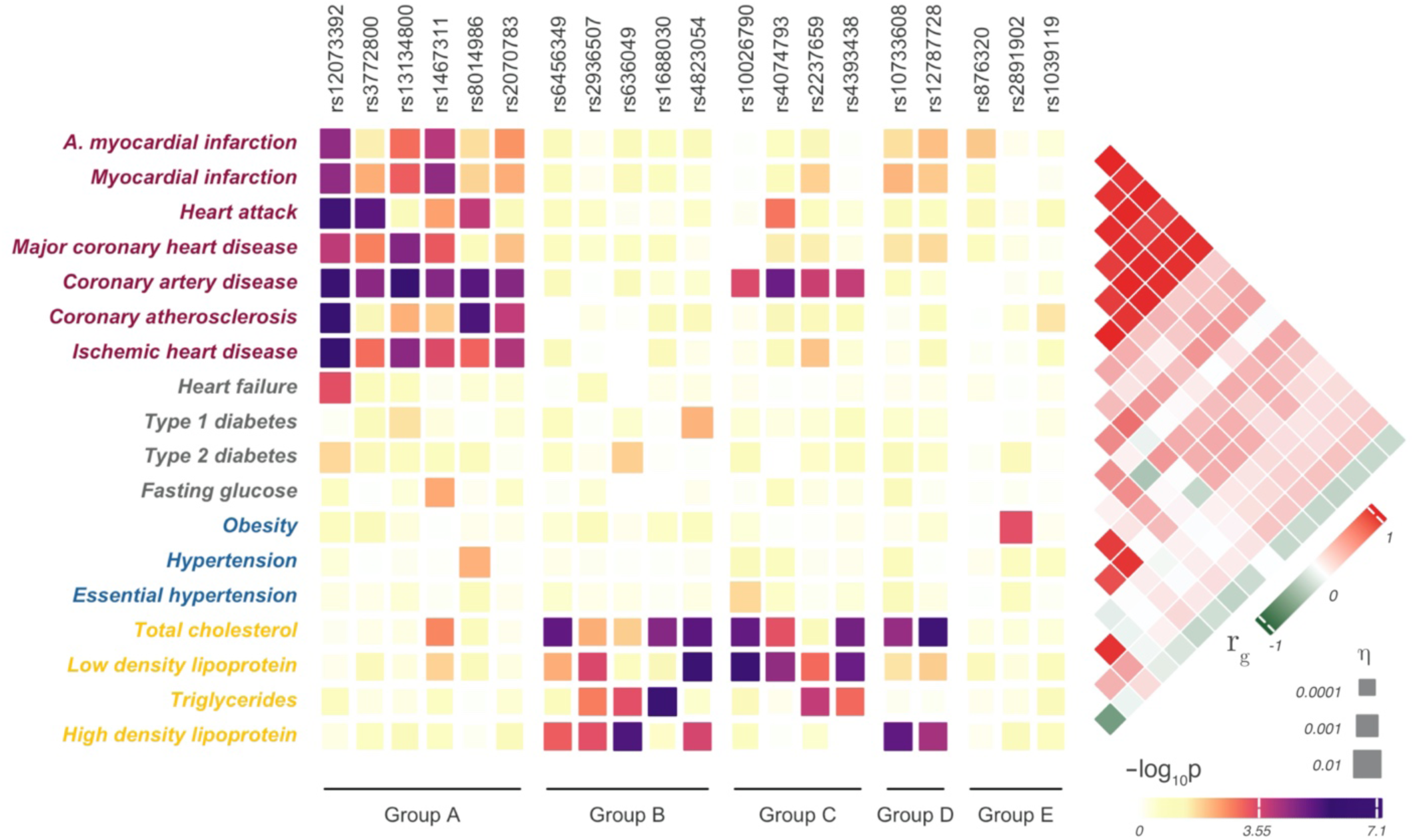
Distinct association patterns of 20 novel variants identified by PLEIO. Each box represents the association of a variant with a trait, where the size of the box indicates the magnitude of the standardized effect size (*η*) and the color of the box indicates the statistical significance. The right-side heatmap shows the genetic correlations. We divided the variants into five groups based on their association patterns.

The first group had associations driven by seven binary traits: 6 traits from the UK Biobank (acute myocardial infarction, myocardial infarction, heart attack, major coronary heart disease, coronary atherosclerosis, and ischemic heart disease) and one trait (coronary artery disease) from CARDIoGRAM+C4D. These seven traits showed high genetic correlations (**Figure 5**). The variants showing this pattern were rs12073392 in the *FAM1777B* gene (1q41), rs3772800 near the *AC022336.2* gene (3q21.2), rs13134800 near the *LINC02502* gene (4q27), rs1467311 near the *AL162389.1* gene (9q31.2), rs8014986 in the *HHIPL1* gene (14q32.2), and rs2070783 near the *PECAM1* gene (17q23.3).

The second group had associations driven by lipid phenotypes (triglycerides, low density lipoprotein; LDL, high density lipoprotein; HDL, and total cholesterol). The variants showing this pattern were rs6456349 in the *E2F3* gene (6p22.3), rs2936507 in the *AGPAT5* gene (8p23.1), rs636049 in the *MRPL21* gene (11q13.3), rs1688030 in the *HPN* gene (19q13.11), and rs4823054 in the *ASCC2* gene (22q12.2). These variants showed differing associations to the lipid phenotypes. rs6456349 showed the strongest associations to the total cholesterol and HDL. rs636049 showed the strongest associations to HDL and triglycerides. rs1688030 showed the strongest associations to the triglycerides and the total cholesterol. rs4823054 showed the strongest association to LDL and total cholesterol. rs2936507 showed similar strengths of associations to the triglycerides, LDL, HDL, and total cholesterols.

The third group had associations driven by both the coronary artery disease and the lipid phenotypes. The variants showing this pattern were rs10026790 in the *TRMT10A* gene (4q23), rs4074793 in the *ITGA1* gene (5q11.2), rs2237659 in the *COG5* gene (7q22.3), and rs4393438 in the *RASA3* gene (13q34). All four SNPs showed an association (*P* < 0.0001) to LDL.

The fourth group had associations with lipid phenotypes and a moderate association with myocardial infraction. The variants showing this pattern were rs10733608 in the *AKNA* gene (9q32) and rs12787728 in the *TNKS1BP1* gene (11q12.1). Both variants showed associations with HDL and total cholesterol (*P* < 0.0001) but not with LDL and triglycerides.

The fifth group was the variants that were not categorized to the four aforementioned groups. The variants in this group were rs876320 near the *FGFBP1* gene (4p15.32), rs2891902 near the *RPL35AP19* gene (8q24.12), and rs1039119 in the *AC106729.1* gene (16q23.1). rs2891902 showed the strongest association to the obesity (*P* < 0.001) and weak associations to the type 2 diabetes and hypertensions. rs1039119 and rs876320 were interesting because their associations to all traits were weak (*P* > 0.01). The strongest associations of rs1039119 were to coronary atherosclerosis (*P* = 0.02) and triglycerides (*P* = 0.08). However, this SNP’s effect size directions to the seven binary traits in the first group were all concordant to the genetic correlations of these traits. The strongest associations of rs876320 were to acute myocardial infarction (*P* = 0.01), myocardial infarction (*P* = 0.04), and heart attack(*P* = 0.04). This SNP’s effect size directions to these three traits were all concordant to the genetic correlations. Thus, PLEIO seems to have captured the aggregate information in multiple weak associations by considering the fact that the effect size directions were concordant to the genetic correlations.

## 4 Discussion

We proposed PLEIO, a framework to identify and interpret pleiotropic loci using summary statistics of multiple traits. PLEIO increased the statistical power using two strategies. First, we modeled the genetic correlations and heritabilities in our variance component test. Second, we took into account the differences in effect size scales and units among traits. Our method offers interpretation and visualization tools to help understand shared association patterns of pleiotropic loci.

To increase the efficiency, we applied two techniques: the maximum likelihood estimation using the spectral decomposition of the variance, and the importance sampling method. Using these approaches, PLEIO can combine 100 traits with a million SNPs in one day using a single CPU. Thus, any researchers with minimal computing power could utilize our method with ease.

Our method is general and includes other previous meta-analysis methods as special cases. If we set the genetic covariance matrix to a matrix of ones and the environmental correlations to zero, the test is approximately equivalent to the fixed effects meta-analysis method. If we assume environmental correlations, the test is approximately equivalent to the Lin-Sullivan method^16^. If we set the genetic covariance matrix to an identity matrix and the environmental correlations to zero, the resulting test is similar to the heterogeneity test in the Han-Eskin random effects model^15^. If we set the genetic covariance matrix to an identity matrix and assume environmental correlations, the resulting test is similar to the heterogeneity test in the RE2C framework^6^. A difference of PLEIO is that, unlike other methods optimized for specific situations, it learns the genetic covariance and the environmental correlations from data and adjusts itself to that situation. For example, if we have a collection of the studies for the same trait, PLEIO will learn this information and act as if it were a fixed effects meta-analysis method.

There can be various sources for the genetic correlations. When we combine the same traits, the genetic correlations will be nearly perfect. When we combine different traits, we will observe imperfect genetic correlations or even negative genetic correlations. Another situation is that we combine the same traits from multiple ethnicities. In this situation, the genetic correlation is usually imperfect and positive (0 < *r*_*g*_ < 1). Recent methods can estimate genetic correlations across different populations by accounting for the ethnic differences of LD^20,21^. We can use these methods to estimate *r*_*g*_ for the PLEIO analysis, if the traits come from multiple populations.

In a multi-trait analysis, we must decide which traits should be included. Selection of traits can be done based on the literature describing comorbidity, shared candidate genes, or observed genetic correlations. If we include a trait with no pleiotropy to other traits, the power will decrease. We recommend removing traits without a sufficient *r*_*g*_ to other traits. In our real data analysis, all traits had |*r*_*g*_ | > 0.15 to at least one other trait.

There exist two types of multi-trait analyses. The first is a joint analysis, in which the statistics of several traits are combined into one. The goal of this type of analysis is to find pleiotropic loci that are associated to multiple traits. These analyses have the same strengths and weaknesses as typical meta-analysis. Aggregating more traits can provide additional power, but modeling heterogeneity between traits and interpreting results can often be challenging. The second type is an augmentation analysis, in which related traits help the association test of a single trait^10-13^. The goal of this type of analysis is to maximize power for a single trait. In this study, we focused on the former type. Since our framework provides tools to facilitate interpretations, our method can minimize the weaknesses of the joint analysis.

In summary, we proposed a general and flexible framework for the identification and interpretation of pleiotropic loci. We expect that our framework can help discover core genes that contribute to multiple phenotypes, which can lead us to a better understanding of the common etiology of traits and the development of shared drug targets.

## 5 Online Methods

### 5.1 PLEIO

Here we describe our framework, PLEIO (Pleiotropic Locus Exploration and Interpretation using Optimal test). PLEIO aggregates summary statistics of multiple traits to identify pleiotropic loci. Suppose we have *Q* traits that we expect to share genetic components. We can collect *T* sets of genome-wide summary statistics for these traits. can be greater than because more than one study can be included per trait. These traits can be a mixture of qualitative and quantitative traits, whereby the quantitative traits can have differing phenotypic units. Suppose we have *M* SNPs that are shared by all studies we collected. Let 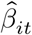 denote the effect size estimate of the *i*th SNP for the *t*th study, 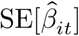 denote the standard error estimate, and *N*_*t*_ denote the number of samples in the *t*th study. Given this input, PLEIO performs a multi-trait joint analysis in the following five steps.

#### 5.1.1 Decomposition of correlation

In the input data, the effect sizes between studies can be correlated. We decompose this correlation into genetic correlation **C**_**g**_ and environmental correlation **C**_**e**_ by applying LD score regression (LDSC) to each pair of studies. **C**_**e**_ reflects the correlated errors of the effect size estimates driven by sample overlap. It is straightforward to obtain **C**_**g**_ and the heritabilities **h**^**2**^using LDSC. We can combine **C**_**g**_ and **h**^**2**^ to the genetic covariance matrix **Ω**.

It is not straightforward to obtain **C**_**e**_ using LDSC, which we describe below. We start by correcting for the study-specific confounding factor by dividing the z-scores by the square-root of the LDSC intercept. Let *z*_*ij*_ and *z*_*ik*_ denote the z-score of SNP *i* for trait *j* or after *k* this correction. We use the weighted sum of z-scores meta-analysis method to combine the two z-scores:

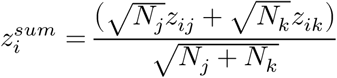

Since this approach ignores the environmental correlations, the variance of 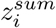 is not 1. We can show that the variance increases by 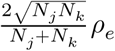, where *ρ*_*e*_ is the environmental correlation. We then use 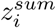 of all SNPs as input to LDSC. We can assume that the inflation of LDSC intercept reflects this variance increase, because the study-specific confounding was already corrected. Let *α*_*sum*_ be the intercept. We can estimate the environmental correlation as

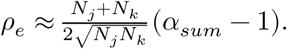

For details of the derivation, see **Supplementary Note**.

#### 5.1.2 Standardization of effect sizes

In the input data, the scales of effect sizes can be heterogeneous between the studies. We calculate the standardized effect sizes of SNP *i* for the trait *t* as

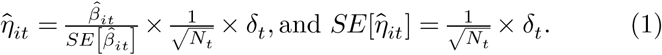

*δ*_*t*_ is a scaling factor that is 1 for quantitative trait and 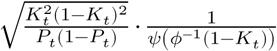 for qualitative trait, where _*t*_ refers to the disease prevalence, *P*_*t*_ = (*N*_*t*_|*y* = 1)/*N*_*t*_ refers to the sample prevalence, *ψ* refers to the probability density function of the standard normal distribution, and *ϕ*^−1^ refers to the inverse of the cumulative density function of the standard normal distribution. For quantitative traits, 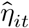 corresponds to the effect size based on the standardized phenotypes and the standardized genotypes. For qualitative traits, 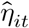 is the effect size for the liability, assuming that the z-score 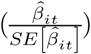 was obtained from a linear model with an observed scale (by setting the phenotypes 0 and 1). Typically, the z-scores come from the logistic regression model rather than the observed scale linear model. However, it is a common practice to use these z-scores as if they came from the linear model^22^. 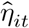 can be used conveniently to interpret the pleiotropic effects of a variant, because in contrast to the original effect size 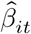, it is independent of the units of phenotypes.

#### 5.1.3 Mapping pleiotropic loci using variance component test

We build a statistical model optimized for the identification of pleiotropic loci. We assume that an individual phenotype is influenced by *K* causal SNPs, whose individual contribution is very small. For simplicity, we assume that *K* causal SNPs are shared by *T* traits. Let ***η***_*i*_ denote a *T* × 1 vector of the true effect sizes of the causal SNP *i* under the standardized scale. A common model is to assume that *K* SNPs have equal contributions. Then, 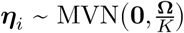, where **Ω** denotes the genetic covariance matrix, of which diagonal elements are the narrow sense heritabilities. We assume ***η***_*i*_ = **0** for non-casual SNPs.

Let 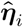 denote the observed effect size and 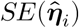 denote the standard error. We can model 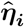 as the sum of the true genetic effect and the error:

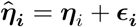

**ϵ**_***i***_ is a random variable denoting the error, which follows **ϵ**_***i***_ ∼ MVN(0, **Σ**) where 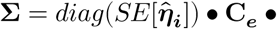 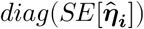. Note that **Σ** is independent of SNP *i*, because 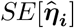 is independent of *i* as shown in equation (1). Thus, 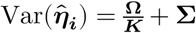 for causal SNPs and 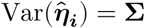 for non-causal SNPs. As described earlier, applying LDSC to 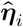 and 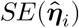 of all *M* SNPs can produce an estimate of the genetic covariance matrix 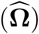 as well as the error correlation 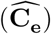.

We then relax the assumption that *K* SNPs have equal contributions. Then, the true effect ***η***_*i*_ needs not have the fixed variance 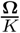. We now model 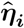 as

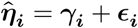

where **γ**_***i***_ is a new random variable denoting the genetic effect that follows 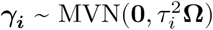, where 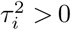 for causal SNPs and 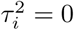 for non-causal SNPs. That is, the scaling factor 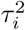 of the variance can model SNP-by-SNP differences in genetic contributions. As a special case, if we set 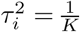 for *K* causal SNPs and 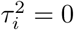 for non-casual SNPs, this model is reduced to the previous model assuming equal contributions of causal SNPs. Note that although we relaxed the assumption of the equal contribution, the variance of **γ**_***i***_ is still proportional to **Ω**, which models the relative heritability differences of the traits and the genetic correlations among the traits. Under this model, testing whether a SNP is causal or not corresponds to testing the null hypothesis 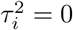 versus the alternative hypothesis 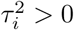.

To this end, we can fit a variance component model to get the maximum likelihood estimate (MLE) 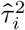 that maximizes 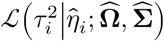.A numerical optimization algorithm such as the pseudo Newton-Raphson method can be used to find 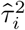. However, updating the value of the likelihood function at each iteration requires a matrix inversion. With a large *T*, this can significantly increase the overall analysis time. To solve this challenge, we developed a novel optimization technique that considerably reduces the computational burden for finding MLE (See **Supplementary Note**). The proposed optimization method carries out a linear transformation on 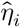 using 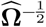. The transformed observed effect sizes follow

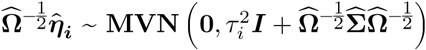

where the corresponding 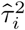 maximizes 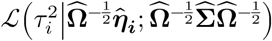 under the constraint of 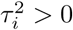. We apply a spectral decomposition 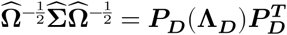. where **Λ**_**D**_ is a diagonal matrix of the eigenvalues, the diagonal elements of which are arranged in ascending order, and ***P***_***D***_ is an eigenvector matrix, the *i*th column of which corresponds to the *i*th eigenvalue. Then, we only need to calculate 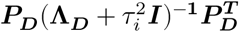 per each iteration, which is much easier to calculate than 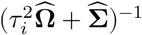. Note that the values of the matrices ***P***_***D***_ and **Λ**_***D***_ remain unchanged with iterations. As a result, we get the log-likelihood ratio test (LRT) statistic

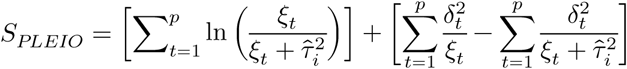

where *p* is the number of non-zero eigenvalues, *ξ*_*t*_ is the *t*th diagonal element of 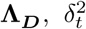 is the *t*th element of the vector 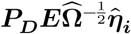, and ***E*** is a diagonal matrix of which the first *p* elements are 1 and the rest are 0. This technique can substantially shorten the time to complete our test, of which the time reduction increases with increasing number of traits (**Supplementary Figure 7 and 8**).

The underlying intuitions of our model are as follows. Our key assumption is that the genetic component **γ**_***i***_ in the effect size is a random variable whose variance is proportional to the genetic covariance matrix. ΩThis implies the following two; First, phenotypes with larger heritability show larger genetic effects. Second, phenotypes show genetic effects concordant to their genetic correlations. Because 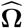 and 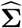 are summarized information from the whole genome, this approach can maximize the overall power. In that sense, our model resembles empirical Bayes approaches^12^.

#### 5.1.4 Assessing statistical significance via importance sampling

Here we describe how to assess an accurate p-value of our LRT statistic, *S*_*PLEIO*_. *S*_*PLEIO*_ asymptotically follows a 50:50 mixture of 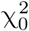 and 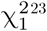. However, the asymptotic approximation is only accurate if the sample size is large. Unfortunately, the sample size of our statistic is *T*, the number of combined studies, and not the total number of individuals that were used to generate *T* statistics. Since *T* is typically small (< 100), the asymptotic distribution does not approximate the null distribution well. Moreover, when *T* is small, it turns out that the null distribution depends on the genetic covariance matrix and error correlation matrix 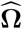 and 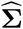 Thus, an alternative approach would be simulating null distributions based on the study-specific factors (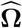 and 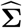). But the standard sampling is overly inefficient for assessing very small p-values (e.g. 5 × 10^−8^).

Instead, we use a novel importance sampling approach to assess the p-value of *S*_*PLEIO*_. Let ***x*** be a random variable denoting the standardized effect sizes. Let ***q***(***x***) denote the probability density function (PDF) of ***x*** under the null. One advantage of using the standardized scale is that the error variance 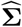 is independent of SNP *i*. Thus, 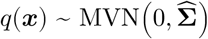 regardless of SNPs. By definition, ∫_***D***_ ***q***(***x***) ***dx*** = 1 where ***D*** = ℝ^*T*^. We can consider *S*_*PLEIO*_ as a function of ***x*** given 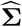 and 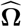. Given an observed *S*_*PLEIO*_ statistic from data, which we call ***θ***, we want to calculate the p-value of it. To this end, let *f* (***x, θ***) denote an indicator function as follows:

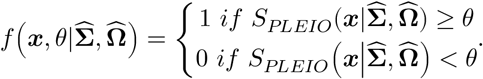

For simplicity, we replace 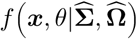 with a simpler expression, *f*(***x***). The p-value of ***θ*** can be expressed as:

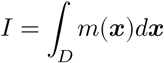

where *m*(***x***) = *f* (***x***)***q***(***x***). To estimate *I*, we can exploit importance sampling algorithm. In importance sampling, we use a sampling distribution *p*(***x***) that differs from ***q***(***x***). Let ***X***^***p***^ ∼ *p*(***x***) denote a *M* × *T* matrix of the sampled effect sizes generated from *p*(***x***), where *M* is the number of sampling. Then, we can estimate *I* using ***X***^***p***^ as follows:

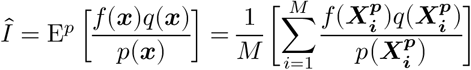

where E^*p*^[⋅] denotes the expectation over ***X***^***p***^, and 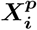 is the *i*th row vector of ***X***^***p***^.

The challenge in importance sampling is to choose an appropriate *p*(***x***). It is particularly challenging in GWAS because the range of p-values is very wide, from 1.0 to 5 × 10^−8^. Thus, it is difficult to select a single distribution that can minimize variance for all range of p-values. To solve this challenge, we generate samples from a mixture distribution. Let *p*_*j*_ (***x***) denote the *j*th sampling distribution where *j* = {1,2, …, *K*}. We let *p*_*α*_(***x***) denote the mixture distribution of *K* sampling distributions. We assume that each sampling distribution has the equal chance to generate a sample such that 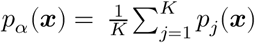. Detailed information on the selection of *p*_*j*_(***x***) can be found in **Supplementary Note**. In our method, we use *p*_*j*_(***x***) as a control variate of *m*(***x***) = *f* (***x***)***q***(***x***), which let us define:

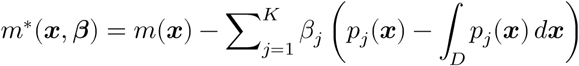

where E[*m*^∗^] = E[*m*] = *I*, and ∫_***D***_ *p*_*j*_(***x***) ***dx*** = 1. The control variate method maximizes the variance reduction of Var(*m*) using the optimal control variate coefficient (***β***^∗^). Then, the variance Var(*m*^∗^) becomes equal to or smaller than Var(*m*). In PLEIO, we use the following control variate approach suggested by Owen and Zhou^24^:

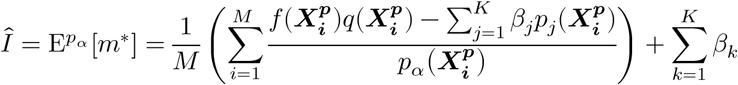

Given ***X***^***p***^ from *p*_*α*_(***x***), we calculate p-values of equally spaced 50 different ***θ*** that are in the range (0,40), which roughly correspond to p-values from 1.0 to 3 × 10^−11^. For each ***θ***, we calculate the optimal ***β*** for the control variate method to maximize the variance reduction of the p-value estimate. See **Supplementary Note** for how we obtained the optimal control variate coefficients (***β***^∗^). Using these 50 points, we interpolate p-values for ***θ*** < 40 using B-spline fit and extrapolate p-values for ***θ*** > 40 using the linear fit on the logarithmic p-value scale.

#### 5.1.5 Pleiotropy plot

PLEIO offers a tool to visualize the pleiotropic effects of a SNP, which we named *pleiotropy plot* (**Figure 4**). This circular plot provides information about the standardized effect sizes, the local heritabilities, and the local Manhattan plots of a SNP. The outer part is partitioned by the traits, each of which contains (1) the effect size of each trait on the original scale as text and on the standardized scale as a horizontal bar, and (2) the local Manhattan plot within a 1 Mb window. The inner part is a ribbon plot linking multiple traits. The ribbon color indicates genetic correlations, and the width at the end indicates the locus heritability (squared standardized effect size).

### 5.2 Data analysis

#### 5.2.1 Collection of GWAS summary statistics

We collected public GWAS summary statistics of 18 traits related to cardiovascular disease from large-scale genetic consortia, as described in **Supplementary table 3**. When a consortium database contained more than one GWAS for the same phenotype, we selected the latest study. We obtained the summary statistics of four quantitative traits from Global Lipids Genetics consortium^17^. The data consisted of the results of GWASs from 94,595 individuals from 23 studies genotyped with GWAS arrays and 93,982 individuals from 37 studies genotyped with the Metabochip array. We obtained the summary statistics of the twelve binary traits in the UK biobank data from the Neale lab website (http://www.nealelab.is/uk-biobank). The data consisted of the results of GWASs from 361,193 individuals in the UK biobank cohort. We obtained the summary data on coronary artery disease (CAD) from CARDIo+C4D consortium^16^. The data consisted of the results of GWAS meta-analysis from 60,801 CAD cases and 123,504 controls from 48 studies. We obtained the summary data on fasting glucose (FG) from MAGIC consortium^19^. The data consisted of the analysis results from 46,186 non-diabetic patients from 21 GWA studies. All samples were Caucasians with European descent. The genotypes of all summary statistics were coordinated to GRCh37 (Hg19).

#### 5.2.2 Summary statistics data QC

For each summary statistics dataset, we removed SNPs that were not included in 1000 Genomes^25^. We checked the consistency of allele pair of each SNP with the corresponding allele pair of the SNP in 1000 Genomes. To eliminate potential strand mismatches, we pruned SNPs with the allele pair GC and AT. The genetic covariance and error correlation were estimated from summary statistics of the remaining SNPs. A total of 1,799,044 SNPs was included in the joint analysis of 18 traits.

#### 5.2.3 Identification of novel pleiotropic loci

In the joint analysis of 18 traits, we identified 10,041 SNPs that were genome-wide significant (*P*_*PLEIO*_ < 5 × 10^−8^). We clumped these SNPs with threshold (*r*^2^ < 0.1) and found 618 approximately independent hits. To estimate LD between SNPs, we used the European samples in the 1000 Genomes data. To determine whether the remaining variants were novel loci, we excluded variants that met any of the following two conditions: (1) The variant had a moderate LD (*r*^2^ > 0.1) with a variant that is listed in GWAS catalog as associated with the CVD-related traits, or (2) The variant already reached the genome-wide significance threshold of 5 × 10^−8^ in the original summary statistics of a single trait. As a result, we identified 20 novel pleiotropic variants.

## 7 Data availability

The summary statistics data used for the multi-trait association analysis are available from UK biobank GWAS results (http://www.nealelab.is/uk-biobank), Global Lipids Genetics consortium (http://lipidgenetics.org), CARDIo+C4D consortium (http://www.cardiogramplusc4d.org), and MAGIC consortium (https://www.magicinvestigators.org/downloads/). The multi-trait association results are available upon request.

## 8 Code availability

PLEIO is freely available at https://github.com/hanlab-SNU/PLEIO.

