## Supplementary Note for "A method to map and interpret pleiotropic loci using summary statistics of multiple traits"

### 1 Estimation of the environmental correlation between a pair of traits.

#### 1.1 LDSC framework

We first describe the LDSC framework of Bulik-Sullivan et. al., (2015)[1,2]. Let $A$ and $B$ denote two different traits. We assume that we genotyped $N_A$ and $N_B$ samples at $M$ SNPs. Let $y_A$ and $y_B$ denote $N_A \times 1$ and $N_B \times 1$ vectors of phenotypes, and let $\boldsymbol{G_A}$ and $\boldsymbol{G_B}$ denote $N_A \times M$ and $N_B \times M$ genotype matrices. $\boldsymbol{G_A}$ and $\boldsymbol{G_B}$ are standardized so that each column follows $N(0{,}1)$. The z-scores of the SNP $j$ can be obtained as $z_{A,j} := \mathrm{G}_{A,j}^T y_A / \sqrt{N_A}$ and $z_{B,j} := \mathrm{G}_{B,j}^T y_B / \sqrt{N_B}$. Let $\boldsymbol{z_A}$ and $\boldsymbol{z_B}$ denote $M \times 1$ vectors of z-scores of traits $A$ and $B$.

Bulik-Sullivan et. al., (2015) derived the following equations[1,2]:

$$E[z_{A,j}^2|\ell_j] = \frac{N_A h_A^2}{M}\ell_j + N_A\alpha_A + 1$$
$$E[z_{B,j}^2|\ell_j] = \frac{N_B h_B^2}{M}\ell_j + N_B\alpha_B + 1 \qquad (1)$$

where $h_A^2$ and $h_B^2$ are narrow sense heritabilities, and $\ell_j$ is the value of LD-score of $j$th SNP, which can be obtained from external reference. LDSC uses both summary statistics $(S_A = \{N_A, \boldsymbol{z_A}\}, S_B = \{N_B, \boldsymbol{z_B}\})$ and LD-score information $(\ell_j, where\ j \in \{1{,}2{,}\dots, M\})$ to estimate the trait heritability ($h_A^2$ and $h_B^2$) along with the intercepts ($N_A\alpha_A + 1$ and $N_B\alpha_B + 1$). To estimate genetic correlation, LDSC uses the following equality:

$$E[z_{A,j} z_{B,j}|\ell_j] = \frac{\sqrt{N_A N_B}\sigma_g}{M}\ell_j + \frac{\sigma N_{AB}}{\sqrt{N_A N_B}} \qquad (2)$$

where $\sigma_g$ denotes the genetic correlation between trait A and B, $N_{AB}$ denotes the shared individuals between samples $N_A$ and $N_B$, and $\sigma$ denotes phenotypic correlations of the shared individuals.

#### 1.2 Environmental correlation

To estimate environmental correlation, we start by correcting for study-specific confounding factors. We define $\Psi(S_K)$ as a function that accepts the summary statistics $S_K$ as input and gives the LDSC intercept value $1+\gamma_K$, where $\gamma_K$ reflects the confounding effects in $S_K$. We can adjust for the confounding effect by dividing z-scores by $\sqrt{1+\gamma_K}$. That is, we replace $S_A$ with $S_A^c = \{N_A, \boldsymbol{z_A^c}\}$ where $\boldsymbol{z_A^c} = \{\frac{z_{A,1}}{\sqrt{1+\gamma_A}}, \frac{z_{A,2}}{\sqrt{1+\gamma_A}}, \dots, \frac{z_{A,M}}{\sqrt{1+\gamma_A}}\}$. Then, $\Psi(S_A^c) = 1$. We apply this confounding elimination procedure to every trait.

Our strategy is to apply the weighted sum of z-scores to combine z-scores of A and B.

$$z_{AB,j} = \frac{\sqrt{N_A} z_{A,j}^c + \sqrt{N_B} z_{B,j}^c}{\sqrt{N_A + N_B}} \qquad (3)$$

Previous studies have shown that the weighted sum of z-scores is approximately equivalent to the popular inverse-variance weighted method[3]. Let $S_{AB}$ denote the summary statistics for $z_{AB,j}$ so that $S_{AB} \in \{N_{A+B}, \boldsymbol{z_{AB}}\}$. We apply LDSC to estimate the results of $\Psi(S_{AB})$ as follows:

$$\Psi(S_{AB}) = 1 + \gamma_{AB}.$$

*No sample overlap*

We can consider two scenarios. First, we suppose that we do not have sample overlap. Then, we will observe no inflation of intercept from $z_{AB,j}$. This is because of the following:

$$E[(z_{AB,j})^2 | \ell_j] = E\left[\frac{N_A\left(z_{A,j}^c\right)^2 + 2\sqrt{N_A N_B} z_{A,j}^c z_{B,j}^c + N_B (z_{B,j}^c)^2}{N_A + N_B} | \ell_j\right]$$
$$= \frac{\left[N_A E\left[\left(z_{A,j}^c\right)^2 | \ell_j\right] + \sqrt{N_A N_B} \mathrm{E}[2 z_{A,j}^c z_{B,j}^c | \ell_j] + N_B \mathrm{E}[(z_{B,j}^c)^2 | \ell_j]\right]}{N_A + N_B}$$
$$= \frac{1}{N_A + N_B}\left[\frac{N_A^2 h_A^2}{M} + \frac{N_B^2 h_B^2}{M} + 2\frac{N_A N_B \sigma_g}{M}\right] \ell_j + 1 + \frac{2\sigma N_{AB}}{N_A + N_B}$$

The last term is zero when there is no sample overlap. Thus, $\gamma_{AB} = 0$.

*Sample overlap*

The second scenario is that we have sample overlap. Then environmental correlations occur. It is convenient to decompose z-scores to the genetic effect and the environmental error. Let $\boldsymbol{\epsilon_K}$ denote random error and let $\boldsymbol{g_K}$ denote the genetic effect of $\boldsymbol{z_K^c}$ so that $z_{K,j}^c = g_{K,j} + \epsilon_{K,j}$. By definition, $\mathrm{E}[\epsilon_{K,j}] = 0$ , $\mathrm{Var}[\epsilon_{K,j}] = 1$ , $\mathrm{E}[g_{K,j}] = 0$ , and $\mathrm{Var}[g_{K,j}] = \frac{N_K h_K^2}{M}\ell_j$ where $\mathrm{Cov}(\epsilon_{K,j}, g_{K,j}) = 0$ . Similarly, $z_{A,j}^c = g_{A,j} + \epsilon_{A,j}$ and $z_{B,j}^c = g_{B,j} + \epsilon_{B,j}$ . Let $\rho_e$ denote the environmental correlation. Simply, $\rho_e = \mathrm{Cor}(\epsilon_{A,j}, \epsilon_{B,j})$.

Then, the following equation holds

$E[(z_{AB,j}|N_{AB} \neq 0)^2|\ell_j]$

$$= \mathrm{E}\left[\left(\frac{\sqrt{N_A}(g_{A,j} + \epsilon_{A,j}) + \sqrt{N_B}(g_{B,j} + \epsilon_{B,j})}{\sqrt{N_A + N_B}}|N_{AB} \neq 0\right)^2 |\ell_j\right]$$

$$= f(\ell_j) + \left[\frac{N_A}{N_A + N_B}\mathrm{Var}(\epsilon_{A,j}) + \frac{N_B}{N_A + N_B}\mathrm{Var}(\epsilon_{B,j}) + 2\sqrt{N_A N_B}\mathrm{Cov}(\epsilon_{A,j}, \epsilon_{B,j})\right]$$

$$= f(\ell_j) + \left[1 + 2\frac{\sqrt{N_A N_B}}{N_A + N_B}\rho_e\right]$$

where $f(\ell_j)$ is a first-order function of $\ell_j$ without a constant term. Therefore, $\gamma_{AB} = 2\frac{\sqrt{N_A N_B}}{N_A + N_B}\rho_e$ is the inflation of the intercept caused by environmental correlation. Since we can estimate $\gamma_{AB}$ from LDSC, we can estimate the environmental correlation as follows:

$$\rho_e = \frac{N_A + N_B}{2\sqrt{N_A N_B}}\gamma_{AB}.$$

### 2 Optimization strategy for variance component test

#### 2.1 LRT statistic

In this section, we describe our optimization strategy that increases the computational efficiency for determining the maximum likelihood estimate (MLE) in the variance component test of PLEIO. Below, we use the letter $L$ to denote the number of traits in the multi-trait joint analysis (not $T$, because we use the letter $T$ to denote the matrix transpose). Let $\hat{\boldsymbol{\eta}}_{\boldsymbol{i}}$ denote a $L \times 1$ vector representing the observed standardized effect sizes for SNP $i$ and $SE(\hat{\boldsymbol{\eta}}_{\boldsymbol{i}})$ denote the vector of the corresponding standard errors. Let $\widehat{\boldsymbol{\Omega}}$ denote a $L \times L$ matrix representing the genetic covariance matrix of $L$ traits, and $\widehat{\boldsymbol{\Sigma}}$ denote a $L \times L$ matrix representing the environmental covariance matrix. Note that $\widehat{\boldsymbol{\Sigma}} = diag\big(SE(\hat{\boldsymbol{\eta}}_{\boldsymbol{i}})\big) \bullet \hat{\mathbf{C}}_{\mathbf{e}} \bullet diag\big(SE(\hat{\boldsymbol{\eta}}_{\boldsymbol{i}})\big)$, where $\hat{\mathbf{C}}_{\boldsymbol{e}}$ is a $L \times L$ matrix representing the environmental correlation matrix, and $diag\big(SE(\hat{\boldsymbol{\eta}}_{\boldsymbol{i}})\big)$ is a diagonal matrix whose diagonal values are $SE(\hat{\boldsymbol{\eta}}_{\boldsymbol{i}})$. As described, $\widehat{\boldsymbol{\Sigma}}$ is independent of SNP $i$ under the standardized scale. The PLEIO's statistic is a log-likelihood ratio test statistic (LRT). The likelihood functions under the null and alternative hypotheses can be shown as follows:

$$\mathcal{L}_0\big(\cdot\,\big|\hat{\boldsymbol{\eta}}_{\boldsymbol{i}};\widehat{\boldsymbol{\Sigma}}\big) = \frac{1}{(2\pi)^{L/2}\big|\widehat{\boldsymbol{\Sigma}}\big|^{1/2}}\exp\left(-\frac{1}{2}\hat{\boldsymbol{\eta}}_{\boldsymbol{i}}^{\boldsymbol{T}}\widehat{\boldsymbol{\Sigma}}^{-\mathbf{1}}\hat{\boldsymbol{\eta}}_{\boldsymbol{i}}\right)$$

$$\mathcal{L}_1\big(\tau^2\big|\hat{\boldsymbol{\eta}}_{\boldsymbol{i}};\widehat{\boldsymbol{\Omega}},\widehat{\boldsymbol{\Sigma}}\big) = \frac{1}{(2\pi)^{L/2}\big|\widehat{\boldsymbol{\Omega}}\tau_i^2+\widehat{\boldsymbol{\Sigma}}\big|^{1/2}}\exp\left(-\frac{1}{2}\hat{\boldsymbol{\eta}}_{\boldsymbol{i}}^{\boldsymbol{T}}\big(\widehat{\boldsymbol{\Omega}}\tau_i^2+\widehat{\boldsymbol{\Sigma}}\big)^{-\mathbf{1}}\hat{\boldsymbol{\eta}}_{\boldsymbol{i}}\right),$$

and the corresponding LRT statistic is

$$S_{PLEIO} = -2\ \ln\left[\frac{\mathcal{L}_0\big(\cdot\,\big|\hat{\boldsymbol{\eta}}_{\boldsymbol{i}};\widehat{\boldsymbol{\Sigma}}\big)}{\sup\big\{\mathcal{L}_1\big(\tau_i^2\big|\hat{\boldsymbol{\eta}}_{\boldsymbol{i}};\widehat{\boldsymbol{\Omega}},\widehat{\boldsymbol{\Sigma}}\big)\colon \tau^2 \geq 0\big\}}\right].$$

#### 2.2 Efficient optimization

Our goal is to find $\tau^2$ that satisfies $\sup\big\{\mathcal{L}_1\big(\tau_i^2\big|\hat{\boldsymbol{\eta}}_{\boldsymbol{i}};\widehat{\boldsymbol{\Omega}},\widehat{\boldsymbol{\Sigma}}\big)\colon \tau_i^2 \geq 0\big\}$ under the alternative model, $\hat{\boldsymbol{\eta}}_{\boldsymbol{i}} \sim \mathrm{MVN}\big(\mathbf{0}, \tau^2\widehat{\boldsymbol{\Omega}} + \widehat{\boldsymbol{\Sigma}}\big)$, to obtain the LRT statistic. To find MLE $\hat{\tau}^2$, one possible way is to use an iterative optimization technique (e.g. quasi Newton's method). This optimization, however, requires a burdensome calculation of the matrix inversion of the variance covariance matrix in the function $\mathcal{L}_1$

for each iteration. Instead, we propose a novel optimization technique that avoids the repeated inversions.

We first define a spectral (eigen) decomposition of a symmetric and positive semidefinite matrix $\mathbf{A}$ of size $L \times L$ as follows: let $\boldsymbol{\xi_A}$ denote the $L \times 1$ vector representing eigen values of $\mathbf{A}$, and $\boldsymbol{P_A}$ denote the $L \times L$ matrix whose $i$th column vector indicates the corresponding eigenvector. By definition, $\boldsymbol{P_A}$ is an orthonormal matrix so that $\boldsymbol{P_A^{-1}} = \boldsymbol{P_A^T}$. Suppose the values of $\boldsymbol{\xi_A}$ are sorted in descending order, and the corresponding column order of $\boldsymbol{P_A}$ is also sorted as well. Let $\xi_{A,i}$ be the $i$th eigen value. Then, $\xi_{A,i} \geq 0$ where $i = \{1,2, \dots, L\}$.

Note that any covariance matrix (including $\widehat{\boldsymbol{\Omega}}$ and $\widehat{\boldsymbol{\Sigma}}_{\boldsymbol{i}}$) is symmetric and positive semidefinite by definition. Let $\widehat{\boldsymbol{\Omega}}^g$ be the generalized (pseudo) inverse of $\widehat{\boldsymbol{\Omega}}$. Since $\frac{1}{\xi_{\widehat{\boldsymbol{\Omega}},i}} > 0$ for all non-zero $\xi_{\widehat{\boldsymbol{\Omega}},i}$, $\widehat{\boldsymbol{\Omega}}^g$ is also positive semidefinite (PSD) and symmetric, as is $\left[\widehat{\boldsymbol{\Omega}}^g\right]^{\frac{1}{2}}$. Then, we apply the following linear transformation to $\widehat{\boldsymbol{\eta}}_{\boldsymbol{i}}$:

$$\left[\widehat{\boldsymbol{\Omega}}^g\right]^{\frac{1}{2}}\widehat{\boldsymbol{\eta}}_{\boldsymbol{i}} \sim MVN\left(\mathbf{0}, \left[\widehat{\boldsymbol{\Omega}}^g\right]^{\frac{1}{2}}\widehat{\boldsymbol{\Sigma}}\left[\widehat{\boldsymbol{\Omega}}^g\right]^{\frac{1}{2}} + \tau_i^2 I\right)$$

whose log-likelihood functions under the null and alternative hypotheses can be shown as:

$$\ell_0'\left(\cdot\left|\left[\widehat{\boldsymbol{\Omega}}^g\right]^{\frac{1}{2}}\widehat{\boldsymbol{\eta}}_{\boldsymbol{i}}; \boldsymbol{D}\right.\right) = -\frac{1}{2}\left[L\ln(2\pi) + \ln(|\boldsymbol{D}|) + \left(\left[\widehat{\boldsymbol{\Omega}}^g\right]^{\frac{1}{2}}\widehat{\boldsymbol{\eta}}_{\boldsymbol{i}}\right)^{\boldsymbol{T}} \boldsymbol{D^{-1}}\left(\left[\widehat{\boldsymbol{\Omega}}^g\right]^{\frac{1}{2}}\widehat{\boldsymbol{\eta}}_{\boldsymbol{i}}\right)\right]$$

and

$$\ell_1'\left(\tau_i^2\left|\left[\widehat{\boldsymbol{\Omega}}^g\right]^{\frac{1}{2}}\widehat{\boldsymbol{\eta}}_{\boldsymbol{i}}; \boldsymbol{K}\right.\right) = -\frac{1}{2}\left[L\ln(2\pi) + \ln(|\boldsymbol{K}|) + \left(\left[\widehat{\boldsymbol{\Omega}}^g\right]^{\frac{1}{2}}\widehat{\boldsymbol{\eta}}_{\boldsymbol{i}}\right)^{\boldsymbol{T}} \boldsymbol{K^{-1}}\left(\left[\widehat{\boldsymbol{\Omega}}^g\right]^{\frac{1}{2}}\widehat{\boldsymbol{\eta}}_{\boldsymbol{i}}\right)\right],$$

where $\boldsymbol{D} = \left[\widehat{\boldsymbol{\Omega}}^g\right]^{\frac{1}{2}}\widehat{\boldsymbol{\Sigma}}\left[\widehat{\boldsymbol{\Omega}}^g\right]^{\frac{1}{2}}$ and $\boldsymbol{K} = \boldsymbol{D} + \tau_{\boldsymbol{i}}^2\boldsymbol{I}$. The product of two symmetric real PSD matrices, $\mathrm{A}^{-\frac{1}{2}}B\mathrm{A}^{-\frac{1}{2}}$, is also symmetric real PSD, and the value of $\tau_{\boldsymbol{i}}^2$ is strictly non-negative. Therefore, $\boldsymbol{D}$ and $\boldsymbol{K}$ are symmetric real PSD, and are covariance matrices. Note that by applying this transformation, we made the second term in $\boldsymbol{K}$ a diagonal matrix. Under this condition, we can apply a optimization technique similar to ones used in EMMA[4] or RE2C[5]. The following equalities hold:

$$\begin{aligned}\boldsymbol{K}\boldsymbol{P_D} &= (\boldsymbol{D}+\tau^2\boldsymbol{I})\boldsymbol{P_D}\\ &= (\boldsymbol{D}\boldsymbol{P_D}+\boldsymbol{P_D}\tau^2) = \boldsymbol{P_D}\boldsymbol{\Lambda_D}+\boldsymbol{P_D}\tau^2\\ &= \boldsymbol{P_D}(\boldsymbol{\Lambda_D}+\tau^2\boldsymbol{I}),\end{aligned}$$

and

$$\boldsymbol{K} = \boldsymbol{P_D}(\boldsymbol{\Lambda_D}+\tau_i^2\boldsymbol{I})\boldsymbol{P_D^T}$$

where $\boldsymbol{\Lambda_D}$ is a $L\times L$ diagonal matrix whose $i$th element is $\xi_{D,i}$. By the definition of spectral decomposition, the following statements are true:

1. For a real symmetric PSD matrix $\boldsymbol{A}$, $|\boldsymbol{A}| = \prod_{i=1}^{p}\xi_{A,i}$.
2. $\boldsymbol{A^g} = (\boldsymbol{P_A}\boldsymbol{E})^{\boldsymbol{T}}[\boldsymbol{\Lambda_A^+}]^{-\mathbf{1}}(\boldsymbol{P_A}\boldsymbol{E})$

where $p$ is the number of positive eigenvalues of $\mathbf{A}$, $\boldsymbol{E}$ is a diagonal matrix whose first $p$ diagonal elements are 1 and 0 otherwise, and $\boldsymbol{\Lambda_A^+}$ is a $p\times p$ diagonal matrix whose $i$th diagonal element is $\xi_{A,i}$. Then, we can rewrite $\ell_1'$ as follows:

$$\begin{aligned}\ell_1' &= -\frac{1}{2}\left[L\ln(2\pi)+\sum\nolimits_{t=1}^{p}\ln(\xi_{D,t}+\tau_{\boldsymbol{i}}^2)+\left(\left[\widehat{\boldsymbol{\Omega}}^g\right]^{\frac{1}{2}}\hat{\boldsymbol{\eta}}_{\boldsymbol{i}}\right)^{\boldsymbol{T}}(\boldsymbol{P_D}\boldsymbol{E})^{\boldsymbol{T}}[\boldsymbol{\Lambda_D^+}]^{-\mathbf{1}}(\boldsymbol{P_D}\boldsymbol{E})\left(\left[\widehat{\boldsymbol{\Omega}}^g\right]^{\frac{1}{2}}\hat{\boldsymbol{\eta}}_{\boldsymbol{i}}\right)\right]\\ &= -\frac{1}{2}\left[L\ln(2\pi)+\sum\nolimits_{t=1}^{p}\ln(\xi_{D,t}+\tau_{\boldsymbol{i}}^2)+\left(\boldsymbol{P_D}\boldsymbol{E}\left[\widehat{\boldsymbol{\Omega}}^g\right]^{\frac{1}{2}}\hat{\boldsymbol{\eta}}_{\boldsymbol{i}}\right)^{\boldsymbol{T}}[\boldsymbol{\Lambda_D^+}]^{-\mathbf{1}}\left(\boldsymbol{P_D}\boldsymbol{E}\left[\widehat{\boldsymbol{\Omega}}^g\right]^{\frac{1}{2}}\hat{\boldsymbol{\eta}}_{\boldsymbol{i}}\right)\right]\\ &= -\frac{1}{2}\left[L\ln(2\pi)+\sum\nolimits_{t=1}^{p}\ln(\xi_{D,t}+\tau_{\boldsymbol{i}}^2)+\sum_{t=1}^{p}\frac{\delta_t^2}{\xi_{D,t}+\tau_i^2}\right]\end{aligned}$$

where $\delta_t^2$ is the $t$th element of the vector $\boldsymbol{P_D}\boldsymbol{E}\left[\widehat{\boldsymbol{\Omega}}^g\right]^{\frac{1}{2}}\hat{\boldsymbol{\eta}}_{\boldsymbol{i}}$. Using the equation above, we can derive the first and second derivative of the likelihood function $\ell_1'$ as:

$$\frac{d\ell_1'}{d\tau_{\boldsymbol{i}}^2} = -\frac{1}{2}\left[\sum\nolimits_{t=1}^{p}\frac{1}{\xi_{D,t}+\tau_{\boldsymbol{i}}^2}-\sum_{t=1}^{p}\frac{\delta_t^2}{(\xi_{D,t}+\tau_i^2)^2}\right]$$

$$\frac{d^2\ell_1'}{d(\tau_{\boldsymbol{i}}^2)^2} = -\frac{1}{2}\left[\sum\nolimits_{t=1}^{p} \frac{1}{(\xi_{D,t}+\tau_{\boldsymbol{i}}^2)^2} + 2\sum_{t=1}^{p} \frac{\delta_t^2}{(\xi_{D,t}+\tau_i^2)^3}\right].$$

In PLEIO, we use $\frac{d\ell_1'}{d\tau_{\boldsymbol{i}}^2}$ as the root function of the Newton Raphson algorithm and $\frac{d^2\ell_1'}{d(\tau_{\boldsymbol{i}}^2)^2}$ as its first derivative function. The initial value of the algorithm can be found using a grid search algorithm. We generate multiple values of $\tau_{\boldsymbol{i}}^2$ ranged from $10^{-9}$ to $10^6$ and use $\tau_{\boldsymbol{i}}^2$ that maximizes the likelihood as an initial value. Finally, we estimate the value of PLEIO as follows:

$$S_{PLEIO} = \left[\sum\nolimits_{t=1}^{p} \ln\left(\frac{\xi_{D,t}}{\xi_{D,t}+\hat{\tau}_i^2}\right)\right] + \left[\sum_{t=1}^{p} \frac{\delta_t^2}{\xi_{D,t}} - \sum_{t=1}^{p} \frac{\delta_t^2}{\xi_{D,t}+\hat{\tau}_i^2}\right]$$

This simple form of $S_{PLEIO}$ shows that our method can optimize $\tau_{\boldsymbol{i}}^2$ with a single matrix inversion of $\boldsymbol{D}$. Recall that optimization of $\tau_{\boldsymbol{i}}^2$ using a naïve quasi Newton Raphson approach would have required multiple matrix inversions. We tested the computing efficiency of our method by comparing it to that of the standard approach using the optimization function implemented in the python scipy library (scipy.optimize.minimize) taking into account the two variance covariance matrices. **Supplementary Figure 7 and 8** show that the computational time of the proposed model was faster than the time of the standard optimization. The reduction rate of computational time was approximately linear in relation to the number of studies, where the reduction rate increased approximately 8% per number of studies. In other words, the proposed model can compute 16-fold (1600%) faster than scipy.optimization for cross-disease joint analysis of 200 studies (8%×200=1600%) (**Supplementary Figure 8**).

### 3 P-value estimation

#### 3.1 Problem definition

We describe how PLEIO estimates the p-value. Suppose we have an observed statistic $\hat{S}_{PLEIO}$. Then, the p-value is $\mathrm{P}\left(S_{PLEIO} \geq \hat{S}_{PLEIO} \middle| H_0\right)$. Let $\boldsymbol{x}$ denote a random variable representing the standardized effect size $\boldsymbol{\eta}$, $\widehat{\boldsymbol{\Omega}}$ denote the $L \times L$ genetic covariance matrix, and $\widehat{\boldsymbol{\Sigma}}$ denote the $L \times L$ environmental covariance matrix. In the following, we will assume that the estimated $\widehat{\boldsymbol{\Sigma}}$ represents the true environmental variance. Under the null, $\boldsymbol{x}$ will follow $\mathrm{MVN}\left(0, \widehat{\boldsymbol{\Sigma}}\right)$. Let the statistic $S_{PLEIO}$ denote a function of $\boldsymbol{x}$ given $\widehat{\boldsymbol{\Omega}}$ and $\widehat{\boldsymbol{\Sigma}}$. We can define an indicator function

$$f\left(\boldsymbol{x}, \theta | \widehat{\boldsymbol{\Sigma}}, \widehat{\boldsymbol{\Omega}}\right) = \begin{cases} 1 \; if \; S_{PLEIO}(\boldsymbol{x} | \widehat{\boldsymbol{\Sigma}}, \widehat{\boldsymbol{\Omega}}) \geq \theta \\ 0 \; if \; S_{PLEIO}\left(\boldsymbol{x} \middle| \widehat{\boldsymbol{\Sigma}}, \widehat{\boldsymbol{\Omega}}\right) < \theta \end{cases}.$$

For simplicity, we replace $f\left(\boldsymbol{x}, \theta | \widehat{\boldsymbol{\Sigma}}, \widehat{\boldsymbol{\Omega}}\right)$ with a simpler expression, $f(\boldsymbol{x})$. Let $q(\boldsymbol{x})$ be the probability density function of $\mathrm{MVN}\left(0, \widehat{\boldsymbol{\Sigma}}\right)$. For a given observed LRT statistic $\theta$, the p-value can be defined as the expected value of the integrand $f(\boldsymbol{x})q(\boldsymbol{x})$ as follows,

$$I = \int_D f(\boldsymbol{x}) q(\boldsymbol{x}) dx$$

where $D = \mathbb{R}^L$. Our goal is to estimate $I$ accurately and efficiently.

#### 3.2 Asymptotic approach

The simplest way to approximate the p-value $I$ is to use the asymptotic distribution. In general, an LRT statistic using the value of the likelihood for random components in a linear mixed model asymptotically follows a mixture of Chi-squared distributions under the null. In our situation, since $\tau^2$ is a non-negative variance estimate, the statistics asymptotically follow a 50:50 mixture of one and zero degree of freedom Chi-square distributions[6]. In a cross-disease joint analysis, however, the validity of this asymptote holds if only if the number of combined studies is large. In practice, it is uncommon to combine more than 100 statistics. Therefore, the asymptotic approach will not be an exact solution for us.

##### 3.3 Standard Monte Carlo approach

A possible alternative is Monte Carlo approach. In Monte Carlo method, we repeatedly draw samples from $q$. Let $\boldsymbol{X^q}$ denote a set of samples generated from $q$. Then, $I = \mathrm{E}^{\mathrm{q}}[f(\boldsymbol{X^q})]$ where $\mathrm{E}^{\mathrm{q}}[\cdot]$ denotes expectation for $\boldsymbol{X^q} \sim q$. Assume that we have sampled $N$ observations and let $\boldsymbol{X_i^q}$ be the $i$th observation. The Monte Carlo estimator of $I$ is $\hat{I} = \frac{1}{N}\sum_{i=1}^{N} f(\boldsymbol{X_i^q})$. However, the use of Monte Carlo integration can be computationally intensive if the region of interest in $\boldsymbol{X}$ is located at the tails of $q$ distribution. In genome-wide analyses, the p-value of interest is often as small as $5 \times 10^{-8}$. To get reasonable accuracy for such a small value, more than 10 million samples are required. This can be computationally intensive since the maximum likelihood estimation must be carried out for each sample to calculate $S_{PLEIO}$.

In previous studies, the standard Monte Carlo approach was used in the context of meta-analysis of GWAS. Han and Eskin adapted a strategy to pre-calculate $I$ for every $\theta$ using the Monte Carlo sampling[7]. For each possible number of studies ($L$), they generated 10M null samples and tabulated the relationship between $I$ and $\theta$. This was possible because they need not assume any genetic correlation or environmental correlation. Since their statistical model did not include any correlations, the standard MVN with mean zero and variance $\boldsymbol{I}$ (identity matrix) represented the null distribution of various situations well; therefore, pre-sampling from that distribution was sufficient. In a subsequent study, Cue et al.[5] extended the model to include the environmental correlation caused by sample overlap. Because the environmental correlation can vary from situation to situation, it was not possible to calculate the table in advance for all possible situations. Fortunately, the environmental correlation always becomes positive if it is due to a sample overlap of controls. Inspired by this, Cue et al. [5] developed a heuristic to summarize the strength of overall positive correlations between studies in one value (the average correlation $\bar{r}$), and tabulated $I$ for each possible $\bar{r}$. Although these previous studies have used the standard Monte Carlo approach or its variation, we cannot apply these approaches directly in our context. This is because every analysis will have unique $\widehat{\boldsymbol{\Omega}}$ and $\widehat{\boldsymbol{\Sigma}}$, and it is not possible to pre-calculate the p-value table for every possible $\widehat{\boldsymbol{\Omega}}$ and $\widehat{\boldsymbol{\Sigma}}$.

##### 3.4 Importance sampling approach

We have developed an importance sampling approach to solve this challenge. Since each analysis study with our method will have unique $\widehat{\boldsymbol{\Omega}}$ and $\widehat{\boldsymbol{\Sigma}}$, our strategy is to calculate the p-value table in a study-specific manner. The use of the standardized effect size $\boldsymbol{\eta}$ helps in this situation, because $SE(\boldsymbol{\eta})$ is independent of the SNPs. Therefore, under the null, $\boldsymbol{\eta}$ always follows $MVN\left(0, \widehat{\boldsymbol{\Sigma}}\right)$, regardless of SNPs. Thus, once we successfully build the null distribution of $S_{PLEIO}$, we can use the distribution repeatedly for all SNPs.

In importance sampling, we define the sampling distribution $p(\boldsymbol{x})$, which is a positive probability density function in $D$. Then,

$$I = \int_D f(\boldsymbol{x})q(\boldsymbol{x})dx = \int_D \frac{f(\boldsymbol{x})q(\boldsymbol{x})}{p(\boldsymbol{x})}p(\boldsymbol{x})dx = \mathrm{E^p}\left[\frac{f(\boldsymbol{X^p})q(\boldsymbol{X^p})}{p(\boldsymbol{X^p})}\right]$$

and

$$\hat{I} = \frac{1}{M}\sum_{i=1}^{M}\frac{f(\boldsymbol{X_i^p})q(\boldsymbol{X_i^p})}{p(\boldsymbol{X_i^p})},$$

where $\mathrm{E^p}[\cdot]$ denotes expectation for $\boldsymbol{X^p} \sim p$. The two practical constraints on importance sampling are; it must be feasible to generate $\boldsymbol{X^p}$, and we must be able to compute $\frac{f(\boldsymbol{x})q(\boldsymbol{x})}{p(\boldsymbol{x})}$. In importance sampling method, the variance of $\hat{I}$ can be shown as $\mathrm{Var}(\hat{I}) = \frac{\sigma_p^2}{M}$ where $\sigma_p^2$ is the standard deviation of the random variable $\int \frac{f(\boldsymbol{x})q(\boldsymbol{x})}{p(\boldsymbol{x})}$ where

$$\sigma_p^2 = \int_D \left(\frac{f(\boldsymbol{x})q(\boldsymbol{x})}{p(\boldsymbol{x})} - I\right)^2 p(\boldsymbol{x})d\boldsymbol{x}$$

$$= \int_D \frac{f(\boldsymbol{x})^2 q(\boldsymbol{x})^2}{p(\boldsymbol{x})} d\boldsymbol{x} - 2\int_D f(\boldsymbol{x})q(\boldsymbol{x})d\boldsymbol{x} + I^2$$

$$= \int_D \frac{f(\boldsymbol{x})^2 q(\boldsymbol{x})^2}{p(\boldsymbol{x})} d\boldsymbol{x} - I^2.$$

Suppose $f(\boldsymbol{x}) > 0$ and $I > 0$. In this case, the optimal $p(\boldsymbol{x})$ can be defined as $p^*(\boldsymbol{x}) = \frac{f(\boldsymbol{x})q(\boldsymbol{x})}{I}$ where this density gives $\sigma_p^2 = 0$. Since $I$ is an unknown constant that depends on the value of $\theta$, we cannot use $p^*(\boldsymbol{x})$. However, it is clear that a good sampling density of the importance sampling method will be roughly proportional to $f(\boldsymbol{x})q(\boldsymbol{x})$.

In PLEIO, it is challenging to choose a good sampling distribution $p$. In GWAS, the p-values can be as big as 1.0 or as small as $5 \times 10^{-8}$, or can be even smaller. Thus, we have a wide range of $\theta$. Depending on $\theta$, $f(\boldsymbol{x})$ changes. If we choose $p$ that resembles $f(\boldsymbol{x})q(\boldsymbol{x})$ for a large $\theta$, it can give a large variance for a small $\theta$. Reversely, if we choose $p$ that resembles $f(\boldsymbol{x})q(\boldsymbol{x})$ for a small $\theta$, it can give a large variance for a large $\theta$. To solve this challenge, we decided to use multiple sampling distributions $p_j(\boldsymbol{x})$ where $j = \{1,2, \ldots K\}$. $p_1(\boldsymbol{x})$ is $q(\boldsymbol{x})$, the probability density function of the original null distribution that follows $\mathrm{MVN}(0, \widehat{\boldsymbol{\Sigma}})$. Then we increase the variance of each coordinate by a factor of $\varphi^2$, where $\varphi \in \{1.1, 1.2, 1.3, 1.4, 1.7, 2, 2.5, 3.0, 4.0, 5.0\}$. Thus, in total, we use 11 sampling distributions such that each $p_j(\boldsymbol{x})$ follows $\mathrm{MVN}(\mathbf{0}, \mathrm{c}_\mathrm{j}^2 \widehat{\boldsymbol{\Sigma}})$, where $c_j$ is a constant value ranged from 1 to 5. Let $\boldsymbol{X}^{\boldsymbol{p}_j}$ denote a matrix representing samples generated from the $j$th sampling distribution, and let $\alpha_j$ denote the proportion of samples generated from the $j$th sampling distribution. Then, the matrix $\boldsymbol{X}^{\boldsymbol{p}_j}$ has a size of $\alpha_j M \times T$. In PLEIO, we generate $\boldsymbol{X}^{\boldsymbol{p}}$ from each $p_j(\boldsymbol{x})$ uniformly, so that $\alpha_j \approx \frac{1}{K}$. In the importance sampling method using multiple sampling distributions, the p-value of a given $\theta$ can be estimated by follows:

$$\hat{I} = \frac{1}{M}\sum_{i=1}^{M}\frac{f(\boldsymbol{X_i^p})q(\boldsymbol{X_i^p})}{p_\alpha(\boldsymbol{X_i^p})},$$

where $p_\alpha(\boldsymbol{X_i^p}) = \sum_{j=1}^{K}\alpha_j p_j(\boldsymbol{X_i^p})$, and $\sum_{j=1}^{K}\alpha_j = 1$.

Recently, Owen and Zhou (2000) [8] proposed a novel importance sampling approach to minimize variance of the estimate in the situation where multiple sampling distributions are used. We employed this approach. The method generates $\boldsymbol{X^p}$ from $p_j(\boldsymbol{x})$ and uses $p_j(\boldsymbol{X^p})/p_\alpha(\boldsymbol{X^p})$ as the control variates of $f(\boldsymbol{X^p})q(\boldsymbol{X^p})/p_\alpha(\boldsymbol{X^p})$. Assuming high correlations between $f(\boldsymbol{x})q(\boldsymbol{x})$ and $p_j(\boldsymbol{x})$, we expect a large reduction in variance when estimating $\mathrm{E}\big(f(\boldsymbol{x})\big)$ using the control variate method. Note that $p_j(\boldsymbol{x})$ is a probability density function and therefore has the expected value $\int_D p_j(\boldsymbol{x})d\boldsymbol{x} = 1$. The expression of the importance sampling with the control variate method can be shown as follows:

$$\hat{I} = \frac{1}{M}\left(\sum_{i=1}^{M}\frac{f(\boldsymbol{X_i^p})q(\boldsymbol{X_i^p}) - \sum_{j=1}^{K}\beta_j p_j(\boldsymbol{X_i^p})}{p_\alpha(\boldsymbol{X_i^p})}\right) + \sum_{k=1}^{K}\beta_k\mu_{p_k}$$

where $\mu_{p_j} = \mathrm{E}^p[p_j(\boldsymbol{X_i^p})/p_\alpha(\boldsymbol{X_i^p})] = \int p_j(\boldsymbol{x})d\boldsymbol{x} = 1$ for any $j$. Define $m^* = \frac{f(\boldsymbol{x})q(\boldsymbol{x}) - \sum_{j=1}^{K}\beta_j p_j(\boldsymbol{x})}{p_\alpha(\boldsymbol{x})} + \sum_{k=1}^{K}\beta_k\mu_{p_k}$ such that $\mathrm{E}[m^*] = I$. The variance of $m^*$ is then,

$$\begin{aligned}
\mathrm{Var}(m^*) = &-\beta_1\mathrm{Cov}\left(\frac{f(\boldsymbol{x})q(\boldsymbol{x})}{p_\alpha(\boldsymbol{x})}, \frac{p_1(\boldsymbol{x})}{p_\alpha(\boldsymbol{x})}\right) + \sum_{l=1}^{K}\beta_1\beta_l\mathrm{Cov}\left(\frac{p_1(\boldsymbol{x})}{p_\alpha(\boldsymbol{x})}, \frac{p_l(\boldsymbol{x})}{p_\alpha(\boldsymbol{x})}\right) \\
&-\beta_2\mathrm{Cov}\left(\frac{f(\boldsymbol{x})q(\boldsymbol{x})}{p_\alpha(\boldsymbol{x})}, \frac{p_2(\boldsymbol{x})}{p_\alpha(\boldsymbol{x})}\right) + \sum_{l=1}^{K}\beta_2\beta_l\mathrm{Cov}\left(\frac{p_2(\boldsymbol{x})}{p_\alpha(\boldsymbol{x})}, \frac{p_l(\boldsymbol{x})}{p_\alpha(\boldsymbol{x})}\right) \\
&-\beta_3\mathrm{Cov}\left(\frac{f(\boldsymbol{x})q(\boldsymbol{x})}{p_\alpha(\boldsymbol{x})}, \frac{p_3(\boldsymbol{x})}{p_\alpha(\boldsymbol{x})}\right) + \sum_{l=1}^{K}\beta_3\beta_l\mathrm{Cov}\left(\frac{p_3(\boldsymbol{x})}{p_\alpha(\boldsymbol{x})}, \frac{p_l(\boldsymbol{x})}{p_\alpha(\boldsymbol{x})}\right) \\
&\vdots \\
&-\beta_K\mathrm{Cov}\left(\frac{f(\boldsymbol{x})q(\boldsymbol{x})}{p_\alpha(\boldsymbol{x})}, \frac{p_K(\boldsymbol{x})}{p_\alpha(\boldsymbol{x})}\right) + \sum_{l=1}^{K}\beta_K\beta_l\mathrm{Cov}\left(\frac{p_K(\boldsymbol{x})}{p_\alpha(\boldsymbol{x})}, \frac{p_l(\boldsymbol{x})}{p_\alpha(\boldsymbol{x})}\right) + \mathrm{Var}\left(\frac{f(\boldsymbol{x})q(\boldsymbol{x})}{p_\alpha(\boldsymbol{x})}\right)
\end{aligned}$$

because $\text{Var}(\beta_k \mu_{p_k}) = 0$. By definition, the optimal $\boldsymbol{\beta}$ can be found by solving the following partial derivatives.

$$\frac{\partial \text{Var}(m^*)}{\partial \beta_j} = -\text{Cov}\left(\frac{f(\boldsymbol{x})q(\boldsymbol{x})}{p_\alpha(\boldsymbol{x})}, \frac{p_j(\boldsymbol{x})}{p_\alpha(\boldsymbol{x})}\right) + \sum_{l=1}^{K} \beta_l \text{Cov}\left(\frac{p_j(\boldsymbol{x})}{p_\alpha(\boldsymbol{x})}, \frac{p_l(\boldsymbol{x})}{p_\alpha(\boldsymbol{x})}\right)$$

which generates $K(=11)$ equations and $K$ unknown variables ($\boldsymbol{\beta}$) as follows:

$$\begin{aligned}
\frac{\partial \text{Var}(m^*)}{\partial \beta_1} &= -\text{Cov}\left(\frac{f(\boldsymbol{x})q(\boldsymbol{x})}{p_\alpha(\boldsymbol{x})}, \frac{p_1(\boldsymbol{x})}{p_\alpha(\boldsymbol{x})}\right) + \sum_{l=1}^{K} \beta_l \text{Cov}\left(\frac{p_1(\boldsymbol{x})}{p_\alpha(\boldsymbol{x})}, \frac{p_l(\boldsymbol{x})}{p_\alpha(\boldsymbol{x})}\right) \\
\frac{\partial \text{Var}(m^*)}{\partial \beta_2} &= -\text{Cov}\left(\frac{f(\boldsymbol{x})q(\boldsymbol{x})}{p_\alpha(\boldsymbol{x})}, \frac{p_2(\boldsymbol{x})}{p_\alpha(\boldsymbol{x})}\right) + \sum_{l=1}^{K} \beta_l \text{Cov}\left(\frac{p_2(\boldsymbol{x})}{p_\alpha(\boldsymbol{x})}, \frac{p_l(\boldsymbol{x})}{p_\alpha(\boldsymbol{x})}\right) \\
&\vdots \\
\frac{\partial \text{Var}(m^*)}{\partial \beta_K} &= -\text{Cov}\left(\frac{f(\boldsymbol{x})q(\boldsymbol{x})}{p_\alpha(\boldsymbol{x})}, \frac{p_K(\boldsymbol{x})}{p_\alpha(\boldsymbol{x})}\right) + \sum_{l=1}^{K} \beta_l \text{Cov}\left(\frac{p_K(\boldsymbol{x})}{p_\alpha(\boldsymbol{x})}, \frac{p_l(\boldsymbol{x})}{p_\alpha(\boldsymbol{x})}\right).
\end{aligned}$$

Here, the function of $\text{Var}(m^*)$ is a quadratic function for $\beta_j$. Therefore, the $\boldsymbol{\beta}^*$ which maximizes the variance of $\text{Var}(m^*)$ satisfies $\frac{\partial \text{Var}(m^*)}{\partial \beta_j^*} = 0$, the root of the partial derivative. Thus, the optimal $\boldsymbol{\beta}^*$ $\left(\boldsymbol{\beta} \middle| \frac{\partial \text{Var}(m^*)}{\partial \beta_j} = 0 \text{ and } \boldsymbol{X}_{\boldsymbol{i}}^{\boldsymbol{p}}\right)$ can be obtained by solving the linear equation:

$$\begin{aligned}
\text{Var}(P_1)\beta_1^* + \text{Cov}(P_1, P_2)\beta_2^* + \cdots + \text{Cov}(P_1, P_K)\beta_K^* &= \text{Cov}(FQ\ , P_1) \\
\text{Cov}(P_2, P_1)\beta_1^* + \text{Var}(P_2)\beta_2^* + \cdots + \text{Cov}(P_2, P_K)\beta_K^* &= \text{Cov}(FQ\ , P_2) \\
&\vdots \\
\text{Cov}(P_K, P_1)\beta_1^* + \text{Cov}(P_K, P_2)\beta_2^* + \cdots + \text{Var}(P_K)\beta_K^* &= \text{Cov}(FQ\ , P_K)
\end{aligned}$$

which can be shown as follows:

$$\begin{bmatrix} \text{Var}(P_1) & \text{Cov}(P_1, P_2) & \dots & \text{Cov}(P_1, P_K) \\ \text{Cov}(P_2, P_1) & \text{Var}(P_2) & \cdots & \text{Cov}(P_2, P_K) \\ \vdots & \vdots & \ddots & \vdots \\ \text{Cov}(P_K, P_1) & \text{Cov}(P_K, P_2) & \dots & \text{Var}(P_K) \end{bmatrix} \begin{bmatrix} \beta_1^* \\ \beta_2^* \\ \vdots \\ \beta_K^* \end{bmatrix} = \begin{bmatrix} \text{Cov}(FQ\ , P_1) \\ \text{Cov}(FQ\ , P_2) \\ \vdots \\ \text{Cov}(FQ\ , P_K) \end{bmatrix}$$

and

$$\begin{bmatrix} \beta_1^* \\ \beta_2^* \\ \vdots \\ \beta_K^* \end{bmatrix} = \begin{bmatrix} \mathrm{Var}(P_1) & \mathrm{Cov}(P_1, P_2) & \dots & \mathrm{Cov}(P_1, P_K) \\ \mathrm{Cov}(P_2, P_1) & \mathrm{Var}(P_2) & \cdots & \mathrm{Cov}(P_2, P_K) \\ \vdots & \vdots & \ddots & \vdots \\ \mathrm{Cov}(P_K, P_1) & \mathrm{Cov}(P_K, P_2) & \dots & \mathrm{Var}(P_K) \end{bmatrix}^{-1} \begin{bmatrix} \mathrm{Cov}(FQ\ , P_1) \\ \mathrm{Cov}(FQ\ , P_2) \\ \vdots \\ \mathrm{Cov}(FQ\ , P_K) \end{bmatrix},$$

where $P_l = \frac{p_j(\boldsymbol{X_i^p})}{p_\alpha(\boldsymbol{X_i^p})}$, and $FQ = \frac{f(\boldsymbol{X_i^p})q(\boldsymbol{X_i^p})}{p_\alpha(\boldsymbol{X_i^p})}$. Owen and Zhou[8] showed that If at least one of $p_j(\boldsymbol{x}) > 0$ whenever $f(\boldsymbol{x})q(\boldsymbol{x}) > 0$, then $\hat{I}_{\alpha,\beta}$ is unbiased and $\mathrm{Var}(\hat{I}_{\alpha,\beta}) \le \mathrm{Var}\left(\hat{I}_{\alpha_j p_j}\right)$ for any $j$ where $\hat{I}_{\alpha_j p_j} = \frac{1}{n_j}\sum_{i=1}^{n_j} \frac{f(\boldsymbol{X_i^p})q(\boldsymbol{X_i^p})}{p_j(\boldsymbol{X_i^p})}$.

##### 3.5 Implementation

The implementation of the p-value estimation in PLEIO is as follows. After calculating $\widehat{\boldsymbol{\Omega}}$ and $\widehat{\boldsymbol{\Sigma}}$ using LDSC, we assume that these values are true values and generate the null samples by using importance sampling method. The default number of sampling is 100K, where each of 11 distributions being used equally. For each sample, we use our efficient transformation technique for the Newton Raphson method to determine the maximum likelihood estimate $\hat{\tau}^2$ and calculate $S_{PLEIO}$. Then, we calculate p-values of 50 different $\theta$ that are in the range (0, 40). For each $\theta$, we calculate the optimal $\beta$ for the control variate method and use the method to calculate the p-value from our null samples. Using these 50 points, we interpolate p-values for $\theta < 40$ using B-spline fitting and extrapolate p-values for $\theta > 40$ using linear fitting on the log p-value scale.

### 4 Simulations

#### 4.1 FPR simulation

We elaborate how we generated SNP effects in the FPR simulation. In each simulation of the given environmental correlation $\mathbf{C_e}$, we simulated observed effect sizes of $L$ studies from $\hat{\boldsymbol{\eta}}_{\boldsymbol{sim}} \sim \text{MVN}(\mathbf{0}, \boldsymbol{\Sigma}_{\boldsymbol{sim}})$ where $\boldsymbol{\Sigma}_{\mathbf{sim}} = \mathbf{C}_e$. We assumed a significance threshold level $\alpha$ and estimated FPR at $\alpha$ as the proportion of simulations in which the p-value was $\leq \alpha$.

#### 4.2 Power simulation

We elaborate how we generated SNP effects in the power simulation. We assumed a SNP whose minor allele frequency is 0.3 ($p = 0.3$) under the Hardy Weinberg equilibrium. For $L$ traits, we assumed specific non-zero heritabilities and complex genetic correlations between those traits. For the sake of simplicity, we assumed no environmental correlations between these traits ($\mathbf{C_e}$ is a diagonal matrix). Each trait is either binary or quantitative, and we assumed that the phenotypic units of the quantitative traits may vary. After determining the type and the unit of the phenotype for each trait, we simulated observed effect sizes of the SNP as follows. Let $\mathbf{h^2} = (h_1^2, h_2^2, \ldots)$ be an $L \times 1$ vector representing the heritabilities of the $L$ traits, and $\mathbf{C_g}$ be an $L \times L$ matrix representing the genetic correlation.

First, we explain how we generated effect sizes for quantitative traits ($\mathbf{Q}$). Let $\boldsymbol{\beta_i} = (\beta_{i1}, \beta_{i2}, \ldots)$ be the $L \times 1$ vector of true effect size of the SNP $i$. To simulate observed effect sizes, we first sample the true effect size $\boldsymbol{\beta_i} \sim \text{MVN}\left(0, \frac{1}{M_{true}} \boldsymbol{\Omega}_{\mathbf{sim}}\right)$ where $\boldsymbol{\Omega}_{\mathbf{sim}} = \text{diag}\left(\sqrt{\mathbf{h^2}}\right) \cdot \mathbf{C_g} \cdot \text{diag}\left(\sqrt{\mathbf{h^2}}\right)$. $M_{true}$ denotes the number of causal variants, which is set to 1000 in our simulation. Let $N$ denote the number of samples, and $h_t^2$ denotes the $t$th element of the $\mathbf{h^2}$. For each trait, we simulated the $N \times 1$ vector of genotypes $\boldsymbol{x_t} = (x_{t1}, x_{t2}, \ldots, x_{tN})$ from $x_{tj} \sim Binomial(2, p)$ and standardized $\boldsymbol{x_t}$ so that $\boldsymbol{x_{t,std}} = \frac{\boldsymbol{x_t} - 2p}{2p(1-p)}$. We then generated $\boldsymbol{y}$, the $N \times 1$ vector of phenotypes, from the linear model $\boldsymbol{y_t} = \beta_{it} \boldsymbol{x_{t,std}} + \boldsymbol{\epsilon_t}$ where

$\boldsymbol{\epsilon_t} = (\epsilon_{t1}, \epsilon_{t2}, \dots, \epsilon_{tN})$ is a $N \times 1$ vector representing the error such that $\epsilon_{tj} \sim \mathrm{N}\left(0, 1 - \frac{1}{M_{true}}\ h_t^2\right)$. We obtained the observed effect size $\left(\hat{\beta}_{it}\right)$ from the linear regression using $\boldsymbol{x_{t,std}}$ as the regressor and $\boldsymbol{y_t}$ as the regressand.

Second, we explain how we generated effect sizes for binary traits (**B**). Because this simulation is slightly more complicated, we describe the procedure in four steps. We assumed the study design with half cases (N/2) and half controls (N/2).

**[Step 1: Sampling of true effect size under observed linear scale]**

We sample the true effect size $\boldsymbol{\beta_i} \sim \mathrm{MVN}\left(0, \frac{1}{M_{true}} \boldsymbol{\Omega_{sim}}\right)$ where $\boldsymbol{\Omega_{sim}} = \mathrm{diag}\left(\sqrt{\mathbf{h^2}}\right) \cdot \mathbf{C_g} \cdot \mathrm{diag}\left(\sqrt{\mathbf{h^2}}\right)$. Note that $\boldsymbol{\beta_i}$ is the true effect size under the liability scale. Next, we convert this true effect size into the observed scale $\boldsymbol{\beta_{i,obs}} = \boldsymbol{\beta_i}/\delta_t$. The scaling factor $\delta_t$ is $\sqrt{\frac{K_t^2(1-K_t)^2}{P_t(1-P_t)} \cdot \frac{1}{\psi(\phi^{-1}(1-K_t))}}$, where $K_t$ refers to the disease prevalence of trait $t$, $P_t$ refers to the sample prevalence (0.5 in our balanced design), $\psi$ refers to the probability density function of the standard normal distribution, and $\phi^{-1}$ refers to the inverse of the cumulative density function of the standard normal distribution.

**[Step 2: Searching for relative risk corresponding to the true effect size]**

Now, we search for the relative risk that would give us $\boldsymbol{\beta_{i,obs}}$ under the observed linear scale. This can be done as follows. We assume a specific relative risk $\gamma$. Given the disease prevalence $K_t$ , the expected case minor allele frequency (MAF) is $p^+ = \frac{\gamma p}{p(\gamma-1)+1}$ and the expected control MAF is $p^- = \frac{p - p^+ K_t}{1-K_t}$ where $p = 0.3$ is the population MAF. Under the Hardy-Weinberg equilibrium, we can construct a reference genotype $\overline{\boldsymbol{x_t}}$ such that the case MAF is exactly $p^+$ and control MAF is exactly $p^-$ (after ignoring the integer rounding). Given this $\overline{\boldsymbol{x_t}}$ and the phenotype $\boldsymbol{y_t}$ consisting of half ones and half zeros, we can perform linear regression to get the effect size under the observed linear scale $\overline{\boldsymbol{\beta_{i,obs}}}$ (after standardizing $\overline{\boldsymbol{x_t}}$). Finally, we search the relative risk $\gamma$ that gives us $\boldsymbol{\beta_{i,obs}} = \overline{\boldsymbol{\beta_{i,obs}}}$.

**[Step 3: Generating genotypes]**

Now that we know the relative risk (and thereby the expected case MAF and control MAF) that corresponds to $\boldsymbol{\beta_{i,obs}}$, we can generate simulated genotypes $\boldsymbol{x_t}$. We generate the case genotypes from $x_{tj} \sim Binomial(2, p^{+})$ and the control genotypes from $x_{tj} \sim Binomial(2, p^{-})$.

**[Step 4: Logistic regression]**

It is common that the heritability calculations of binary traits are based on the (observed and liability scale) linear model. This was why we had to derive the relative risk and the case and control MAFs through the observed scale linear model. However, in association analyses, the logistic regression model is commonly used. To simulate a realistic situation, we applied logistic regression to $\boldsymbol{y_t}$ and the sampled $\boldsymbol{x_t}$. This way, we obtained the log odds ratios and the standard errors for $L$ traits. This information was used as the final input for binary traits in our power analysis.
