## Supplementaryu Figure 1-8 and Supplementary Table 1-3 for "A method to map and interpret pleiotropic loci using summary statistics of multiple traits"

### 1 Supplementary figures

**Supplementary Figure 1.** A toy example designed to understand the association analysis carried out by PLEIO.

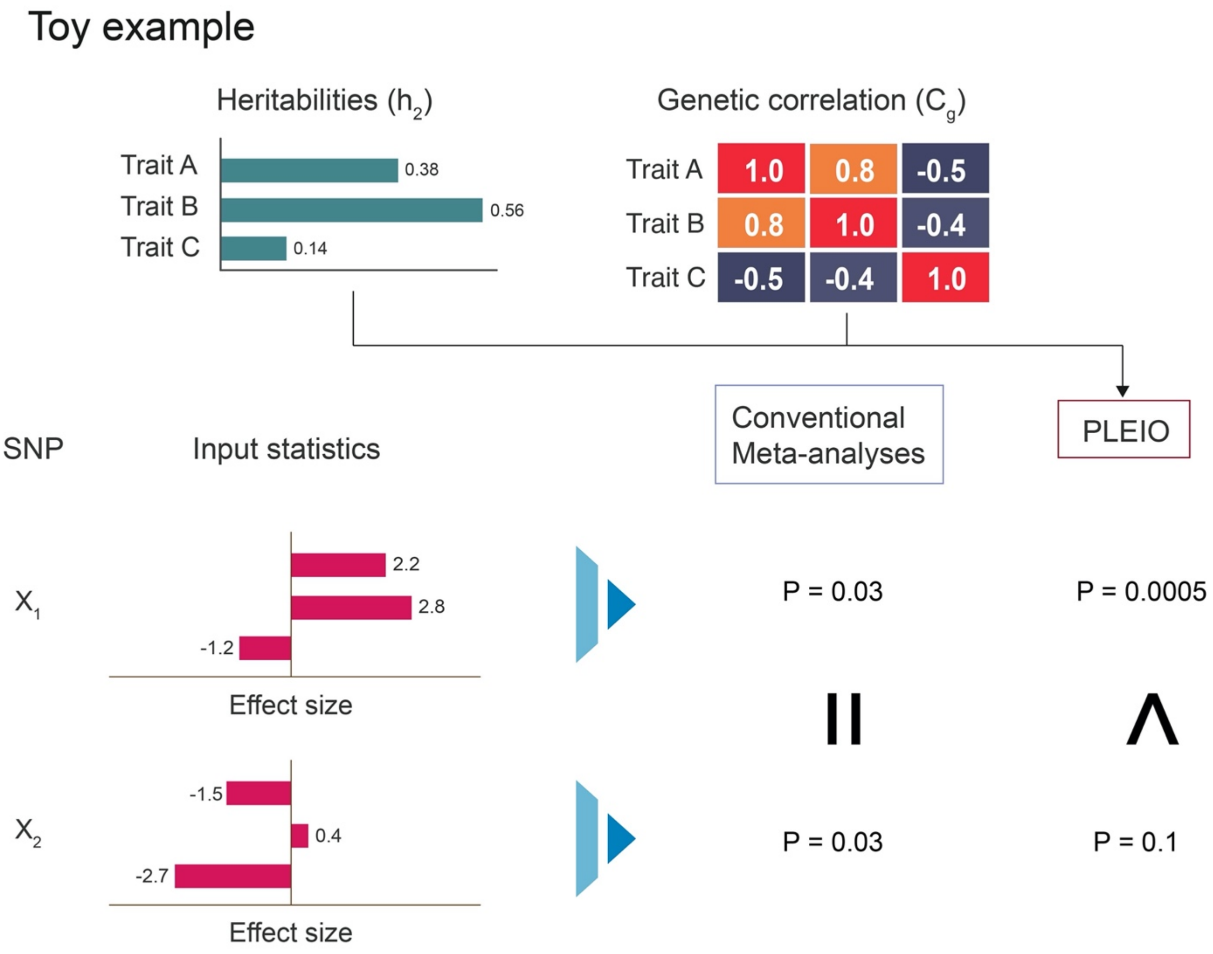

**Supplementary Figure 2.** Power test result under the combined condition of various sources of heterogeneity. (1) the heritabilities of the traits differ. (2) The genetic correlation structure among traits is complex, having both negative and positive correlations. (3) The traits are a mixture of quantitative and binary traits where phenotypic units differ among traits.

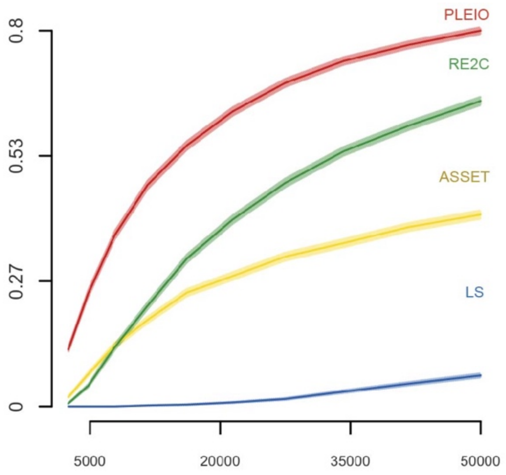


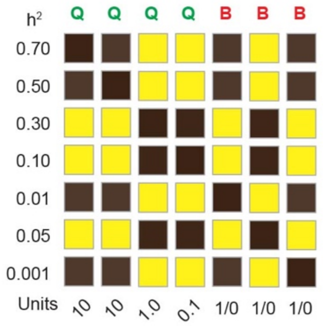

**Supplementary Figure 3. Genetic correlation and environmental correlation among 18 traits.** The 18 × 18 matrix shows the genetic (upper triangular) and environmental (lower triangular) correlations among 18 traits. The labels on the left and the top are the names of the traits, and the labels on the right indicate the heritabilities along with the names of database and the types of the phenotypes (green: binary, red: quantitative).

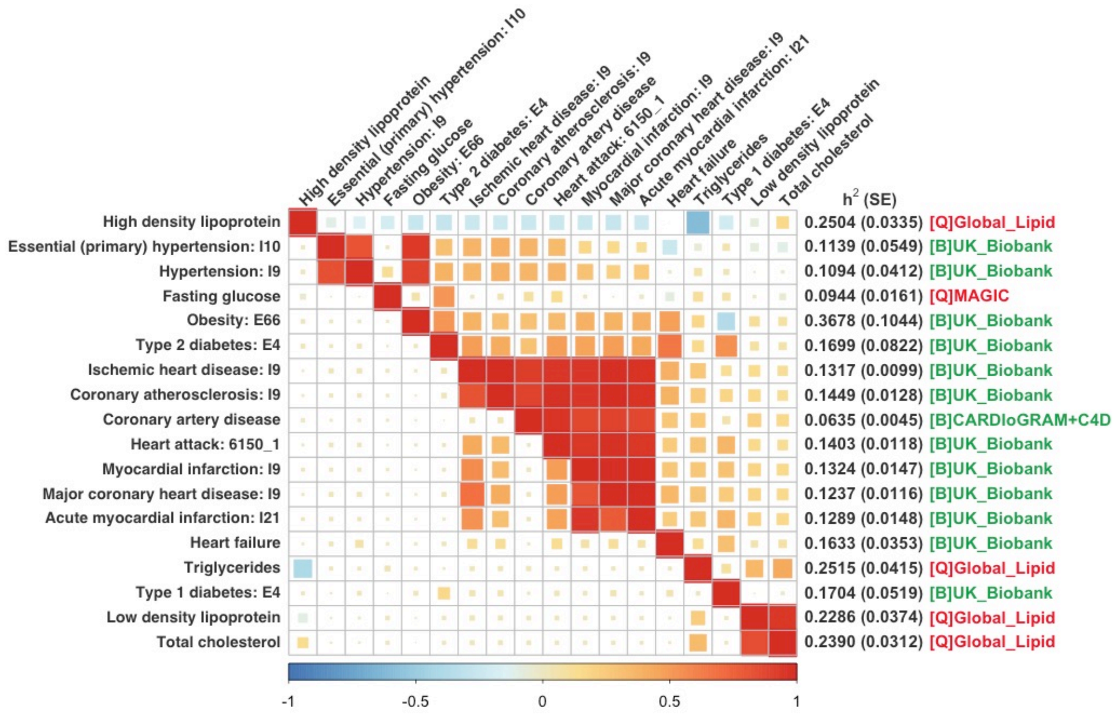

**Supplementary Figure 4.** Local Manhattan plots of the 20 novel variants identified using PLEIO in real data analysis of 18 CVD-related traits.

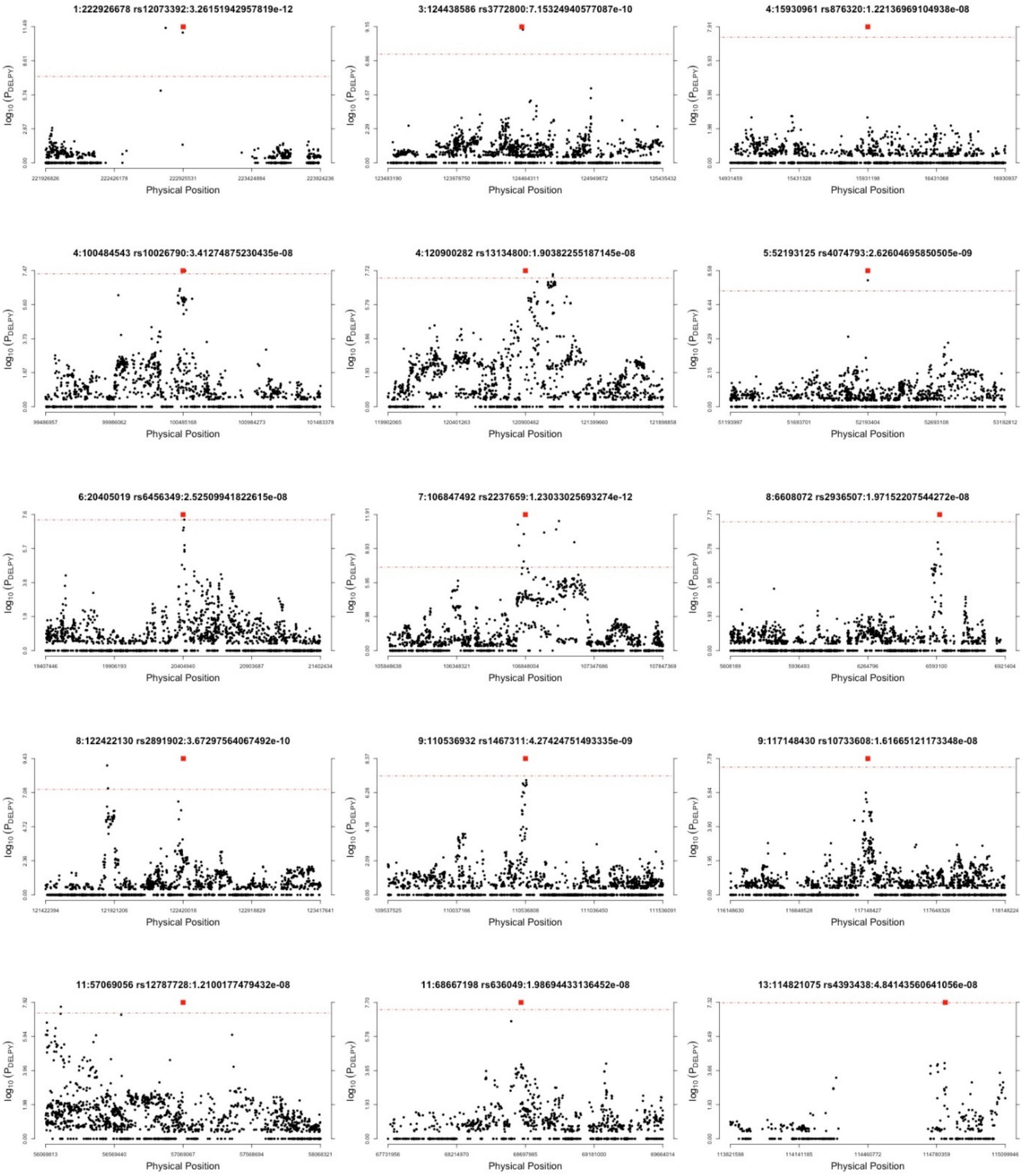

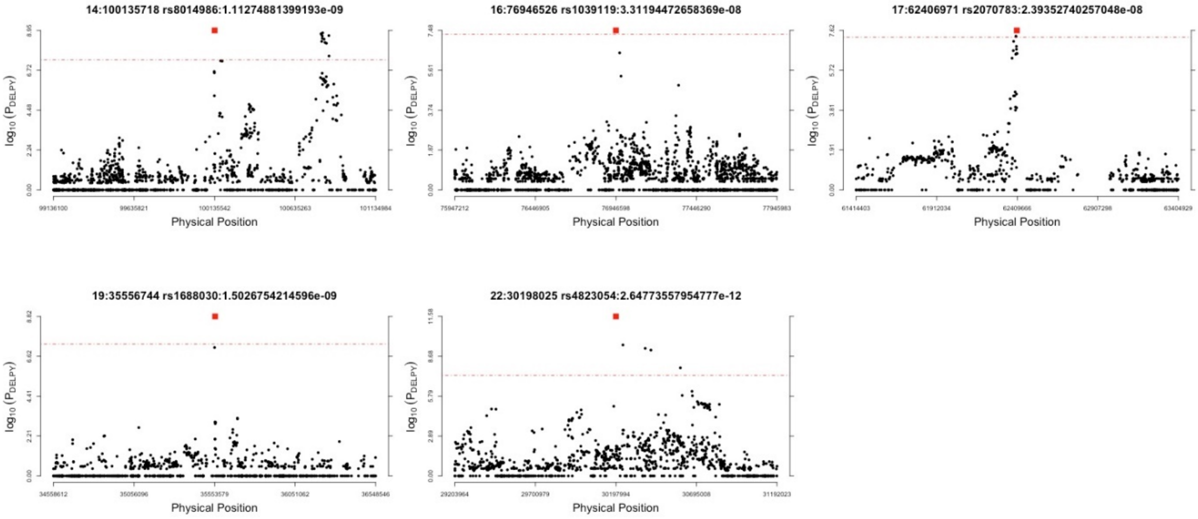
14:100135718 rs8014986:1.11274881399193e-09
16:76946526 rs1039119:3.31194472658369e-08
17:62406971 rs2070783:2.39352740257048e-08
19:35556744 rs1688030:1.5026754214596e-09
22:30198025 rs4823054:2.64773557954777e-12
$\log_{10}(P_{DELPY})$
Physical Position

**Supplementary Figure 5.** Mirrored Manhattan plot of association analysis of 18 CVD-related traits. The upper side shows the PLEIO's p-values, and the bottom side shows the smallest p-values among the 18 individual GWASs.

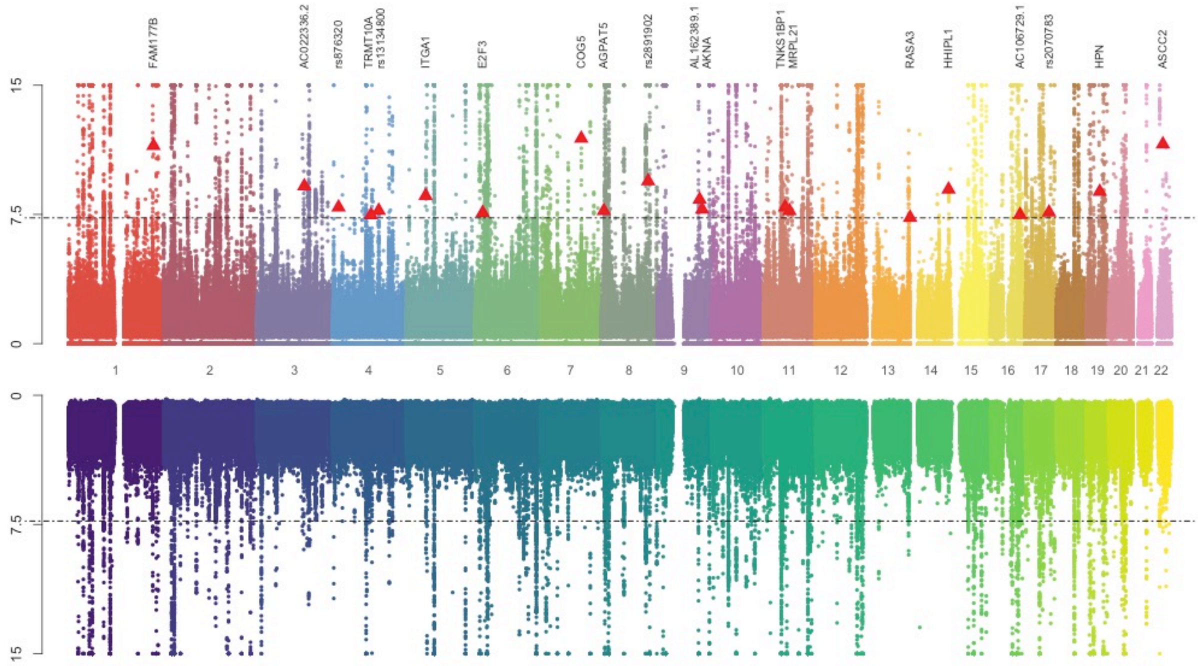

Supplementary Figure 6 A. Pleiotropy plot for rs12073392.

[Group 1: Driven by seven binary traits]

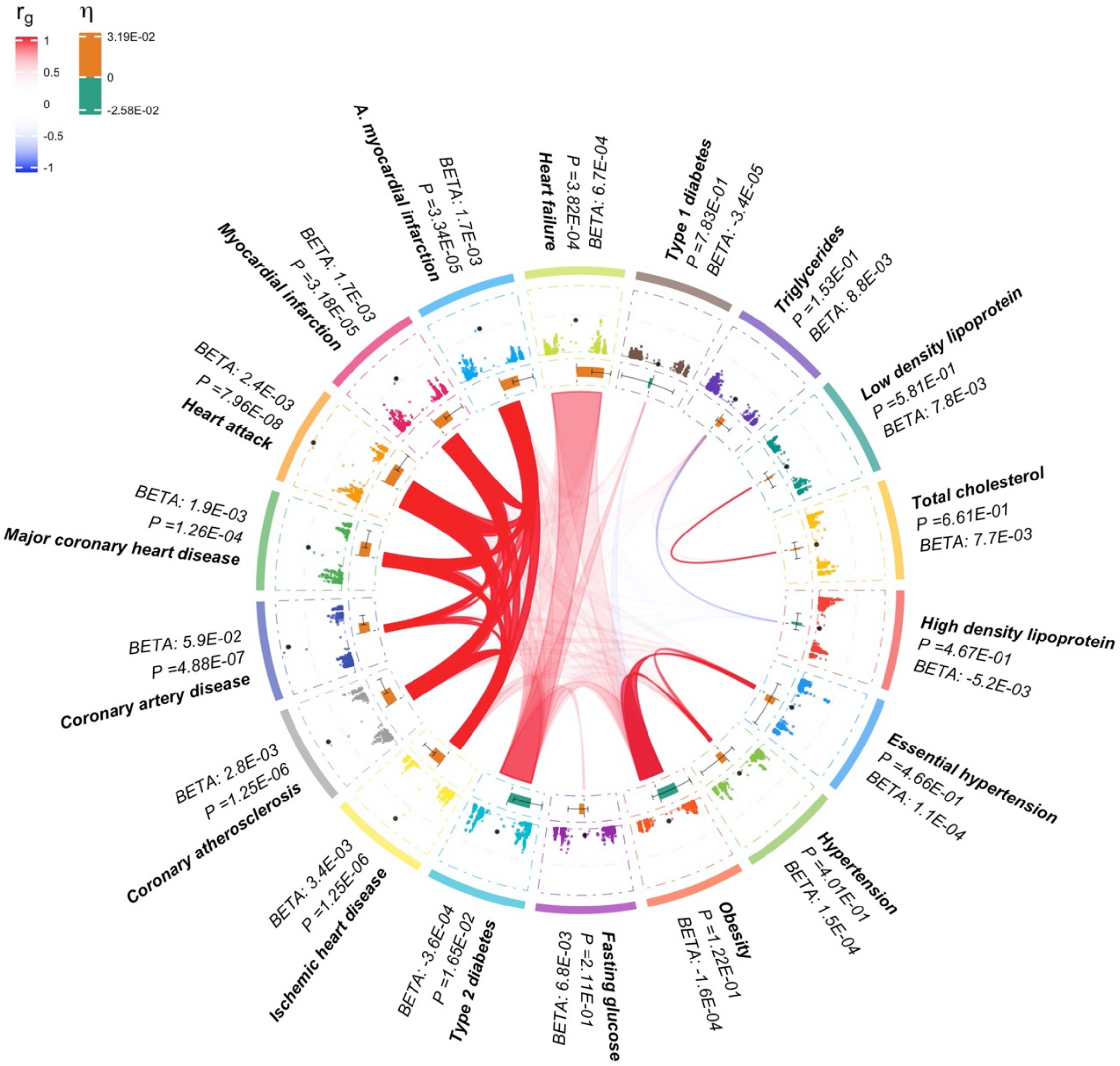

Supplementary Figure 6 B. Pleiotropy plot for rs3772800

[Group 1: Driven by seven binary traits]

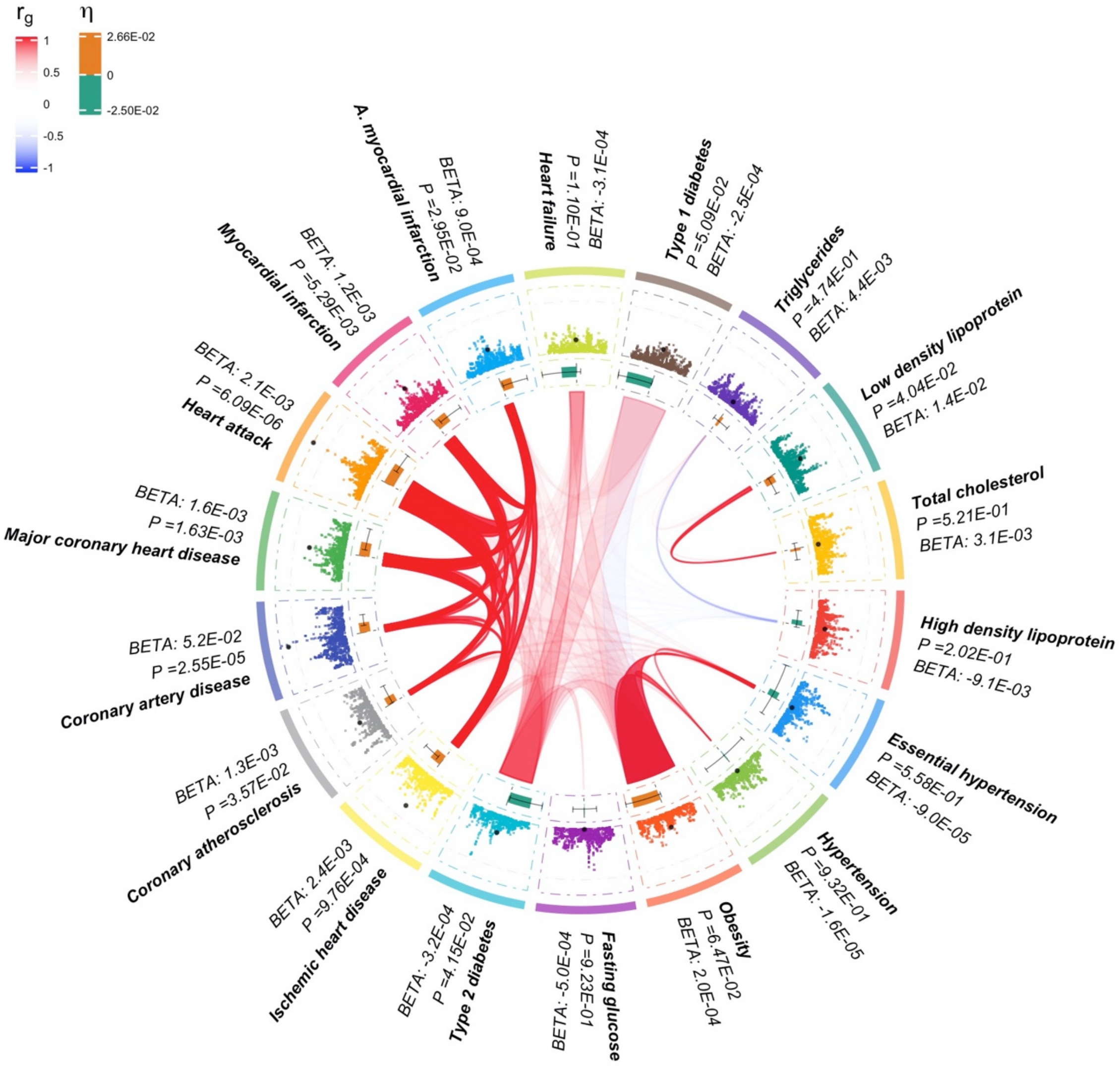

Supplementary Figure 6 C. Pleiotropy plot for rs13134800.

[Group 1: Driven by seven binary traits]

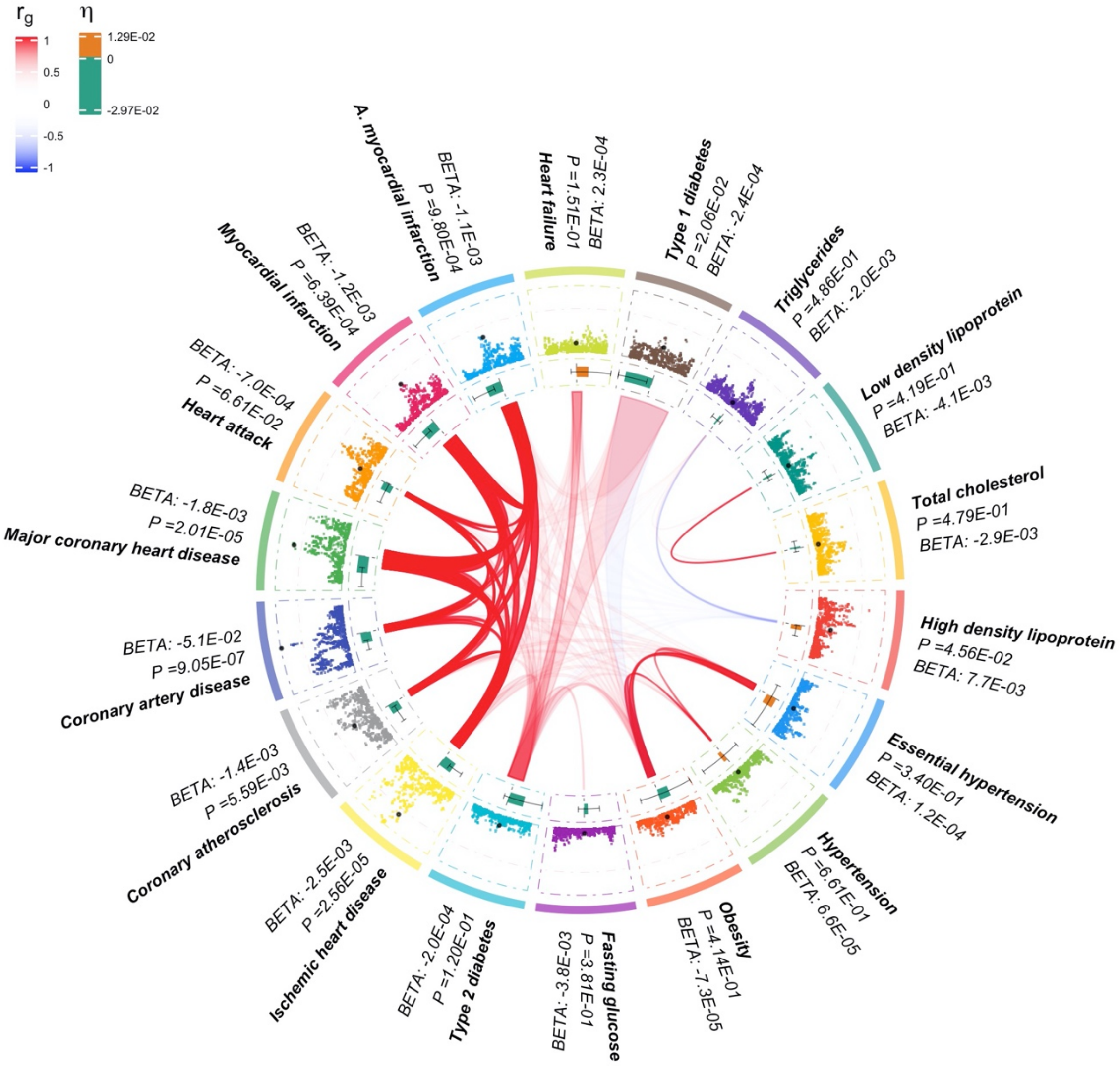

Supplementary Figure 6 D. Pleiotropy plot for rs1467311

[Group 1: Driven by seven binary traits]

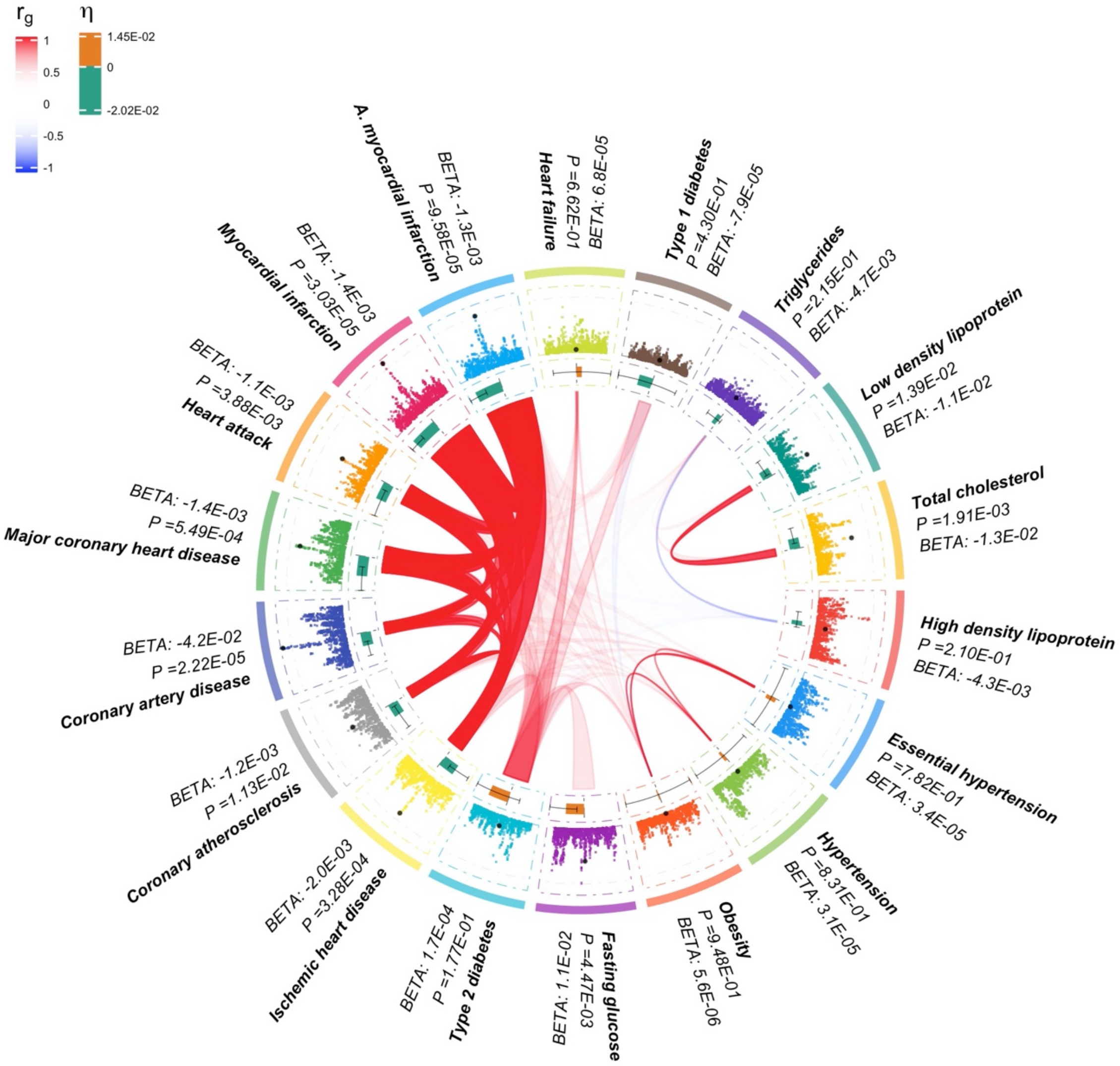

Supplementary Figure 6 E. Pleiotropy plot for rs8014986.

[Group 1: Driven by seven CVD binary traits]

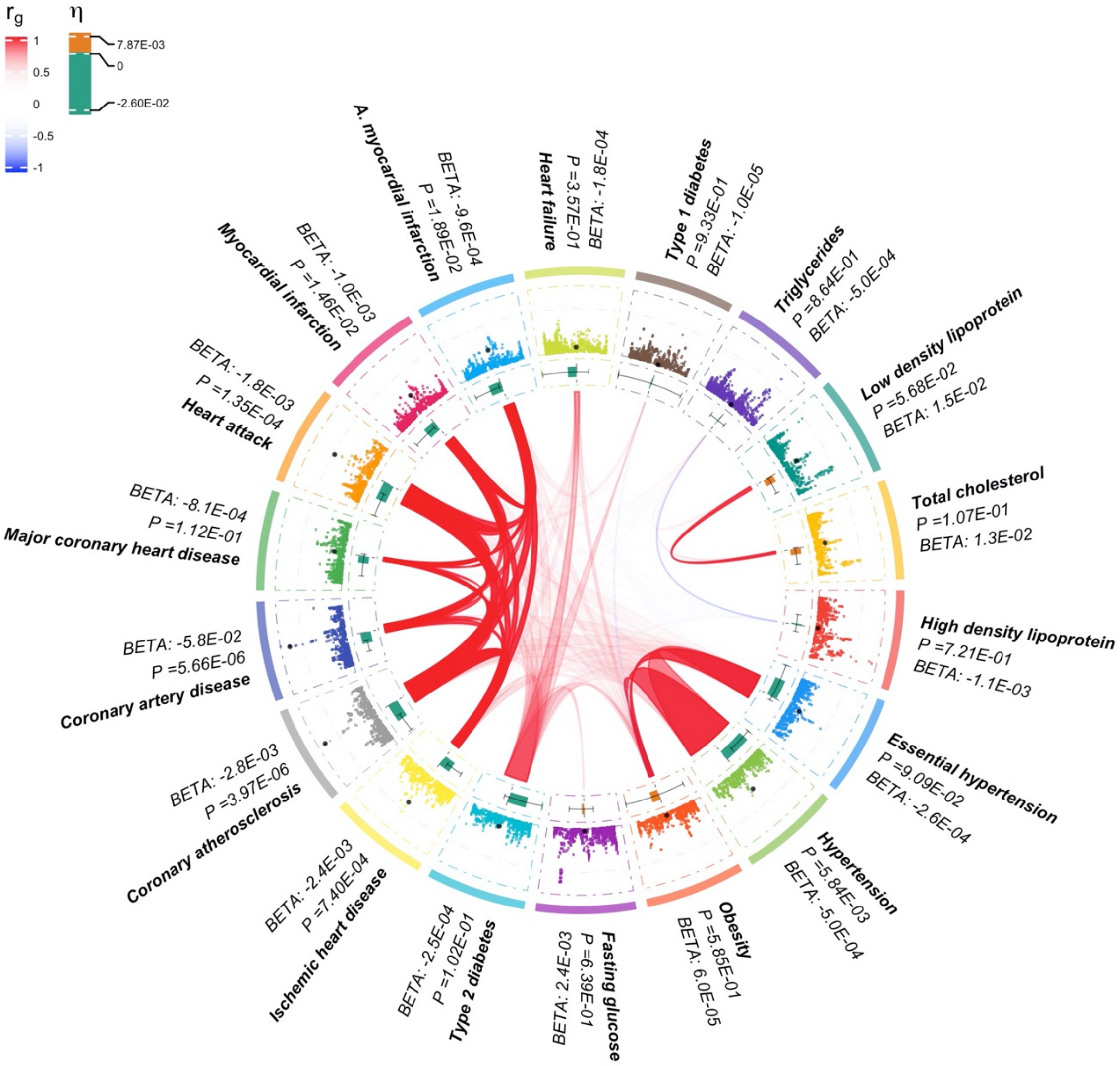

Supplementary Figure 6 F. Pleiotropy plot for rs2070783

[Group 1: Driven by seven CVD binary traits]

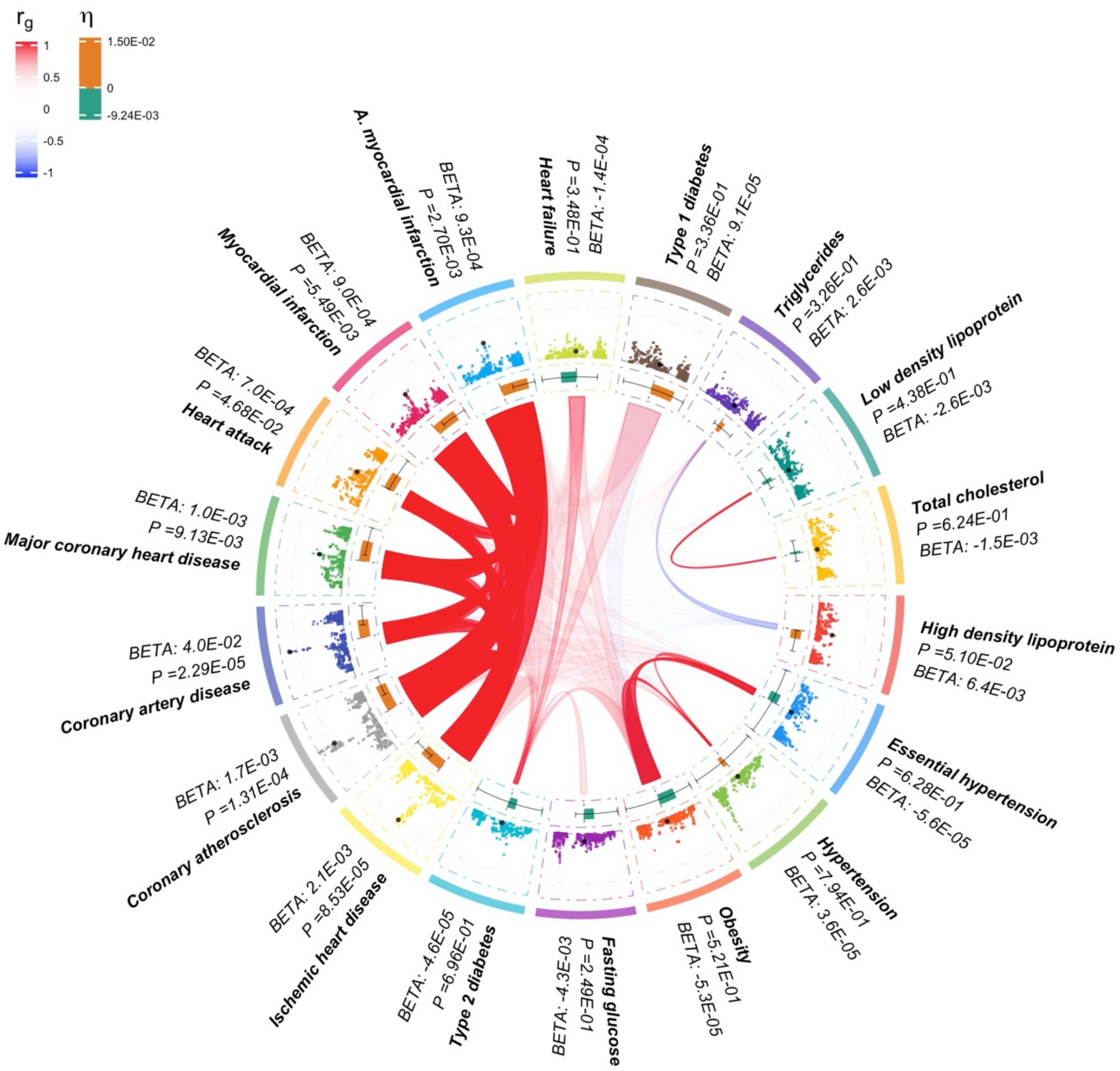

Supplementary Figure 6 G. Pleiotropy plot for rs6456349.

[Group 2: Driven by lipid phenotypes (triglycerides, LDL, HDL, and total cholesterol)]

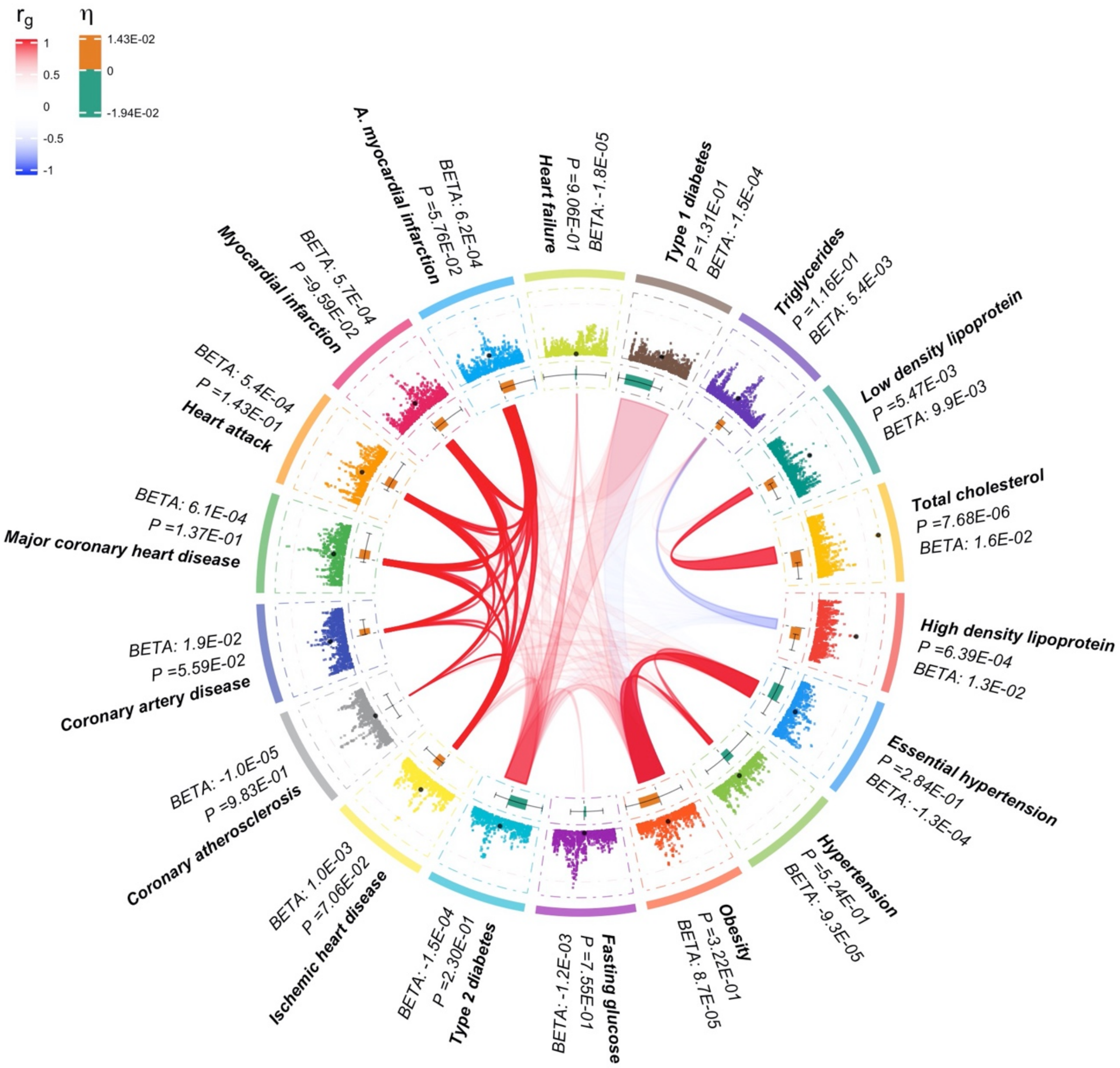

Supplementary Figure 6 H. Pleiotropy plot for rs2936507.

[Group 2: Driven by lipid phenotypes (triglycerides, LDL, HDL, and total cholesterol)]

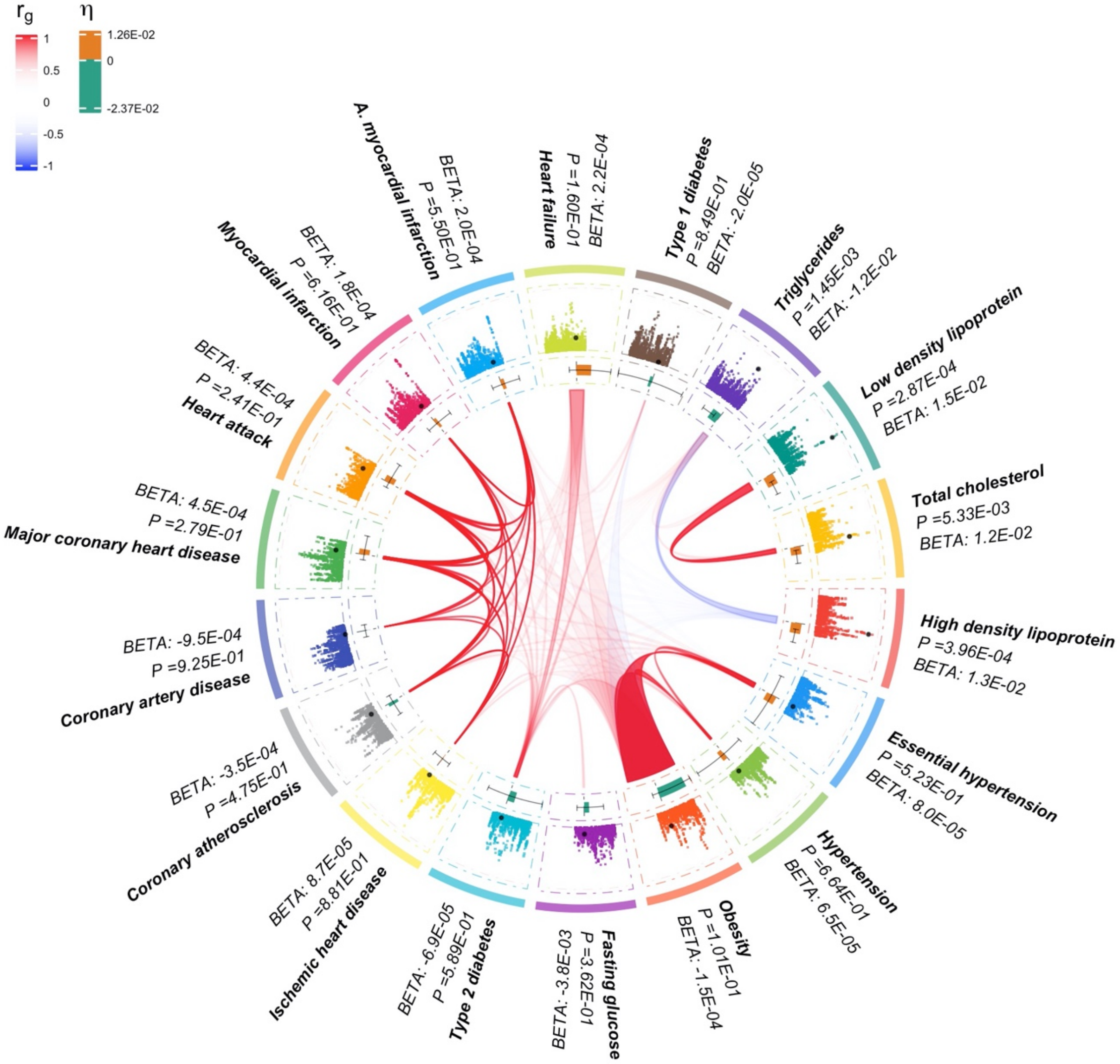

Supplementary Figure 6 I. Pleiotropy plot for rs636049.

[Group 2: Driven by lipid phenotypes (triglycerides, LDL, HDL, and total cholesterol)]

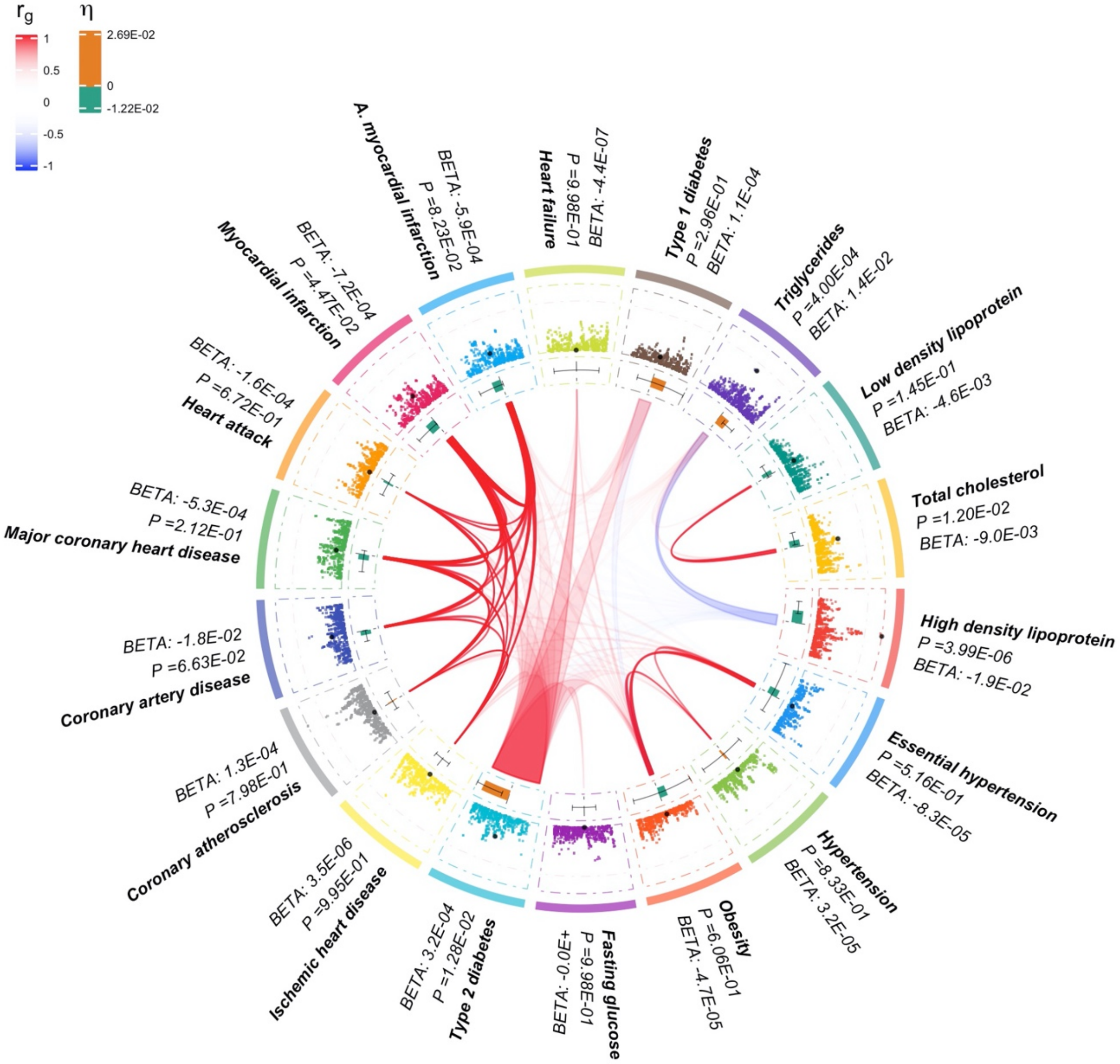

Supplementary Figure 6 J. Pleiotropy plot for rs1688030.

[Group 2: Driven by lipid phenotypes (triglycerides, LDL, HDL, and total cholesterol)]

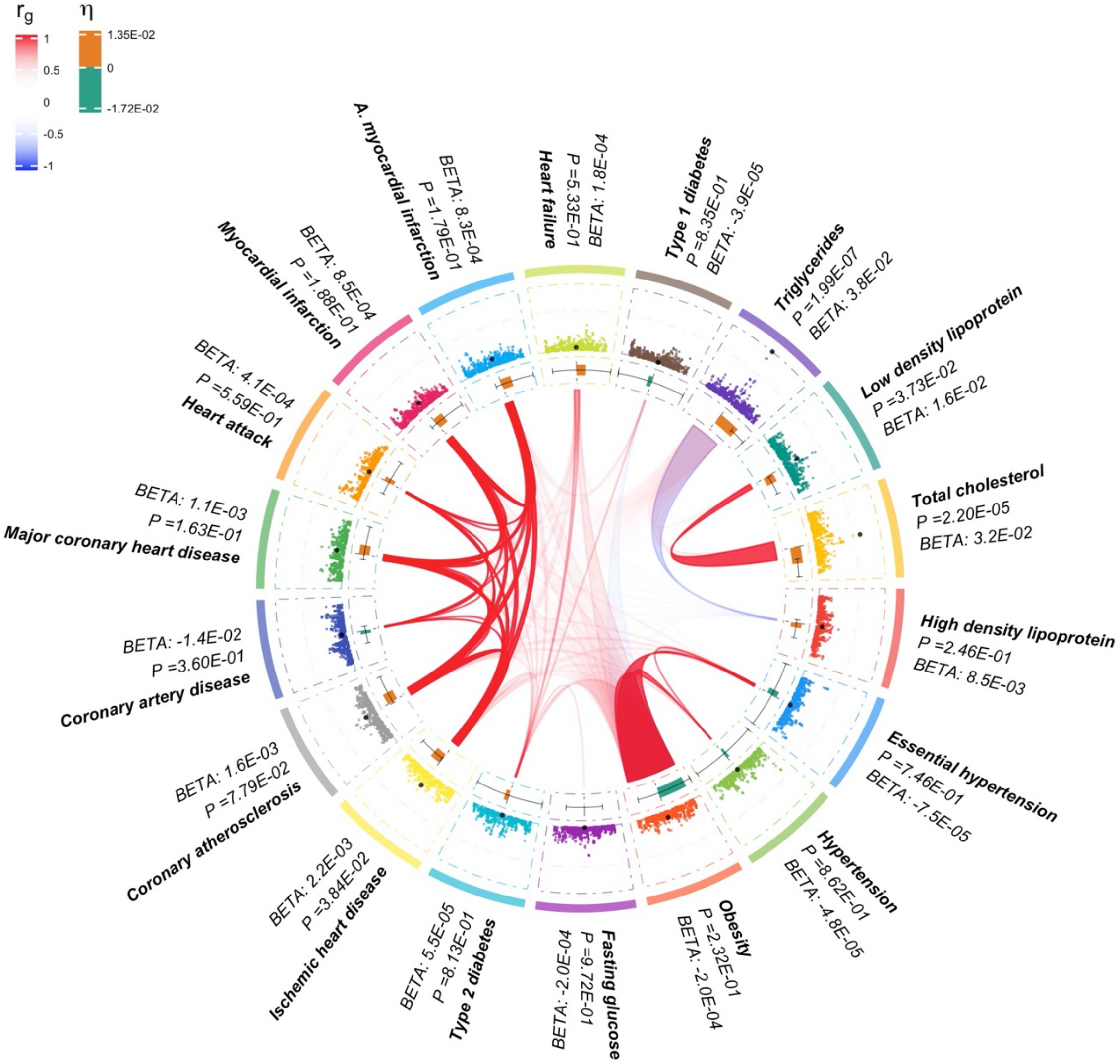

Supplementary Figure 6 K. Pleiotropy plot for rs4823054.

[Group 2: Driven by lipid phenotypes (triglycerides, LDL, HDL, and total cholesterol)]

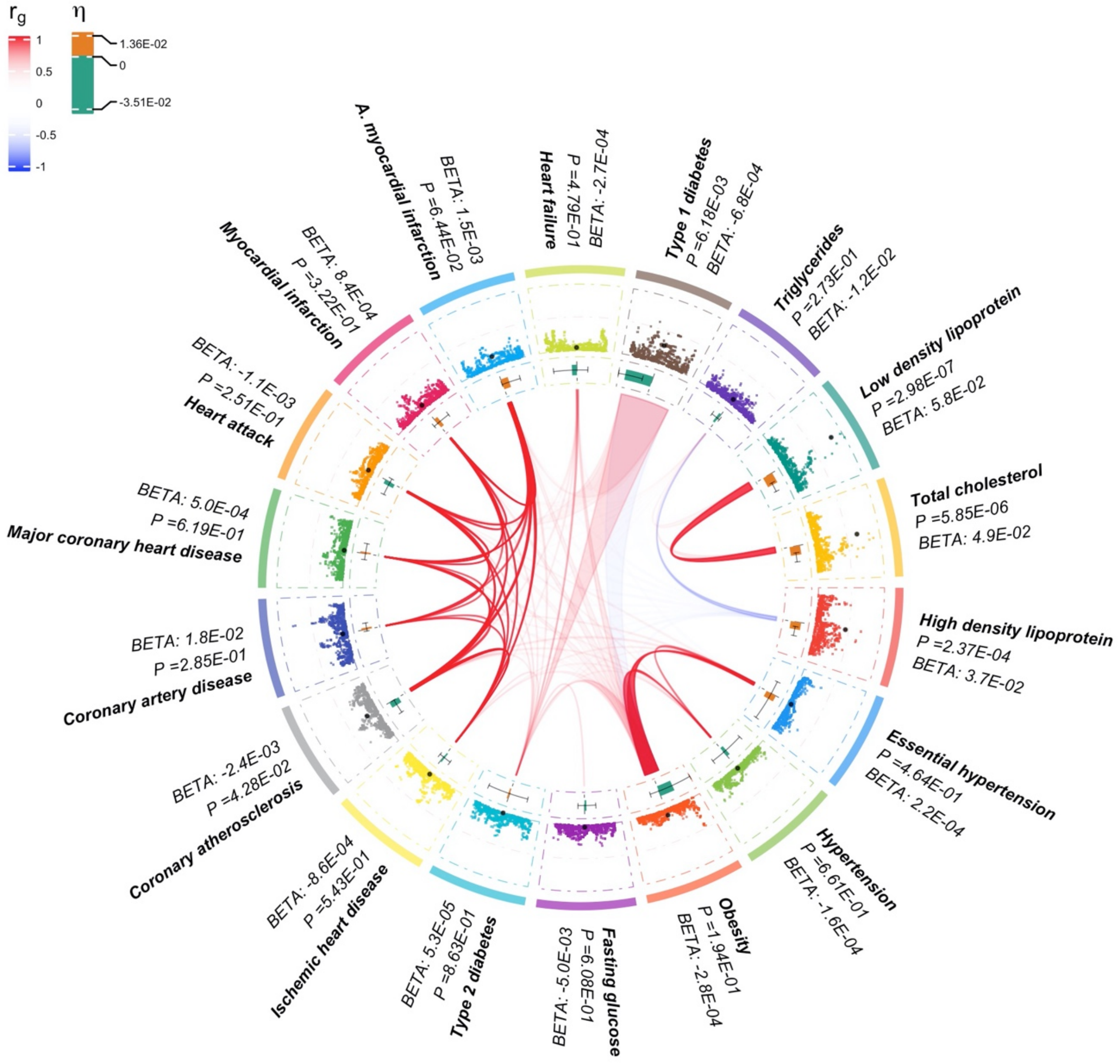

Supplementary Figure 6 L. Pleiotropy plot for rs10026790.

[Group 3: Driven by both coronary artery disease and lipid phenotypes]

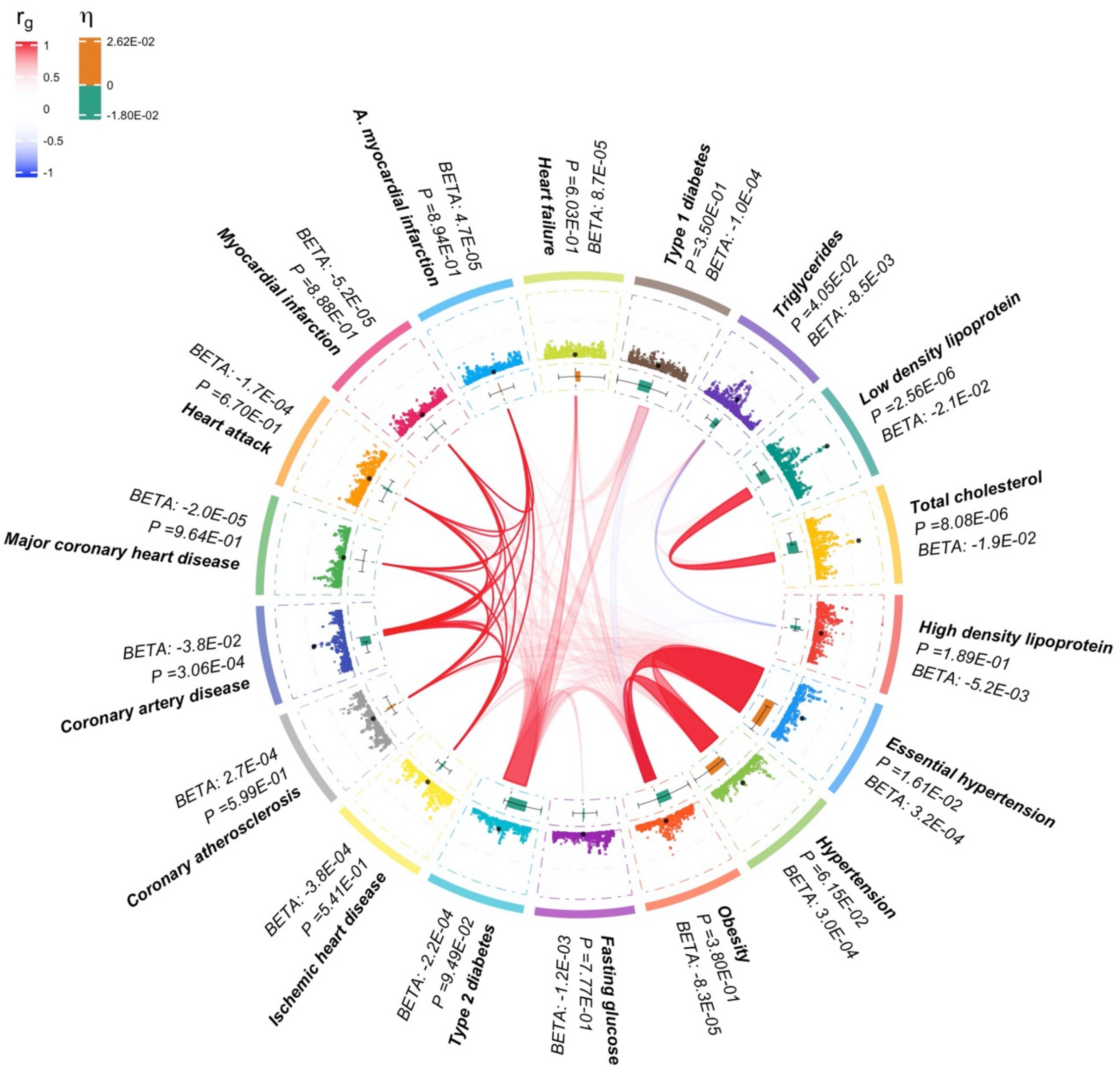

Supplementary Figure 6 M. Pleiotropy plot for rs4074793.

[Group 3: Driven by both coronary artery disease and lipid phenotypes]

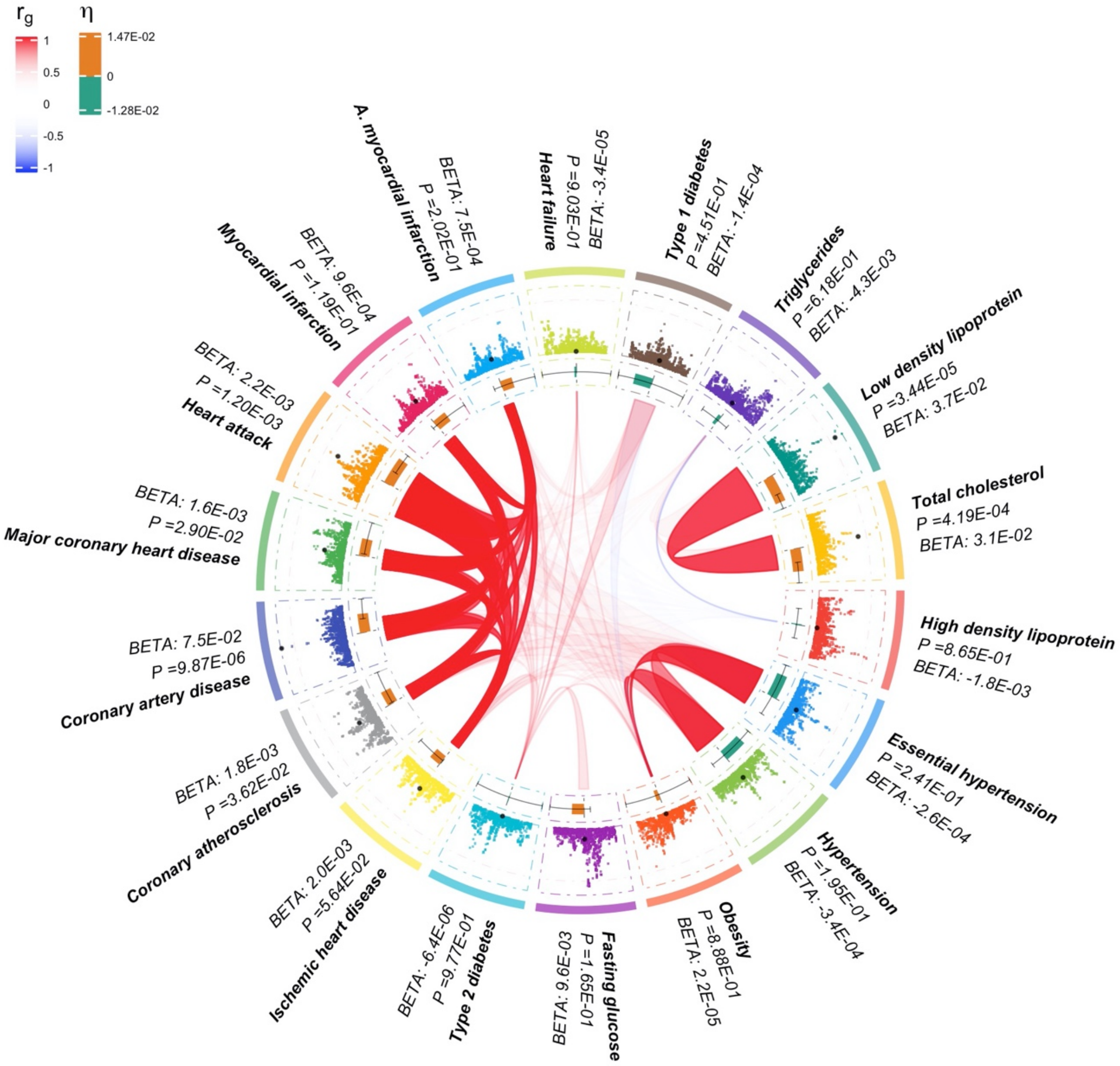

Supplementary Figure 6 N. Pleiotropy plot for rs2237659.

[Group 3: Driven by both coronary artery disease and lipid phenotypes]

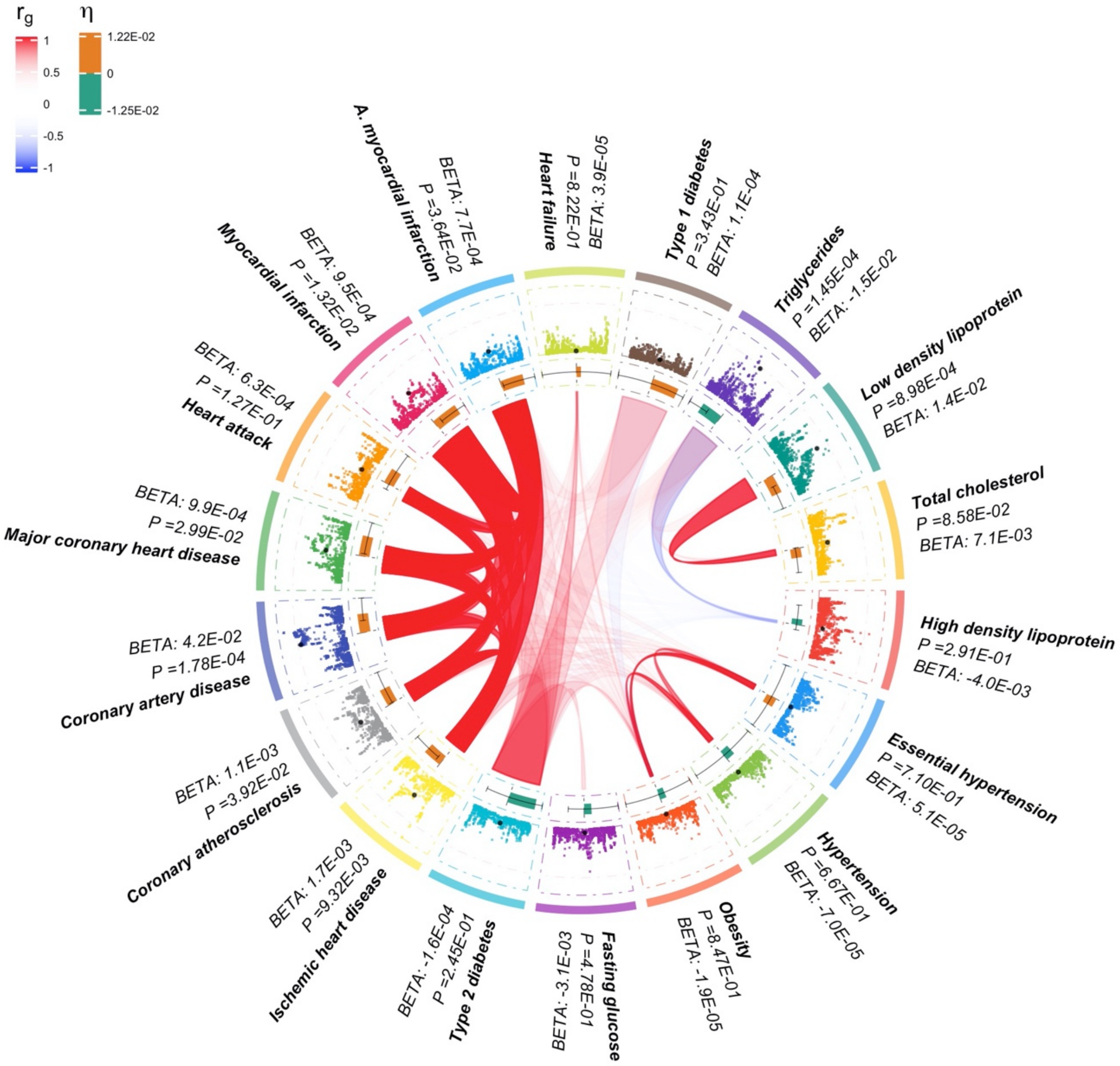

Supplementary Figure 6 O. Pleiotropy plot for rs4393438.

[Group 3: Driven by both coronary artery disease and lipid phenotypes]

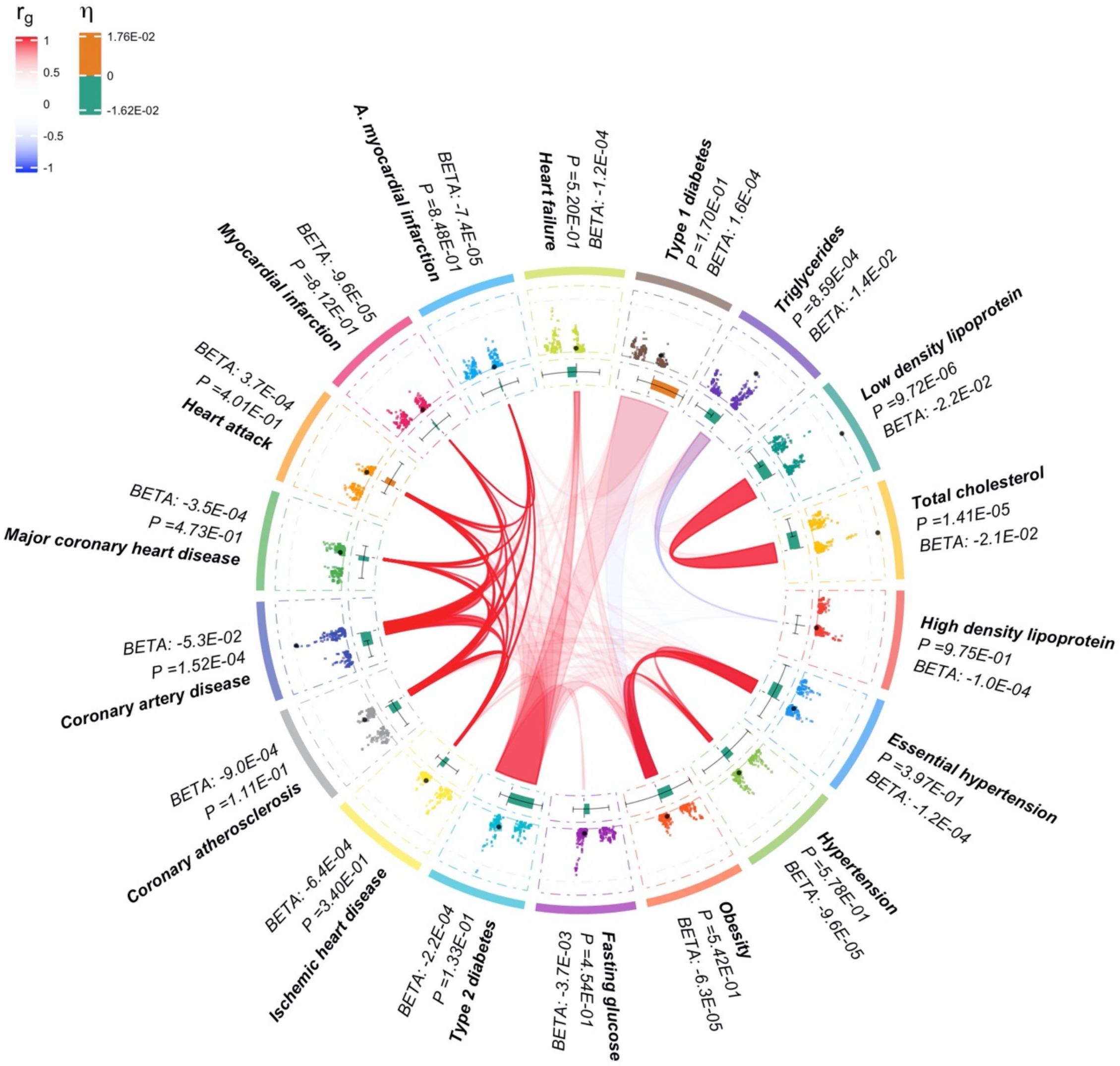

Supplementary Figure 6 P. Pleiotropy plot for rs10733608.

[Group 4: Driven by lipid phenotypes and myocardial infraction]

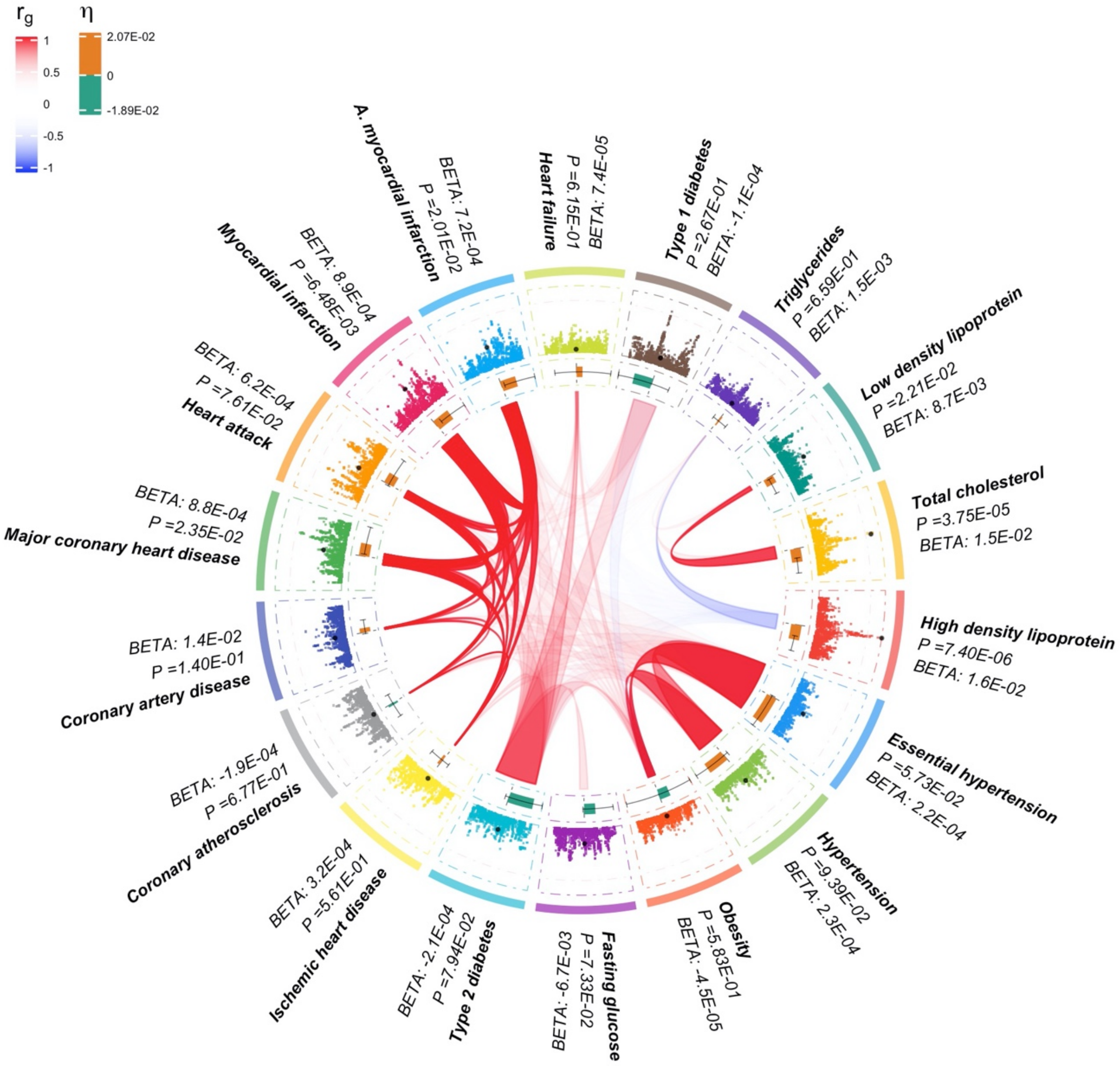

Supplementary Figure 6 Q. Pleiotropy plot for rs12787728.

[Group 4: Driven by lipid phenotypes and myocardial infraction]

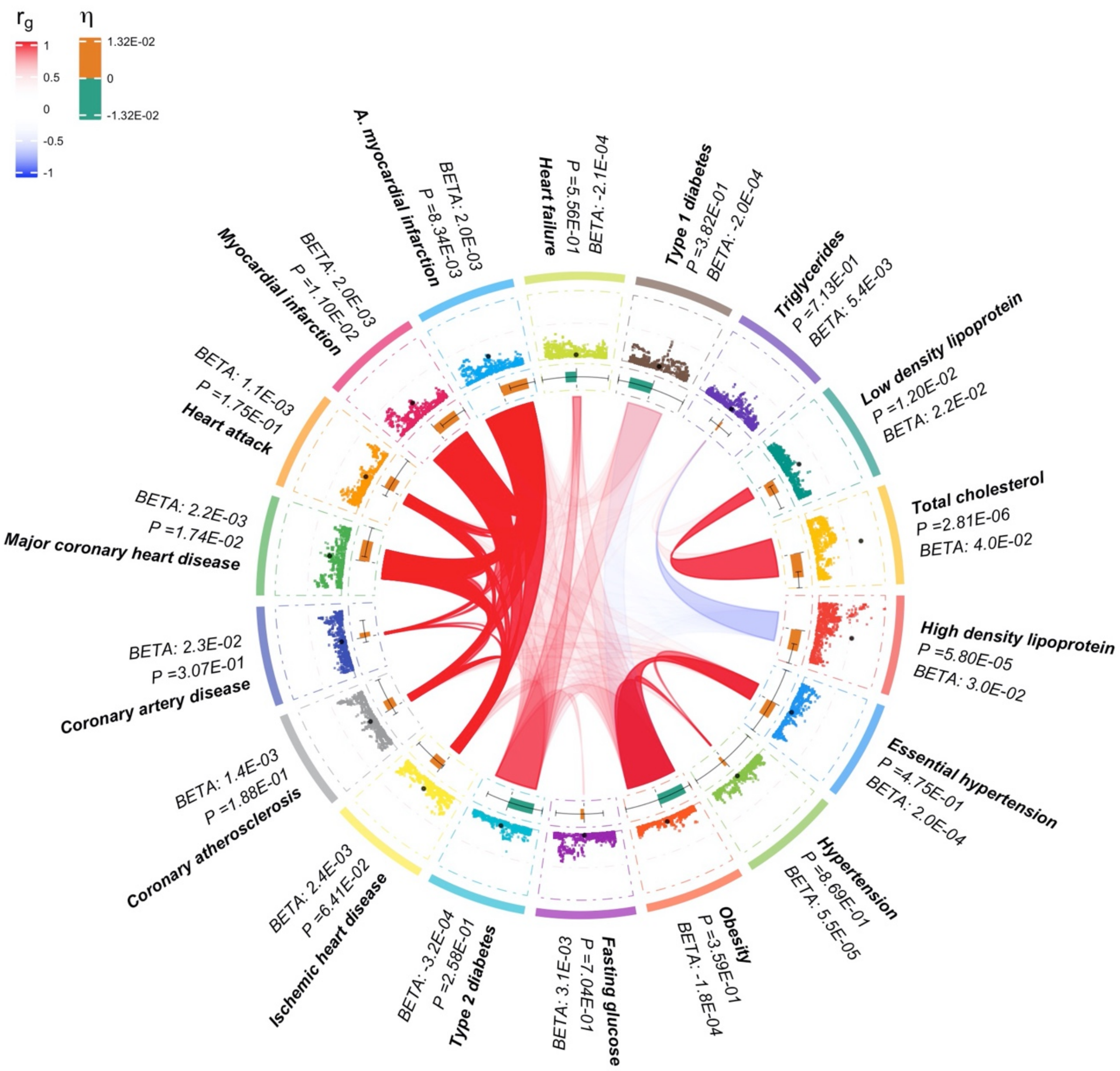

Supplementary Figure 6 R. Pleiotropy plot for rs876320.

[Group 5: Others]

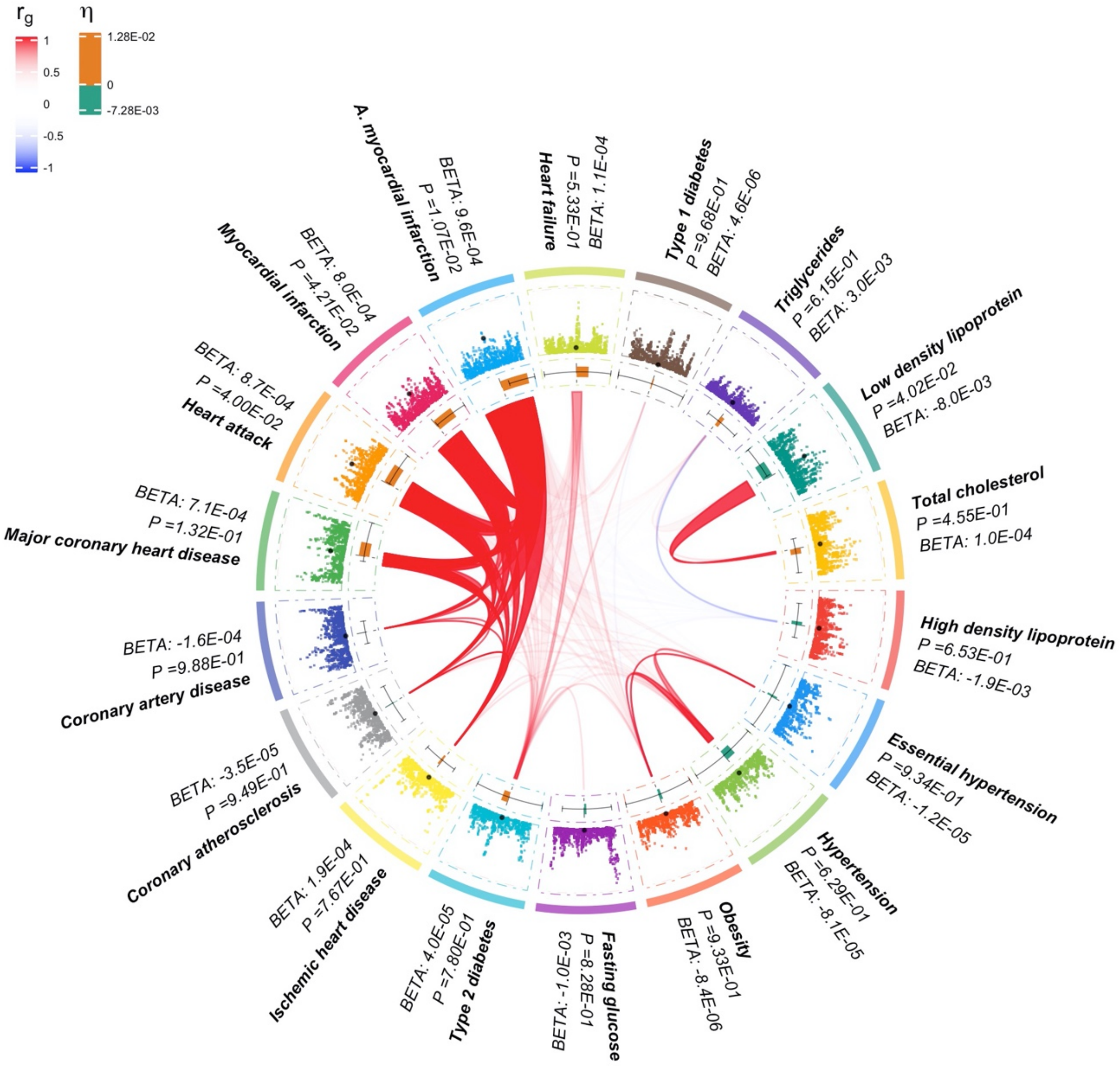

Supplementary Figure 6 S. Pleiotropy plot for rs2891902.

[Group 5: Others]

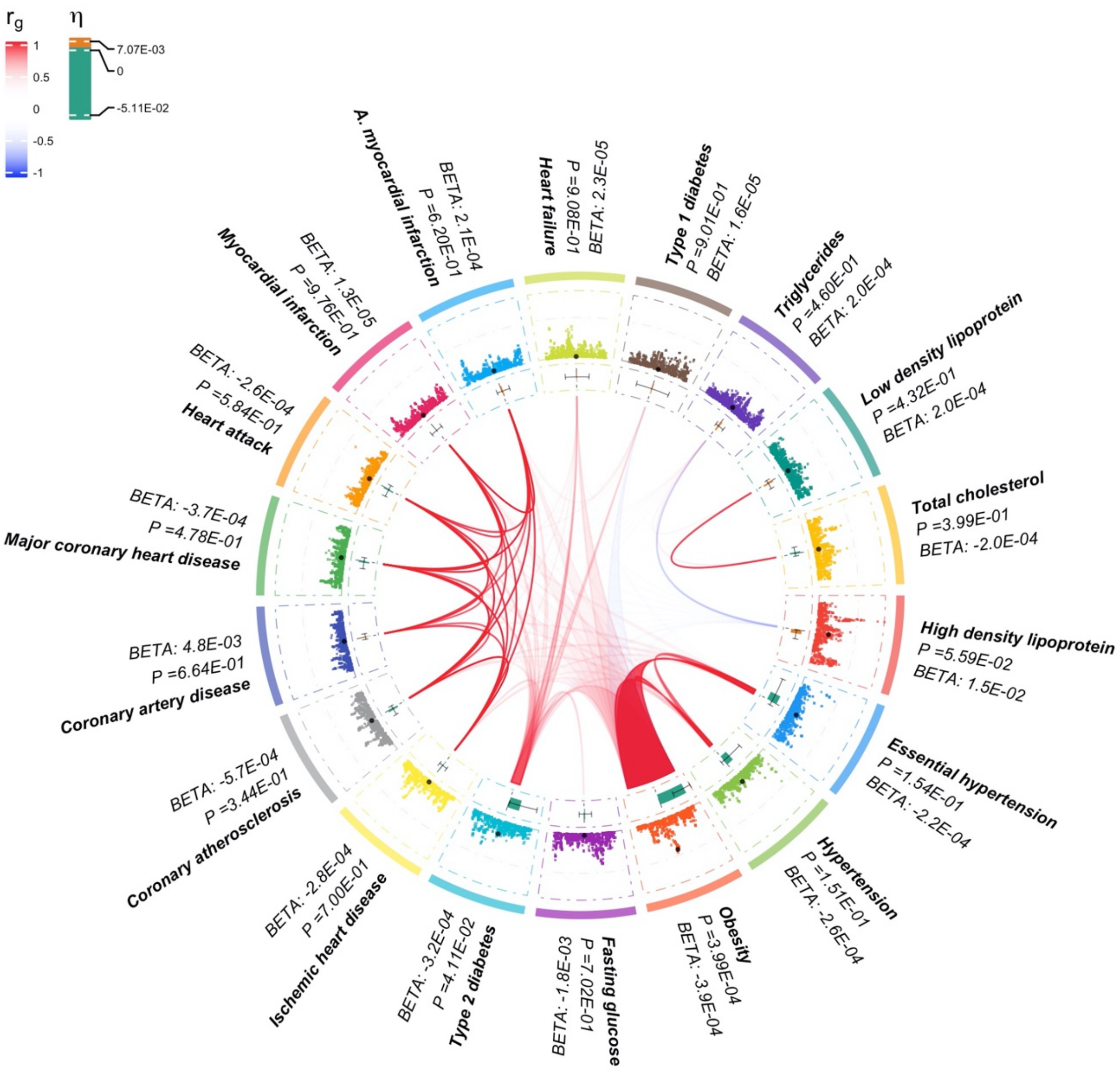

Supplementary Figure 6 T. Pleiotropy plot for rs1039119.

[Group 5: Others]

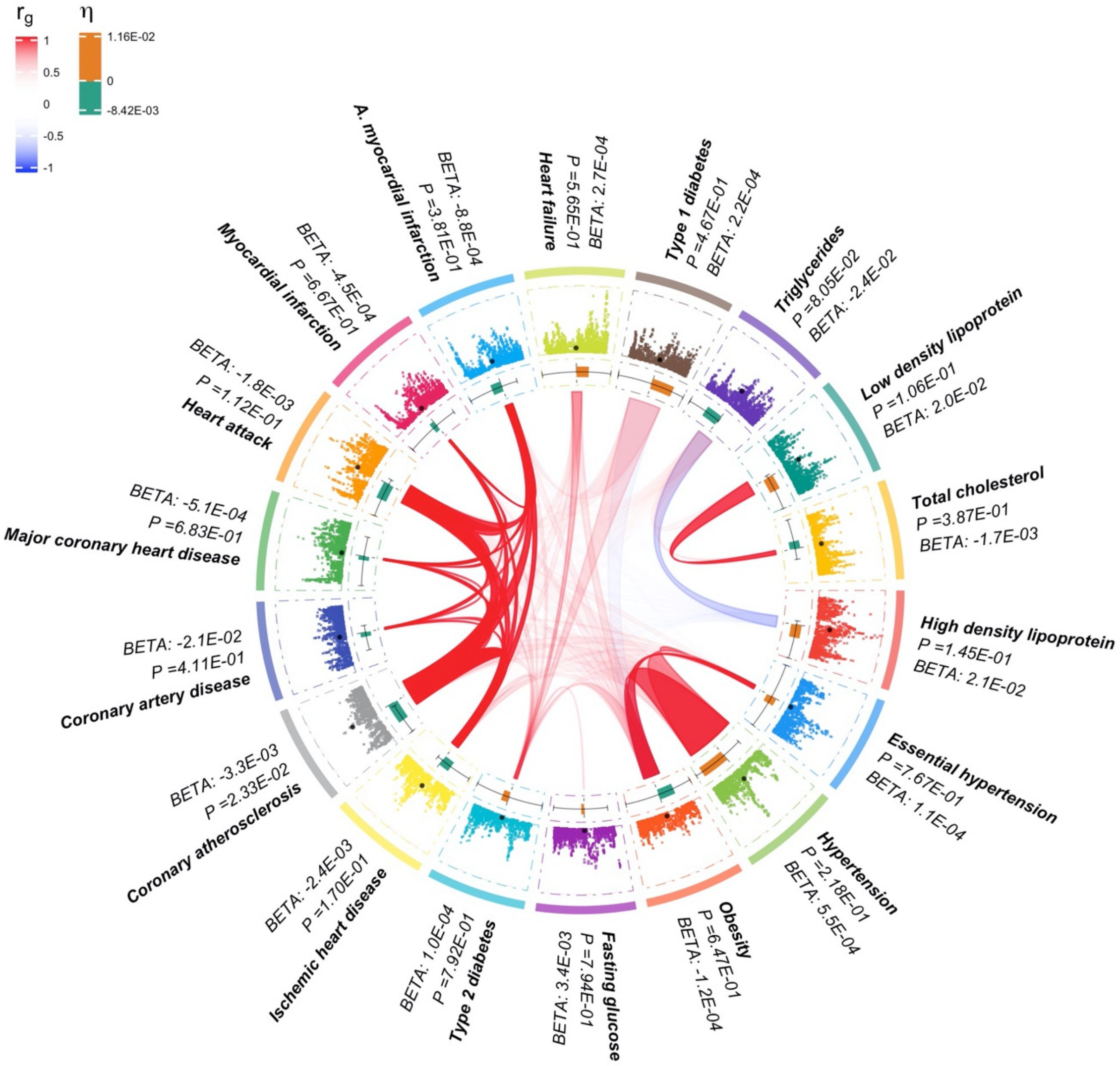

**Supplementary Figure 7.** The computation time of PLEIO for testing 10K SNPs. We measured computation time for combining various numbers of traits from 5 to 200. Dark grey bars are the run times of our optimized version with the Newton-Raphson method. Light grey bars are the run times of the standard implementation using the python optimization function for pseudo Newton-Raphson method.

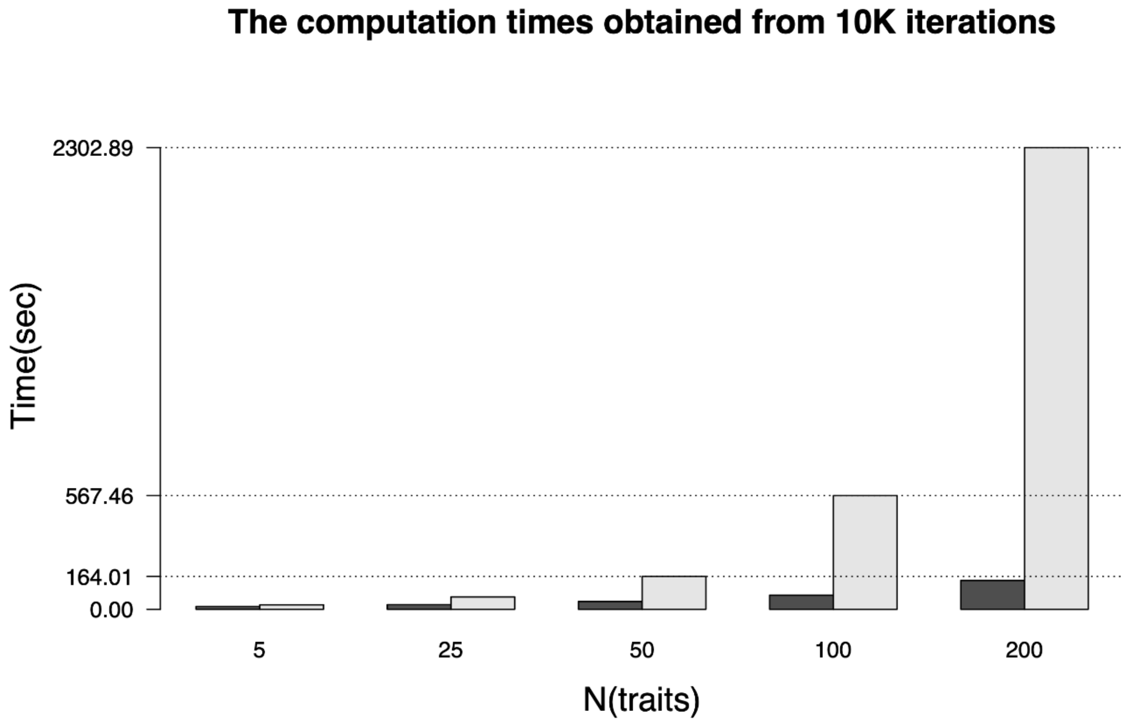

**Supplementary Figure 8.** The computational gain from using the optimized Newton-Raphson method. We measured time to test 10K SNPs using PLEIO, assuming various numbers of traits. Each point indicates the ratio of run times between our optimized Newton-Raphson method (denominator) and the standard python optimization function using pseudo Newton-Raphson method (numerator).

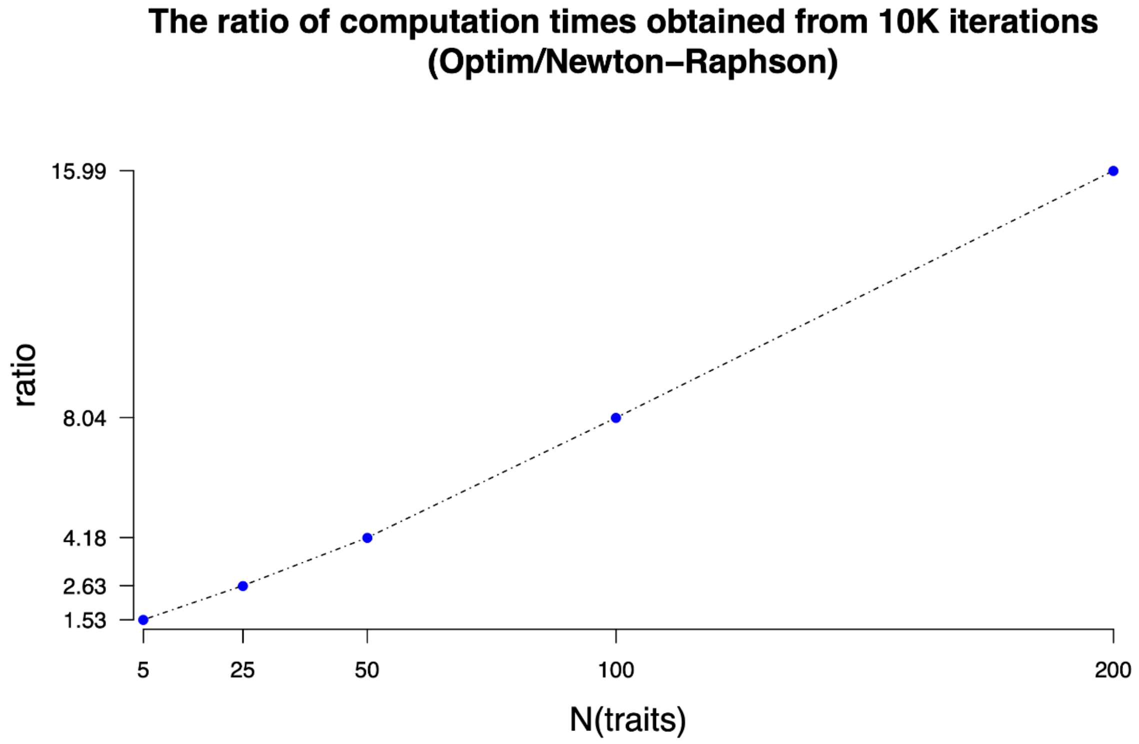

### 2 Supplementary tables

**Supplementary Table 1 PLEIO's FPR in various simulation conditions.** In this simulation, $10^7$ null study sets were generated for each of the 24 situations, and the FPR was calculated at $\alpha = 0.05$. Each test consisted of a unique combination of four parameters: T, $\mathbf{h^2}$, $\boldsymbol{C_g}$, and $\boldsymbol{C_e}$. We changed the number of studies ($T$) to 5, 10, and 20. "Equal $\mathbf{h^2}$" denotes that the heritability is fixed to the value of 0.5, and "Diff $\mathbf{h^2}$" denotes that the values of the heritabilities increase from 0.1 to 0.5. "Uniform $\mathbf{C_g}$" denotes a genetic correlation matrix whose non-diagonal value is fixed to 0.3, and "partitioned $\mathbf{C_g}$" denotes a genetic correlation matrix consisting of two sub-groups where each non-diagonal value within group is fixed to 0.3 and each non-diagonal value between groups is fixed to 0. "No $\mathbf{C_e}$" denotes an environmental correlation matrix whose non-diagonal value is fixed to 0, and "Uniform $\mathbf{C_e}$" denotes an environmental correlation matrix whose non-diagonal value is fixed to 0.5.

| | | $\boldsymbol{T}$ **= 5** | | $\boldsymbol{T}$ **= 10** | | $\boldsymbol{T}$ **= 20** | |
|---|---|---|---|---|---|---|---|
| | | Equal $\mathbf{h_g^2}$ | Diff $\mathbf{h_g^2}$ | Equal $\mathbf{h_g^2}$ | Diff $\mathbf{h_g^2}$ | Equal $\mathbf{h_g^2}$ | Diff $\mathbf{h_g^2}$ |
| **Uniform** $\mathbf{C_g}$ | No $\mathbf{C_e}$ | 0.0499 | 0.0497 | 0.0500 | 0.0497 | 0.0505 | 0.0499 |
| | Uniform $\mathbf{C_e}$ | 0.0499 | 0.0499 | 0.0496 | 0.0499 | 0.0496 | 0.0500 |
| **Partitioned** $\mathbf{C_g}$ | No $\mathbf{C_e}$ | 0.0497 | 0.0498 | 0.0499 | 0.0498 | 0.0509 | 0.0502 |
| | Uniform $\mathbf{C_e}$ | 0.0499 | 0.0500 | 0.0495 | 0.0499 | 0.0505 | 0.0502 |

**Supplementary Table 2 PLEIO's FPR at genome-wide thresholds.** In this simulation, we generated $10^9$ null study sets to test FPR in different $\alpha$ ranging from $5 \times 10^{-2}$ to $5 \times 10^{-8}$. We changed the number of studies ($T$) to 5, 10, and 20 and the fixed $\mathbf{h^2}, \mathbf{C_g}$, and $\mathbf{C_e}$ as follows: For $\mathbf{h^2}$, we used a $T \times 1$ vector whose values increase in the range of (0.1, 0.5). For $\mathbf{C_g}$, we used a genetic correlation matrix consisting of two sub-groups where each non-diagonal value within group is fixed to 0.3 and each non-diagonal value between groups is fixed to 0. For $\mathbf{C_e}$, we used a diagonal matrix.

| **FPR** | **T=5** | **T=10** | **T=20** |
|---|---|---|---|
| $\mathbf{5 \times 10^{-2}}$ | $5.02 \times 10^{-2}$ | $5.03 \times 10^{-2}$ | $5.09 \times 10^{-2}$ |
| $\mathbf{5 \times 10^{-3}}$ | $5.02 \times 10^{-3}$ | $5.05 \times 10^{-3}$ | $5.07 \times 10^{-3}$ |
| $\mathbf{5 \times 10^{-4}}$ | $5.00 \times 10^{-4}$ | $5.07 \times 10^{-4}$ | $5.06 \times 10^{-4}$ |
| $\mathbf{5 \times 10^{-5}}$ | $5.01 \times 10^{-5}$ | $5.04 \times 10^{-5}$ | $4.99 \times 10^{-5}$ |
| $\mathbf{5 \times 10^{-6}}$ | $5.00 \times 10^{-6}$ | $4.91 \times 10^{-6}$ | $5.06 \times 10^{-6}$ |
| $\mathbf{5 \times 10^{-7}}$ | $4.86 \times 10^{-7}$ | $5.94 \times 10^{-7}$ | $5.57 \times 10^{-7}$ |
| $\mathbf{5 \times 10^{-8}}$ | $5.50 \times 10^{-8}$ | $5.70 \times 10^{-8}$ | $4.00 \times 10^{-8}$ |

**Supplementary Table 3**. FPR of PLEIO, ASSET, and RE2C at $\alpha = 0.05$ of no $\boldsymbol{C_e}$ and uniform $\boldsymbol{C_e}$ conditions: We estimated FPR assuming zero environmental correlations among studies and positive environmental correlation whose non-diagonal matrix component is fixed to 0.3. We used $N_{sim}$ = 100,000.

| **Condition** | **DELPY** | **ASSET** | **RE2C** |
|---|---|---|---|
| **no $\boldsymbol{C_e}$** | 0.05088 | 0.04539 | 0.04773 |
| **uniform $\boldsymbol{C_e}$** | 0.05028 | 0.04606 | 0.04751 |
